# A Chemical-Genetic Interaction Matrix Reveals Drug Mechanism and Genetic Architecture

**DOI:** 10.64898/2026.01.28.701823

**Authors:** Jasmin Coulombe-Huntington, Thierry Bertomeu, Caroline Huard, Andrew Chatr-aryamontri, Daniel J. St-Cyr, María Sánchez-Osuna, David Papadopoli, Karine Normandin, Mohammadjavad Paydar, Shannon McLaughlan, Corinne St-Denis, Li Zhang, Henry Say, Roger Palou, Chris Stark, Bobby-Joe Breitkreutz, Almer M. van der Sloot, Sandhya Manohar, Hugo Lavoie, Katherine L. B. Borden, Brian Raught, Damien D’Amours, Frank Sicheri, Alain Verreault, Sylvie Mader, Sylvain Meloche, Marc Therrien, Pierre Thibault, Brian Wilhelm, Peter B. Dirks, John D. Aitchison, Elizabeth Patton, Randall W. King, Philippe P. Roux, Guy Sauvageau, Trang Hoang, Anne Marinier, Lea Harrington, Benjamin Kwok, Vincent Archambault, Ivan Topisirovic, Mike Tyers

## Abstract

To probe drug mechanism of action (MOA) and interrogate the genetic architecture of human cells, we carried out isogenic genome-wide CRISPR/Cas9 knockout screens against 310 diverse drugs, bioactive compounds, and stress conditions. Stringent statistical correction for gene knockout fitness defects yielded a large-scale matrix of >12,000 high confidence chemical-genetic interactions (CGIs). This dataset revealed many previously unappreciated off-target effects for well-characterized compounds and novel MOAs for uncharacterized compounds. The CGI matrix uncovered dense genetic modules that yielded new biological insights into phospholipidosis, mitotic regulation, metabolism, the DNA damage response, and mTOR signaling. The dataset allowed identification of multi-drug sensitization and resistance mechanisms, inference of gene function, elaboration of cross-process connectivity, evaluation of the cell type specificity of CGIs, prediction of chemical synergism, and extensive annotation of understudied genes. This resource provides a map of the genetic landscape in human cells and a framework to help guide drug discovery.

## Introduction

Genetic interactions, in which two or more co-occurring mutations alter a phenotype in a non-linear fashion, reflect the functional organization of cells and organisms.^1^ Systematic genetic screens with genome-wide collections of mutant alleles, first carried out in model organisms, uncovered a vast hierarchical genetic network that controls cellular function.^2^ Analysis of pairwise genetic interactions in budding yeast provided the first global view of the functional hierarchy in eukaryotic cells.^3^ With the advent of efficient and precise gene disruption methods based on single guide RNA (sgRNA)-directed nucleases of bacterial CRISPR systems, massive parallelization of genetic perturbation experiments in human cell lines has become possible using programmed genome-wide sgRNA libraries.^4^ To date, thousands of CRISPR screens in human cells have uncovered the fitness effects of loss-of-function mutations in different cell types and/or under different conditions.^5^ These studies have identified cancer cell line-specific vulnerabilities^6^ and helped identify the function of many uncharacterized genes.^7-12^ However, unlike in isogenic model organism systems, which enable studies in uniform genetic backgrounds, the integration of diverse CRISPR screens to build a unified genetic network is complicated by the extreme genetic variability of cancer cell lines, tissue of origin effects, and inter-laboratory reproducibility.^13,14^ Furthermore, the direct mapping of human genetic interactions by CRISPR-based methods requires the time-consuming generation and screening of cell lines that bear specific mutations of interest.^15^

A drug or other bioactive agent can perturb the cellular regulatory network in a manner analogous to a genetic mutation.^16^ Advantages of chemical perturbations are that small molecules can rapidly modulate cellular state, avoid long-term adaptive mutations in response to genetic modifications, and be fine-tuned by concentration.^17^ The systematic exploration of chemical-gene interactions (CGIs) at genome-wide scale, termed chemogenomics, can uncover compound mechanism of action (MOA) and gene function.^18-20^ By analogy to genetic interactions, CGIs may be classified as either synthetic sick/synthetic lethal (SSL, also called enhancing or negative interactions), which increase sensitivity to a compound, or rescue (also called suppressing or positive interactions) which increase resistance to a compound.^19^ The amalgamation of many chemogenomic screens results in a network of CGIs that accurately reflects genetic interaction network architecture.^20^

Recent CRISPR-based chemogenomic studies in human cells have demonstrated the power of genetic profiles to interrogate compound MOA and biological mechanisms but have typically focused on single or limited sets of compounds.^5,21-23^ Here, we report a largescale dataset comprising over 400 genome-wide CRISPR screens against more than 300 bioactive compounds in an isogenic cell line context. We developed a scoring algorithm called CRANKS to normalize for gene knockout fitness defects and generated a network of >12,000 high confidence CGIs. Gene-gene correlations across the CGI matrix unveiled new features of cellular genetic architecture and unraveled numerous new gene functions, modular hierarchies, and inter-process cross-connections. Chemogenomic profiles identified numerous hitherto unknown off-target effects, new MOAs, common patterns of genetic sensitization and resistance, and chemical synergism. This chemogenomic resource further highlights the value of genome-scale CRISPR screens in understanding the functional wiring of human cells and the mechanisms whereby bioactive small molecules alter cellular function.

## Results

### Strategy for chemogenomic screens in human cells

We carried out 428 isogenic screens in the NALM-6 human pre-B lymphocytic cell line using the Extended Knockout (EKO) sgRNA library that targets all annotated human genes, hypothetical genes and splice isoforms.^24^ The pseudo-diploid NALM-6 line is amenable to parallelized CRISPR screens as it grows in suspension and exhibits high efficiency Cas9-mediated indel formation. To determine suitable compound concentrations for screens, we estimated IC30 values, i.e., the concentration that causes 30% inhibition of proliferation compared to solvent control, in 384 well format dose-response curves. The IC30 allowed detection of both genetic rescue (i.e., enhancement of proliferation/survival or positive selection) and synthetic sickness/lethality (SSL, i.e., suppression of proliferation/survival or negative selection). The EKO sgRNA library was introduced into a doxycycline-inducible Cas9 clone^24^ and the uninduced pool was frozen in large aliquots that allowed the same isogenic population to be used for each of 9 different screening rounds with minimal loss of guides that target essential genes. For each screening round, the pool was expanded and Cas9 expression was induced with doxycycline for 7 days, after which the pool was split into separate flasks (Figure 1A). Each round included untreated and vehicle solvent control screens. This isogenic pool approach yielded highly comparable results between screen rounds for compound replicates (see below). We screened 305 unique compounds and 5 conditions that targeted a wide array of biological processes grouped in 10 general categories: regulation of gene expression, cell cycle, infectious disease, signal transduction, organelle function, metabolism, proteostasis, DNA damage/repair, environmental stress and uncharacterized MOA (Figure 1B; Tables S1A and S1B). All compounds were verified by analytical chemistry methods (Table S1C; see Methods).

**Figure 1.**
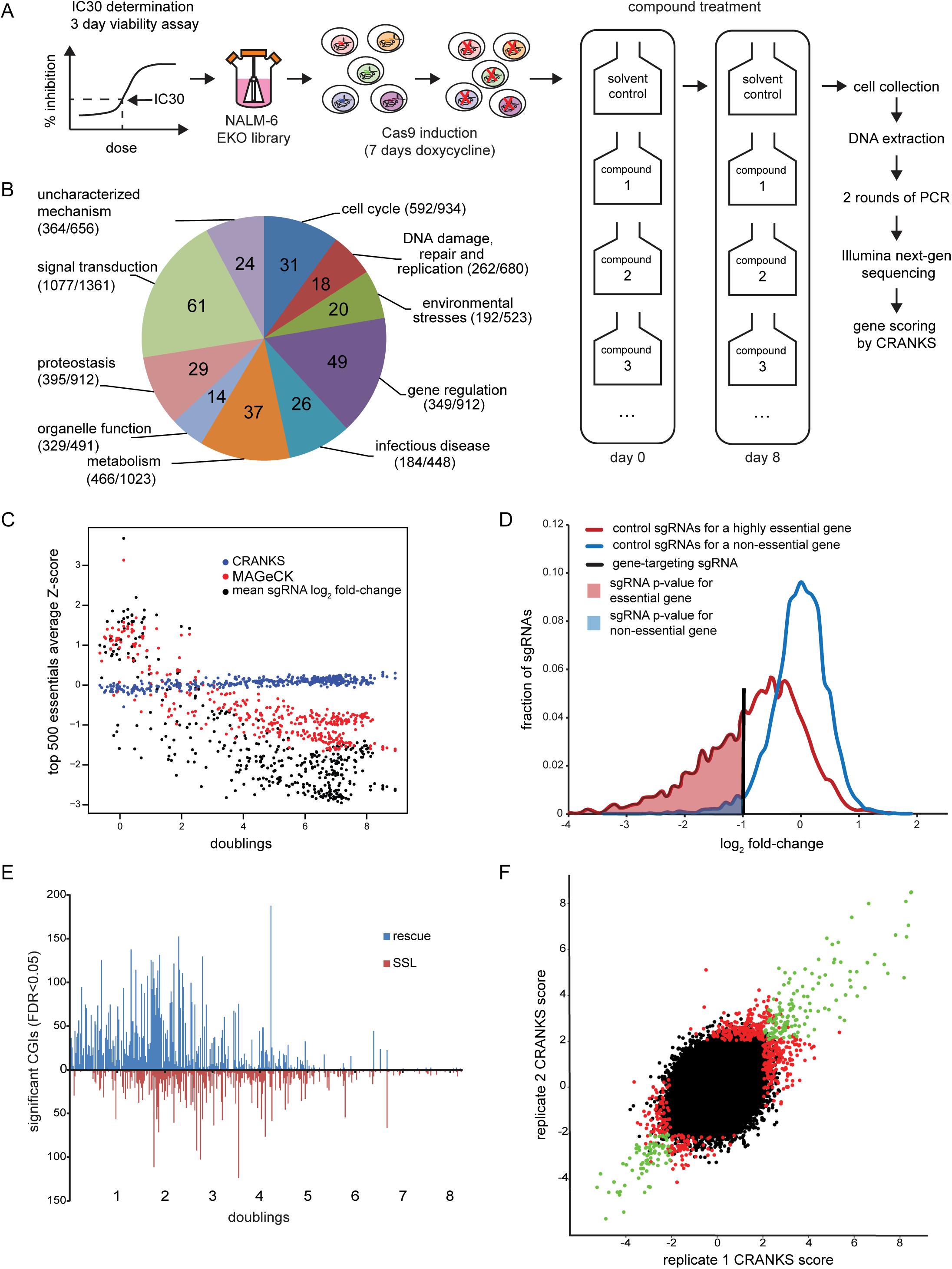
Chemogenomics workflow and dataset overview. (A) Schematic of CRISPR chemical screen pipeline in NALM-6 pre-B cell line. (B) Categories of compounds screened. Total number of SSL and rescue genetic interactions, respectively, for each category indicated in parentheses (see Table S1B). (C) Average z-score of the top 500 essential genes by CRANKS, versus mean sgRNA log_2_ fold-change and the MAGeCK scoring algorithm as a function of the number of population doublings. (D) CRANKS calculation of essential gene sgRNA p-values by comparison to sgRNA fold-changes of genes with similar essentiality. (E) Relationship between the number of CGIs and proliferation per screen. (F) Screen reproducibility for aggregate of 9 different compounds screened in different cycles. Genes shown in green scored as significant in both replicates and those in only one replicate in red. See also Figure S1.

### Screen scoring and essential gene bias correction

Although each screen was run at the estimated IC30, the final number of population doublings in the screens was variable due to culture scale-up, compound half-life and/or sharp dose-response curves that precluded accurate IC30 estimation. These proliferative differences between screens caused variable depletion of sgRNAs that target essential genes as a function of the number of cell divisions rather than time, i.e., compounds that inhibited proliferation caused an apparent enrichment of essential genes compared to control treatments. The average sgRNA log_2_ fold-changes or MAGeCK scores^25^ for the top 500 essential genes were enriched in screens with few population doublings even though all screens lasted 8 days (Figure 1C). To overcome this confounding enrichment effect, we developed a condition-specific RANKS algorithm (CRANKS), that estimates sgRNA-specific p-values by comparing frequency fold-change to a set of ∼5,000 sgRNAs targeting comparably essential genes, as opposed to RANKS,^24^ which uses a fixed control set of sgRNAs (Figure 1D). CRANKS effectively removed the proliferation-driven bias in essential gene scores (Figure 1C). Unlike existing CRISPR screen scoring algorithms, such as log_2_ fold-change^11^, MAGeCK^25^, BAGEL^7^ or DrugZ,^26^ CRANKS controls for gene essentiality without the need for a proliferation-matched control screen. This scoring approach also allowed us to pool control screen read counts from multiple time points, thereby providing a robust estimation of background sgRNA frequencies and reduced batch variation between screen rounds. Across the dataset, 95% of screens with fewer than 6 population doublings produced one or more significant hits (FDR < 0.05) versus 59% of screens with weaker proliferation inhibition (p=1.5e-27, Fisher’s exact test). Screens with extreme proliferation inhibition (<1 doubling) yielded rescue interactions in 95% of cases but generated SSL interactions in only 56% of cases, compared to 94% of screens with 1 to 6 doublings (p=1.1e-9). These results established an empirical “Goldilocks zone” of between 1 and 6 population doublings for optimal discovery of chemical-genetic interactions in single screens (Figure 1E).

To assess screen reproducibility, we replicated screens for 9 compounds in different rounds at the same or a similar dose. We observed strong reproducibility, especially for the highest scores (Figure 1F). The scores were well correlated overall (R=0.68), consistent with a greater lack of sensitivity (false negatives) than specificity (false positives). These results also suggest that many sub-threshold gene scores (FDR>0.05) are likely relevant. Scores were robust to the number of sgRNA read counts (Figure S1A) and number of cells per guide used during screening (Figure S1B). Non-expressed genes (log_2_ reads per million < 3) rarely produced significant CGIs (Figures S1C and S1D; Table S1D), consistent with a low rate of false positive interactions. In total, the dataset contained 12,150 significant CGIs for 3178 genes, including 4210 SSLs, 7940 rescues and 11,480 unique gene-compound/condition interactions (FDR<0.05, Figure 1B; Table S1E). A gene-by-compound heatmap of this CGI matrix self-organized into many functional clusters (Figure S1E; Table S2), a number of which were investigated in detail below.

### The CGI matrix is highly enriched in known target genes

In chemical-genetic screens, detrimental compound effects are alleviated by the loss of genetic effectors of compound action and result in increased clonal proliferation (we refer to genes in this category as rescues). Conversely, loss of genetic inhibitors enhances compound action and result in decreased proliferation (we refer to genes in this category as SSL or sensitizers). Presumptive indels that scored in screens typically represent loss-of-function alleles, although hypermorphic or neomorphic alleles may be generated in some circumstances. For brevity of discussion, hit gene names are taken to imply loss-of-function indel mutations unless indicated otherwise.

Compounds with a well-established MOA served as benchmarks for biological relevance of the dataset. We frequently observed known target effectors and inhibitors as top-scoring genes. To assess how often direct drug targets scored as a top hit, we compiled genes encoding known targets from the Drug Repurposing Hub^27^ and/or curation of the primary literature (Table S3). For 89 of the most productive screens (≥10 hits) with unique compounds, where at least one reported target was well-expressed in NALM-6 (log_2_ RPM ≥ 5.5), known direct target genes were the top hit in 16% of screens, in the top 10 hits in 29% of screens and in the top 100 hits in 47% of screens (Figure 2A). The interpretation of compound-target gene interactions is subject to biological context, as illustrated in the examples below.

**Figure 2.**
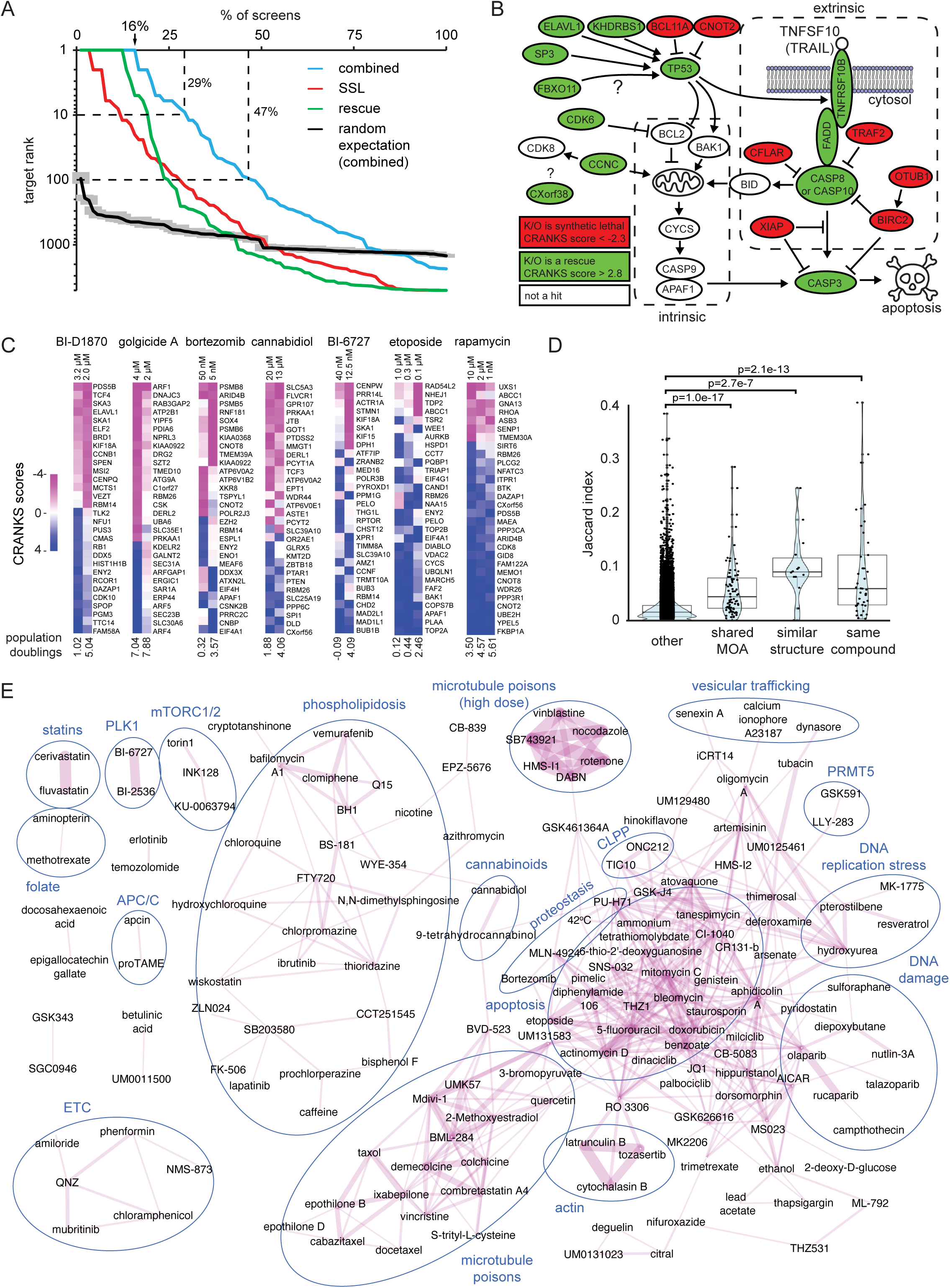
Prediction of compound mechanism of action. (A) Enrichment of known compound targets from the literature within top-scoring well-expressed genes for 89 distinct compounds. For compounds with multiple reported targets, only the top ranked target was used. Random expectation shows the median lowest ranks from 100 random gene orders for each screen with confidence intervals (grey) corresponding to the bootstrapped standard error with 100 resamplings. (B) TRAIL ligand screens uncover the extrinsic apoptotic network. Two independent TRAIL screens carried out at 5ng/mL and 50 ng/mL were merged (see Table S4). Significant hits are overlaid on the core apoptosis network. (C) Effect of compound dose on screen hit profiles. Scores of top 15 rescue and SSL hits are shown for 7 different compounds. Additional SSLs or rescues were included when there were fewer than 15 significant hits (FDR<0.05) of the opposite type. (D) Compounds that share common targets (shared MOA), structurally similar compounds, and screens of the same compound had more similar chemogenomic profiles than unrelated compound pairs. For compounds screened multiple times, the most similar screen pairs are shown. (E) Compound screen similarity network. Edges represent overlapping CGI profiles (Jaccard index ≥ 0.1). Compounds screened multiple times are represented by the screen with the most hits. Cellular processes known or inferred to be targeted by compounds in selected clusters are indicated. APC/C, anaphase promoting complex/cyclosome; ETC, electron transport chain. See also Figures S2-S5.

### Delineation of compound MOA by genetic rescue and sensitization

Many screens yielded insights into specific compound MOAs and biological response networks. The TNF-related apoptosis-inducing ligand (TRAIL) triggers the extrinsic apoptotic pathway by binding to the TNFSF10B receptor and subsequently activating caspase 8/10.^28,29^ We combined results from two TRAIL screens run at different doses that yielded similar profiles (Figure 2B; Table S4). Top rescues included the receptor TNFSF10B, the adaptor protein FADD and the effector caspases CASP8 and CASP10, while top sensitizers included known inhibitors of the pathway (CFLAR, XIAP, TRAF2, BIRC2). Mediators of intrinsic apoptosis (BID, CYCS, CASP9, APAF1) did not have significant scores. We selected 10 hits for validation with two different sgRNAs in TRAIL dose-response (D-R) curves, all of which confirmed the screen hits (Figure S2A). We assessed lineage and cell line specificity of TRAIL-gene interactions in the Jurkat T cell lymphocytic line. Of 10 hit genes tested, 4 modulated sensitivity to TRAIL (TNFRSF10B, FADD, CASP8, TRAF2) while 6 had no appreciable effect (CNOT2, KHDRBS1, CXorf38, CASP3, FBXO11, ZNF217) (Figure S2B). Notably, NALM-6 cells are wild type for the tumor suppressor TP53 whereas Jurkat cells are mutated for TP53^30^, and the 6 genes that failed to modulate TRAIL sensitivity in Jurkat cells are linked to TP53 activity.^31-34^. For example, FBX011 mediates p53 neddylation^33^ and loss of FBXO11 in mice causes similar developmental defects as loss of TP53.^35^ Consistently, FBXO11 is highly correlated with TP53 across our dataset (R=0.52).

Direct chemical activators of target enzymes produced strong rescue signatures. The phorbol ester PMA, which activates protein kinase C (PKC), was rescued by PKC isoforms (PRKCD and PRKCE) and modulated by many known RAS/MAPK genes downstream of PKC, as well as by previously unreported genes, such as TM2D2 (Figure S3A). Similarly, the imipridones ONC201 and ONC212, which activate the mitochondrial CLPP protease, were rescued by CLPP and its obligate mitochondrial processing protease MIPEP.^36^ Intriguingly, the EIF2AK4 kinase (a.k.a. GCN2) and its activator GCN1L1, which inhibit protein synthesis upon nutrient limitation and other types of stress,^37^ strongly rescued the EGFR inhibitor erlotinib and two other unrelated kinase inhibitors, the WEE1 inhibitor MK-1775 and the DYRK3 inhibitor GSK626616, as well as the DNA damaging agent temozolomide, which has a purine-like ring structure (Figures S3B-S3D). None of these compounds share known biological targets but recent work suggests that erlotinib, neratinib and sunitinib activate EIF2AK4 as an off-target effect.^38,39^ These results suggest that MK-1775, GSK626616 and temozolomide may also adventitiously activate EIF2AK4, with potential impacts on drug safety and/or efficacy.

Compounds that inhibited either positive or negative steps in biological pathways also elicited strong rescue phenotypes. Two GSK3 inhibitors, LY2090314 and CHIR-99021, were sensitized by GSK3B itself and rescued by the downstream Wnt pathway transcription factors CTNNB1 (β-catenin) and LEF1 (Figure S3E). Top rescues of the CDK4/6 kinase inhibitor palbociclib included RB1, the repressor of G1/S transcription that is the primary target of CDK4/6, and the ubiquitin ligase subunit AMBRA1 that mediates degradation of D-type cyclins (CCNDs), the obligate activators of CDK4/6^40^ (Figure S4A). Counterintuitively, CCND3 was also a top rescue of palbociclib. This rescue effect was directly proportional to the relative 3’-to-5’ position of each sgRNA (R=0.89, p=6.5e-4, Figure S4B), consistent with the generation of hypermorphic C-terminally-truncated cyclin D3 protein variants resistant to degradation.^41^

Gene knockouts that disrupted toxic drug complexes formed one class of strong rescues. For example, the poly(ADP-ribose) polymerase (PARP) inhibitors rucaparib and olaparib were rescued by PARP1 (Figure S4C; see DNA damage section below). Other toxicity rescue mechanisms included the reduced uptake and/or metabolism of toxic agents, exemplified by the rescue of arsenate by the known phosphate transporters XPR1 and SLC20A1, as well as other phosphate metabolism genes (Figure S4D). This result suggests that arsenate toxicity primarily arises from interference with phosphate metabolism.^42^ Finally, rescue profiles also illuminated unknown mechanisms. For example, sub-optimal proliferation caused by a poor batch of fetal bovine serum (FBS) was rescued by the deoxycytidine kinase DCK, the nucleoside transporter SLC29A1 and the dCMP deaminase DCTD, suggesting the presence of a toxic cytosine nucleoside analog as a contaminant in the FBS. To test this hypothesis, we screened the nucleoside analog cytarabine and observed a similar rescue profile (Figure S4E).

Other examples of expected compound sensitization in the dataset include SSL interactions of proteasome subunits (e.g. PSMB5, PSMB6, PSMB8) with the proteasome inhibitor bortezomib (Figure S4F) and GOT1 sensitization to the mitochondrial complex I inhibitors phenformin and rotenone through effects on aspartate metabolism (Figure S4G).^43^ In a more complex example, the IMPDH inhibitor and anti-viral ribavirin^44^ was moderately sensitized by the redundant paralogs IMPDH1 and IMPDH2 (CRANKS scores -1.31 and -1.43, respectively) while the upstream enzyme AMPD2 strongly sensitized to ribavirin, likely due to depletion of the AMP pool acted on by IMPDH (Figure S4H). Other SSL interactions highlighted potential new mechanisms. For example, ADK sensitized cells to hypoxia (5% O_2_), potentially by limiting AMP production and attenuating cellular energy balance (Figure S5A). Consistently, ADK sensitized cells to the ETC inhibitors phenformin, rotenone and CCCP (Figure S4G), as well as to compounds that affect energy consumption, such as the mitochondrial protein synthesis inhibitor chloramphenicol, and the GSK3β inhibitor LY2090314 (Figure S3E). ADK strongly rescued the effects of two intermediates in purine biosynthesis, cAMP and AICAR (Figures S5B and S5C). AICAR is thought to activate the AMPK kinase that senses cellular energy balance and restricts cellular energy expenditure^45^. However, ADK did not rescue another AMPK activator PF-06409577, despite its strong rescue by AMPK subunits (Figure S5D). These results show that multiple inhibitors of the same target pathway can yield distinct CGI profiles. A final noteworthy sensitization mechanism is the generation of heterozygous deletions or hypomorphic alleles of highly essential target genes. For example, WEE1 (756^th^ most essential), was the 7^th^ strongest SSL for the WEE1 inhibitor MK-1775 (Figure S3C) and RRM2 (891^st^ most essential) was the 7^th^ strongest SSL for the ribonucleotide reductase inhibitor hydroxyurea (Figure S5E).

### Concentration effects on CGI profiles

We found that different doses of the same compound can yield divergent profiles due to bias towards SSL or rescue hits, dose-dependent secondary effects, or compensatory mechanisms (Figure 2C). As expected, higher doses generally favored rescue interactions while lower doses favored SSL interactions, as for the ribosomal protein S6 kinase (S6K) inhibitor BI-D1870. Lower concentrations often generated weaker but still well-correlated signatures, as for the allosteric mTOR inhibitor rapamycin and the vesicle trafficking inhibitor golgicide A. In other cases, a subset of genes exhibited unexpected differences, e.g., for the cannabinoid receptor inhibitor cannabidiol, the topoisomerase II inhibitor etoposide, the 26S proteasome inhibitor bortezomib and the PLK1 inhibitor BI-6727 (Figure 2C). In other instances, higher compound concentrations elicited either completely different signatures (e.g., rotenone, see below) or non-specific general signatures (e.g., phospholipidosis or apoptosis, see below). These results illustrate the variable effects that compound dose can have on biological responses.

### Prediction of mechanism, off-target effects, and specificity

Similar CGI profiles for different compounds strongly predicted MOA. To minimize batch and population doubling effects, we defined screen similarity as the Jaccard index (overlap over the union) of hits (FDR <0.05) in the same direction and genes with CRANKS ≥2 or ≤-2 (Table S5). The variance in this measure of similarity is relatively unaffected by the above-mentioned effects as compared to the Pearson correlation coefficient (Figures S5F and S5G). The same compound at different concentrations, structurally similar compounds (Tanimoto similarity > 0.8) and compounds with known shared MOA generated more similar chemogenomic profiles than other pairs of screens (Figure 2D). A network of screens with a Jaccard similarity of ≥0.1 often grouped compounds as expected but also frequently clustered multiple compounds with different known or putative MOAs (Figure 2E). For example, the mitochondrial complex I inhibitor phenformin^46^ clustered with the putative NF-κB inhibitor QNZ/EVP4593, the putative p97/VCP inhibitor NMS-873, and the EGFR inhibitor mubritinib, all of which have been reported to inhibit mitochondrial complex I.^23,47,48^ In contrast, the potent p97/VCP inhibitor CB-5083 conferred a specific signature that NMS-873 did not share (Figure S5H). Other compounds, including the putative mitochondrial fission inhibitor MDIVI-1, the putative Wnt agonist BML-284, the uncharacterized compound UM0131593, and the uncharacterized biaryl compound (R)-(+)-1,1’-binaphthyl-2,2’-diamine (R-DABN), had pronounced unexpected effects as microtubule polymerization inhibitors and clustered with known anti-mitotic agents (see below). The specificity of CGI profiles was illustrated by a screen with the S-DABN atropisomer (DABN is an axially chiral compound due to steric clashes around a rotatable bond), which had no hits above the FDR threshold and no similarity to the R-DABN atropisomer profile (Figure S5I). Numerous other examples revealed compound mechanisms by clustering of CGI profiles (pairwise profile similarity scores for screens are provided in Table S5 and at https://crispr.tyerslab.com, see Resource Availability). Compound specificity was further illustrated by CGI profiles for two structurally related inhibitors of the E1 activating enzymes for the ubiquitin-related modifiers NEDD8 (NAE1, inhibited by MLN4924, a.k.a. pevonedistat) and SUMO (SAE1, inhibited by ML-792). Each inhibitor forms a sulfamoyl adenosine adduct with its respective E1^49,50^ and exhibited unique CGI hits specific to NEDD8 and SUMO, as well as common hits such as the transcriptional co-repressor BEND3,^51^ which may reflect an intersection of NEDD and SUMO biology (Figure S5J).

### Multi-drug sensitivity and resistance

Gene interaction degree in our CGI dataset approximated a power-law distribution (Figure S6A).^52^ Most hub genes that interacted with many different compounds were strongly biased to be either SSLs or rescues (Figure 3A), termed monochromatic interactions.^53^ For instance, negative mTORC1 regulators, including components of GATOR1 complex (NPRL2, NPRL3, SZT2) and AMPK (PRKAA1) were frequent SSLs while multiple structural components of mTORC1 and mTORC2 (RPTOR, MLST8, MAPKAP1) were frequent rescues (Figures 3B and 3C). These results suggest that reduced mTOR signaling enables cells to survive compound-induced stresses. Similarly, multiple apoptotic mediators (BAK1, TP53, MARCH5, VDAC2, APAF1, FBXO11, UBQLN1, CYCS, CASP9) rescued the toxicity of a variety of compounds (Figures 3C and S6B). An interesting exception to the trend of monochromatic multi-drug genetic modifiers was the CTLH/GID ubiquitin ligase complex, subunits of which exhibited numerous positive and negative genetic interactions (Figure 3D). The CTLH complex targets a multitude of substrates for degradation via different substrate-specific adaptors^54^ and likely exhibits diverse genetic interactions that reflect the functions of its dominant substrates (see also mTOR section below).

**Figure 3.**
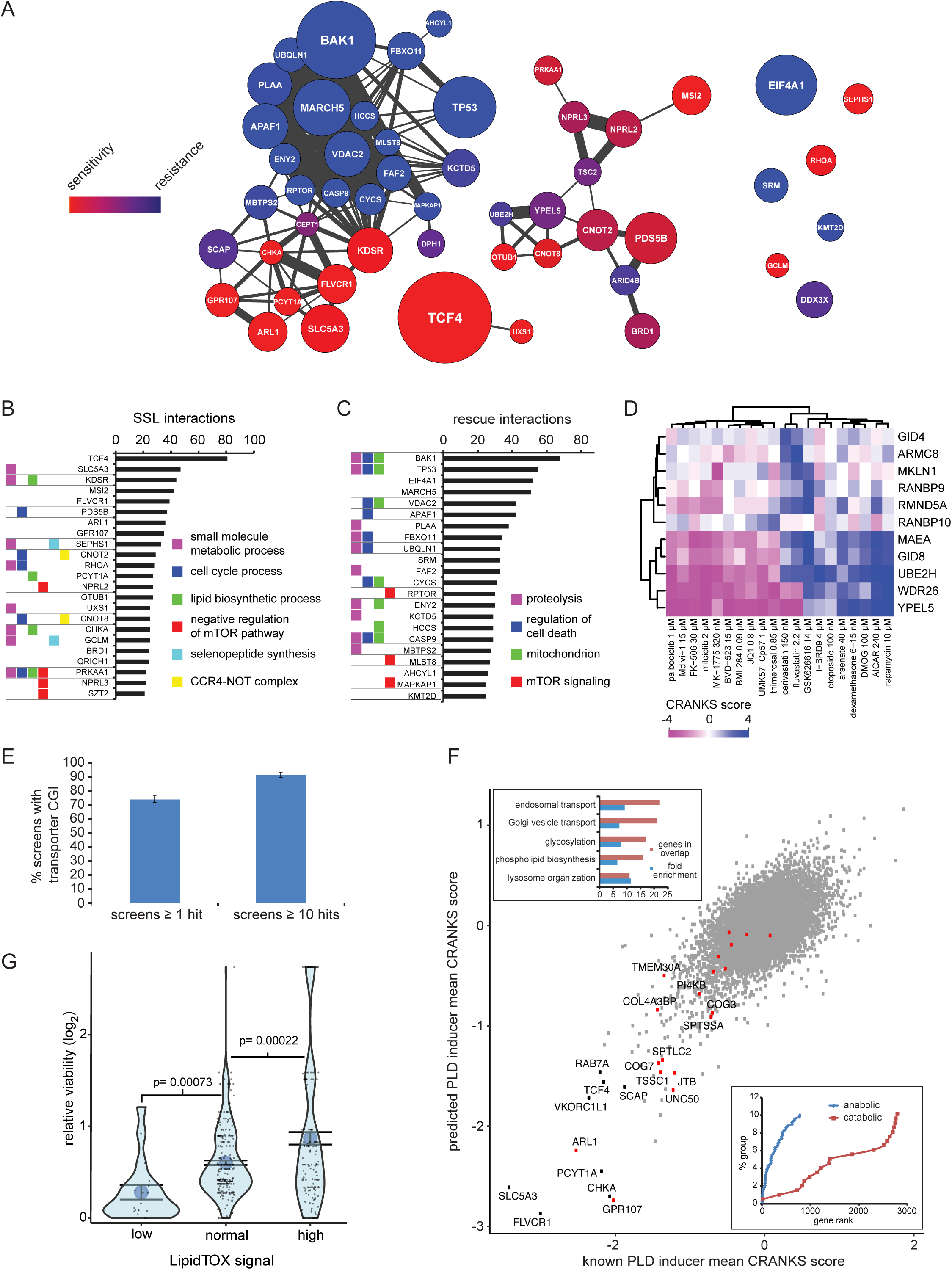
Multi-drug resistance and sensitivity genes. (A) Network of multi-drug sensitivity/resistance genes with correlated CRANKS scores (R≥0.5). Node size is proportional to the number of CGIs, line width to the strength of the correlation and node color to the ratio of SSL and rescue CGIs. (B) Genes and functions that frequently confer chemical sensitivity when deleted. Genes with 20 or more SSL interactions were annotated with selected Gene Ontology terms enriched in these genes (FDR <0.05, Fisher’s exact test). (C) Genes and functions that frequently confer chemical resistance when deleted. Genes with 25 or more rescue interactions were annotated with selected Gene Ontology terms enriched in these genes (FDR <0.05, Fisher’s exact test). (D) Subunits of the CTLH ubiquitin ligase complex exhibit rescue and SSL genetic interactions. (E) Frequency of transporter gene hits in screens with either ≥1 (327 screens) or ≥10 total significant CGIs (210 screens). (F) Mean CRANKS scores across phospholipidosis (PLD) gene cluster screens for known versus predicted PLD inducers. Red dots indicate genes identified by Kumar et al (2020)^58^. Upper inset, GO term enrichment for the top 200 SSL genes by mean score across the entire PLD cluster. Lower inset, enrichment of genes annotated as exclusively involved in lipid synthesis as opposed to lipid catabolism in the PLD cluster. (G) Elevated phospholipid levels correlate with improved cell viability after compound treatment. Relative viability was calculated as the ratio of absolute cell counts in specific wells versus the lowest cell count observed for the compound. LipidTOX Red signal indicated phospholipid levels. See also Figure S6.

Two multi-drug sensitivity genes, FLVCR1 and SLC5A3, are transmembrane transporters that are likely to mediate export of many drugs. As a class, at least one transmembrane transporter was a hit in 91% of screens with ≥10 hits in 74% of screens with ≥1 hits (Figure 3E). These CGIs may link specific compounds to their putative transporters and implicate a number of poorly characterized transporters in compound import or export (Table S6). Previously reported drug-transporter pairs were 2-fold more likely to score (≥1, ≤-1) than unreported pairs (p=0.00043, Fisher’s exact test Figure S6C). For example, most compounds that exhibited interactions with the multi-drug transporter ABCC1 were previously documented substrates (Figure S6D). The identification of transporter substrates through chemical-genetic screens reveals an important determinant of drug sensitivity and resistance, as well as endogenous metabolites.^55^

### A general phospholipidosis response signature

We hypothesized that compounds with similar physico-chemical properties may elicit a common genetic signature due to non-specific effects. Among 31 predicted molecular properties, dose-to-solubility ratio correlated strongly (|R| ≥0.25) with CRANKS scores for the greatest number of genes (Table S7). Top multi-drug sensitive (SSL) mutants, including FLVCR1 (3^rd^), GPR107 (4^th^), PCYT1A (12^th^), CHKA (24^th^) and SLC5A3 (26^th^), were amongst the most correlated genes. For example, the heme transporter FLVCR1^56^ and the GOLD domain seven-transmembrane helix (GOST) protein GPR107^57^ were strongly SSL across compounds with high dose-to-solubility ratios (Figures S6E and S6F). The most correlated genes had CGIs with many different compounds (Figure 3B), with SSL interactions heavily enriched for functions in membrane lipid metabolism (i.e., phospholipids and sphingolipids) or the endomembrane system (i.e., Golgi, endoplasmic reticulum (ER), and vesicle trafficking) (Figure 3F, top inset). These genes form a module that often dominated CGI profiles and clustered a diverse set of 28 compounds (Figure 2E). For example, 71/110 SSLs in a screen with a high concentration of the RAF kinase inhibitor vemurafenib were annotated with GO terms that included Golgi, vesicle or lipid metabolic process (Table S8, p=8.6e-17, Fisher’s exact test). This cluster was highly enriched in compounds with high dose-to-solubility ratios (Table S7, p=3.6e-8, t-test).

We suspected that phospholipidosis (PLD), a common cause of false positive results in cell-based screens,^58,59^ might explain the observed cluster. PLD is thought to be driven by the accumulation of cationic amphiphilic drugs (CADs) in the lysosome due to compound protonation leading to perturbed lysosomal lipid metabolism and phospholipid accumulation.^60-63^ Indeed, of 7 compounds in our dataset that included or were structurally very similar (Tanimoto similarity > 0.8) to verified PLD-inducers,^59^ all were in this cluster (p=4.2e-6, Fisher’s exact test; Table S9). Compounds in the cluster also tended to fit the definition of CADs, i.e. with higher predicted pKa and logP compared to other compounds in the dataset (p=0.0088, U test, and p=0.0036, t-test, respectively, Figure S6G; Table S9). Most of these compounds were screened at high concentrations (≥5 µM), often much higher than the known on-target Kd. We found that 12 of 19 top sensitizers in genetic signatures of 6 known PLD-inducing CADs in genome-wide CRISPRi and gene-trap screens^58^ were shared with the top 134 sensitizers in our *de novo* cluster (GPR107, ARL1, UNC50, JTB, TSSC1, COG7, SPTLC2, COL4A3BP, SPTSSA, COG3, PI4KB, TMEM30A; p=4.3e-21, Fisher’s exact test; Figure 3F; Table S10). To test whether PLD was indeed responsible for the observed genetic signature, we imaged NALM-6 cells treated with 21 compounds from the PLD cluster and the fluorescent phospholipid-binding dye LipidTOX Red for 24 h. Of the 21 compounds, 17 had elevated fluorescence, consistent with phospholipid accumulation, 3 compounds decreased signal, and 1 had no effect (Table S11). We tested 8 additional compounds not in the PLD cluster, of which 3 caused a signal increase, 4 caused a decrease and 1 had no effect (Table S11). Thus, most but not all compounds in our *de novo* cluster appeared to induce PLD.

A fraction of compounds that induced PLD were not CADs^64^, consistent with the divergent physico-chemical properties of some compounds in our PLD cluster. Some outlier compounds affected membrane lipid metabolism (dimethylsphingosine), sphingosine-1 phosphate signaling (FTY720), lysosomal function (bafilomycin A1), or vesicular trafficking (wiskostatin) and were screened at lower (≤3 µM) concentrations (Table S1B), suggesting that the PLD genetic signature can also be elicited via specific targets. We found that dose-to-solubility ratio was a better predictor of the PLD signature than logP and pKa combined (p=3.6e-4, Fisher’s method), suggesting that protonation-independent mechanisms may also induce PLD. For example, the plasticizer bisphenol F resides in the PLD cluster despite lacking a proton acceptor site (Table S9) and bisphenol A caused phospholipid accumulation within 24 h (Table S11). While bisphenols are known endocrine disrupters,^65^ we suggest that PLD may also contribute to bisphenol effects at higher doses. We note that compound aggregation, another known confounder in chemical screens,^66^ reflects the dose-to-solubility ratio. Consistently, the kinase inhibitors vemurafenib and lapatinib, which have strong PLD signatures, aggregate at lower concentrations than the IC30 values used in our screens^67,68^ and induced PLD above their reported critical aggregation concentrations (1.2 µM, 0.6 µM, respectively; Tables S1B and S11).

With regard to PLD mechanism, SSL hits related to lipid metabolism in the PLD cluster were significantly more likely to have anabolic as opposed to catabolic functions (Fig 3F, lower inset, p=9.8e-11, U test). This functional distinction suggests that lipid accumulation may be a protective response to PLD inducers, as hypothesized previously.^69-71^ We suggest that low phospholipid levels may exacerbate the effect of inhibitors of lipid metabolism, whereas supra-physiological phospholipid synthesis may protect cells from compound-mediated toxicity. Indeed, reduced phospholipid levels in LipidTOX Red assays were associated with lower cell viability (p=7.3e-4, t-test), while higher levels were associated with enhanced viability (p=2.2e-4, Figure 3G). The large PLD signature defined here provides a basis to further unravel mechanisms of PLD induction and its protective function. For instance, the vitamin K epoxide reductase subunit VKORC1L1 is a strong SSL in the PLD cluster, consistent with a postulated link between vitamin K and sphingolipid metabolism.^72^

### A mitotic cluster classifies compound MOA and identifies off-target effects

Many compounds yielded mitosis-related CGI signatures including inhibitors of actin or microtubule (MT) polymerization, kinesin motor proteins, polo-like kinase 1 (PLK1), cyclin-dependent kinase 1 (CDK1), the anaphase-promoting complex/cyclosome (APC/C) and aurora kinase A/B (AURKA/B) (Figure 4A). The spindle assembly checkpoint (SAC) prevents chromosome mis-segregation by blocking anaphase until all kinetochores achieve bipolar attachment to the mitotic spindle.^73^ Incorrectly attached kinetochores promote assembly of the mitotic checkpoint complex (BUB1B, BUB3, MAD2L1), which then sequesters CDC20, a substrate receptor of the APC/C ubiquitin ligase that targets mitotic cyclins and securins (PTTG1/2) for degradation to trigger anaphase. High doses of compounds that disrupt spindle function, including clinically approved and experimental chemotherapeutics (e.g., MT poisons such as nocodazole and the kinesin inhibitor SB-743921) were rescued by loss of several known SAC co-factors (Figure 4B). Although SAC inactivation in the presence of severe chromosome mis-attachment causes lethal chromosome aneuploidy, genetic knockout of SAC components allowed an additional round or two of cell division that scored as ersatz rescues in our screens, which we termed zombie screens. Conversely, loss of SAC function sensitized cells to spindle disruption at low concentrations at low doses of MT poisons (e.g., low dose vincristine or vinblastine screens).

**Figure 4.**
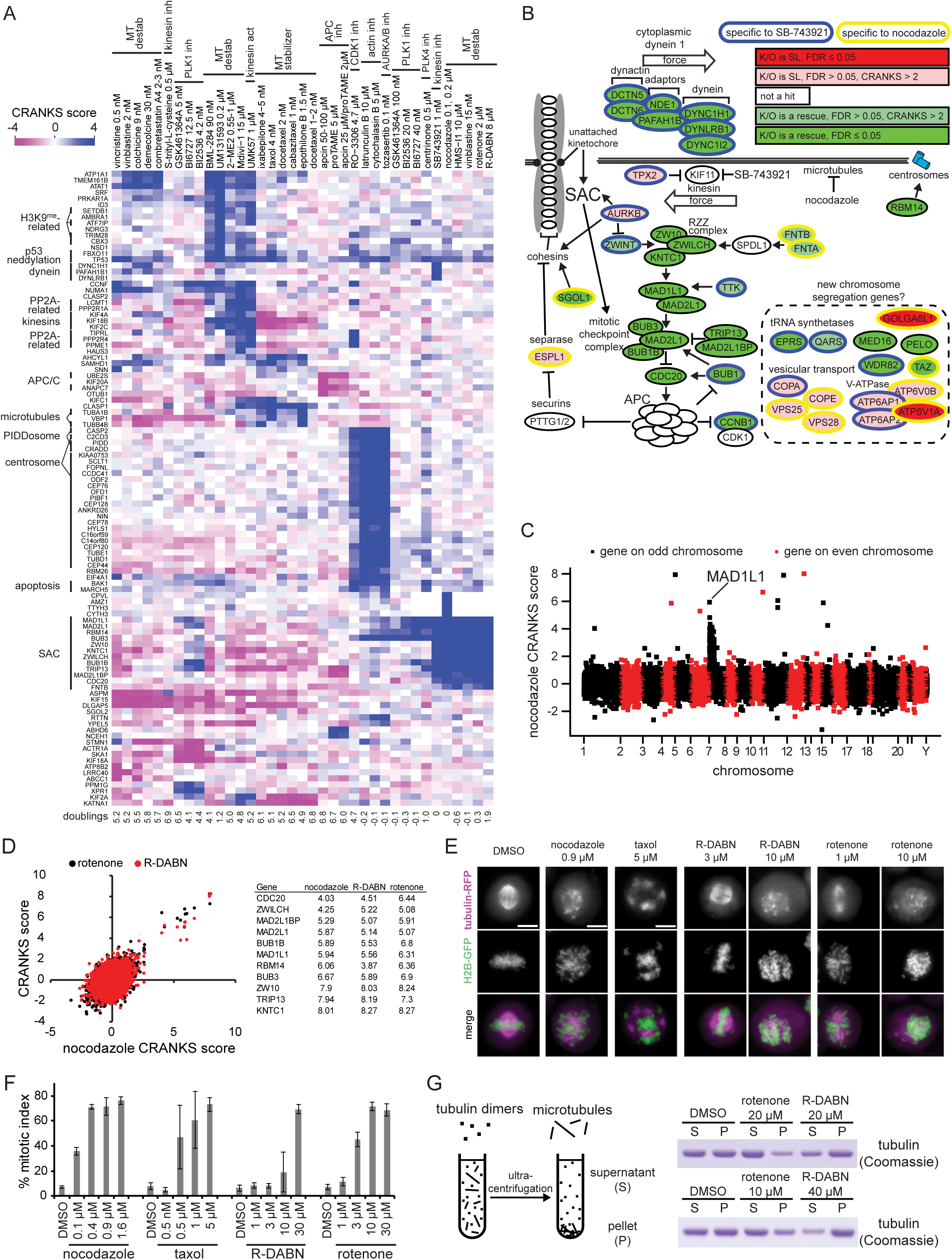
Mitosis network. (A) Heatmap of known mitotic inhibitor CGIs co-clustered with related signatures. Proliferation inhibition caused by each compound is indicated by population doubling. The average of three nocodazole screens (100 nM, 200 nM and 200 nM; Table S12A) is shown. (B) Mitotic network schematic of top hits for the microtubule poison nocodazole and the kinesin spindle protein inhibitor SB-743921. (C) Manhattan-style plot of gene scores in averaged nocodazole screens (Table S12A) as a function of chromosomal position. MAD1L1 resides on the distal end of chromosome 7P. (D) Comparison of R-DABN (8 µM) and rotenone (2 µM) profiles with averaged nocodazole screens (Table S12A). Scores are in order of top rescues in the combined nocodazole screens. (E) Effect of nocodazole, R-DABN and rotenone on chromosome segregation in RPE-1 cells. Taxol served as a positive control for spindle stabilization. Integrated H2B-GFP was used to visualize DNA and tubulin-RFP to visualize the mitotic spindle. Representative images are shown. (F) Quantitative mitotic indices for different doses of nocodazole, DABN and rotenone in RPE-1 cells. Taxol served as a positive control. (G) Effect of R-DABN and rotenone on microtubule polymerization in vitro. See also Figure S7.

Intriguingly, within the SAC zombie screens, the scores of all genes located on the 7p chromosome arm were highly biased toward rescues, including many significant hits (Figures 4C and S7A). As the SAC gene MAD1L1 is located at extreme distal end of 7p, we reasoned that CRISPR-induced DSBs might lead to chromosome arm loss and concomitant loss of the MAD1L1 locus (Figure S7B). No other SAC gene showed a positive score correlation with more proximal loci on the same chromosome, suggesting that MAD1L1 may be acutely limiting for the spindle checkpoint. The DHFR inhibitor aminopterin caused a similar but milder arm loss signature for the folate importer SLC19A1 on chromosome 21q (Figure S7C). The frequent recovery of arm loss events under selection has implications for CRISPR-based cellular therapeutics.^74^

Chemogenomic profiles segregated MT stabilizers, exemplified by paclitaxel (taxol), from MT destabilizers (Figure 4A). Notably, we screened paclitaxel and its derivative docetaxel at low clinically relevant nM concentrations that do not cause SAC-dependent mitotic arrest^75^ and uncovered many interactions with microtubule motors and other factors linked to mitosis. Destabilizers were further segregated into subgroups as colchicine site binders (colchicine, demecolcine, combretastatin) that clustered more closely to each other than vinca site binders (vincristine, vinblastine). The kinesin-5 inhibitors STLC and SB743921, the kinesin-13 agonist UMK57 (27829147) and PLK1 inhibitors also clustered with MT destabilizers driven in part by KIF15, KIF4A, KIF18B, and KIF2C. The MT cluster revealed off-target effects on MT dynamics for the respiratory complex I inhibitor rotenone, the WNT agonist BML-284^76^, the mitochondrial fission inhibitor Mdivi-1,^77^ the estrogen analog 2-methoxy-estradiol,^78^ and two uncharacterized chemical screen hits, UM131593 and R-DABN. Rotenone and R-DABN had similar SAC rescue profiles (Figure 4D) and were predicted to interfere with MT polymerization. Indeed, both compounds perturbed mitotic spindle formation, increased mitotic index and altered MT polymerization *in vitro* (Figures 4E-4G). It has been previously reported that a high concentration of rotenone inhibits proliferation due to MT depolymerization^79^ but this effect is not widely documented. A low dose of rotenone yielded the expected complex I inhibitor signature and consequent clustering with its derivative deguelin, and other complex I inhibitors noted above (Figure S7D). Mitotic CGI profiles also contained numerous other genes and functions, many not previously linked to mitosis, including AHCYL1, FBXO11, SAMHD1, TMEM161B, KLF16, LRRC8A, DPH1 and OTUD6B (Figures 4A and 4B).

The final phase of mitotic division, cytokinesis, occurs upon actin-dependent contraction of the midbody.^80^ Cytokinesis failure leads to supernumerary centrosomes, activation of the PIDDosome complex (PIDD, CRADD, CASP2), and consequent p53-dependent cell death.^81^ PIDDosome subunits were potent rescues of the AURKA/B inhibitor tozasertib, as well as the actin polymerization inhibitors cytochalasin B and latrunculin B. In addition to TP53, these screens recovered all 33 centrosome genes as rescues, including loci that encode the recently characterized TEDC1/2 complex,^82^ likely because improperly formed centrosomes can no longer signal to the PIDDosome (Figures S7E and S7F).

### An mTOR cluster reveals new metabolic links and inter-process cross-talk

The mechanistic target of rapamycin (mTOR) Ser/Thr kinase forms two distinct complexes: mTORC1, which stimulates protein, lipid synthesis and other anabolic processes to activate cell growth and proliferation, and mTORC2, which controls cytoskeletal organization, survival, and lipid and glucose metabolism.^83-85^ Profiles for rapamycin, a natural product that acts as allosteric inhibitor of mTORC1, and three active-site inhibitors (torin1, INK128, KU-0063794) that inhibit both mTORC1 and mTORC2^86^ yielded distinct signatures that mapped to established upstream regulators and downstream effectors of the expansive mTOR signaling network (Figures 5A and S8A; Table S12B). mTORC1 structural components and activators (MTOR, RPTOR, ragulator complex subunits LAMTOR1-5, Rag GTPases RRAGA/RRAGC) sensitized cells to all mTOR inhibitors. Rapamycin forms a ternary complex with mTOR and the immunophilin FKBP12^87^ and consistently FKBP12 rescued the anti-proliferative effects of rapamycin. Conversely, the rapamycin export pump ABCC1^88^ sensitized cells to rapamycin. The negative upstream regulator TSC2 conferred rapamycin resistance while PTEN and the mTORC2-specific subunits RICTOR and MAPKAP1^89,90^ rescued active-site mTOR inhibitors but not rapamycin. Some known mTOR downstream effectors, such as the S6 kinases RPS6KB1/2^84^ however failed to exhibit CGIs, presumably due to functional redundancy and/or the ability of other AGC kinases such as RSK to compensate.^91^

**Figure 5.**
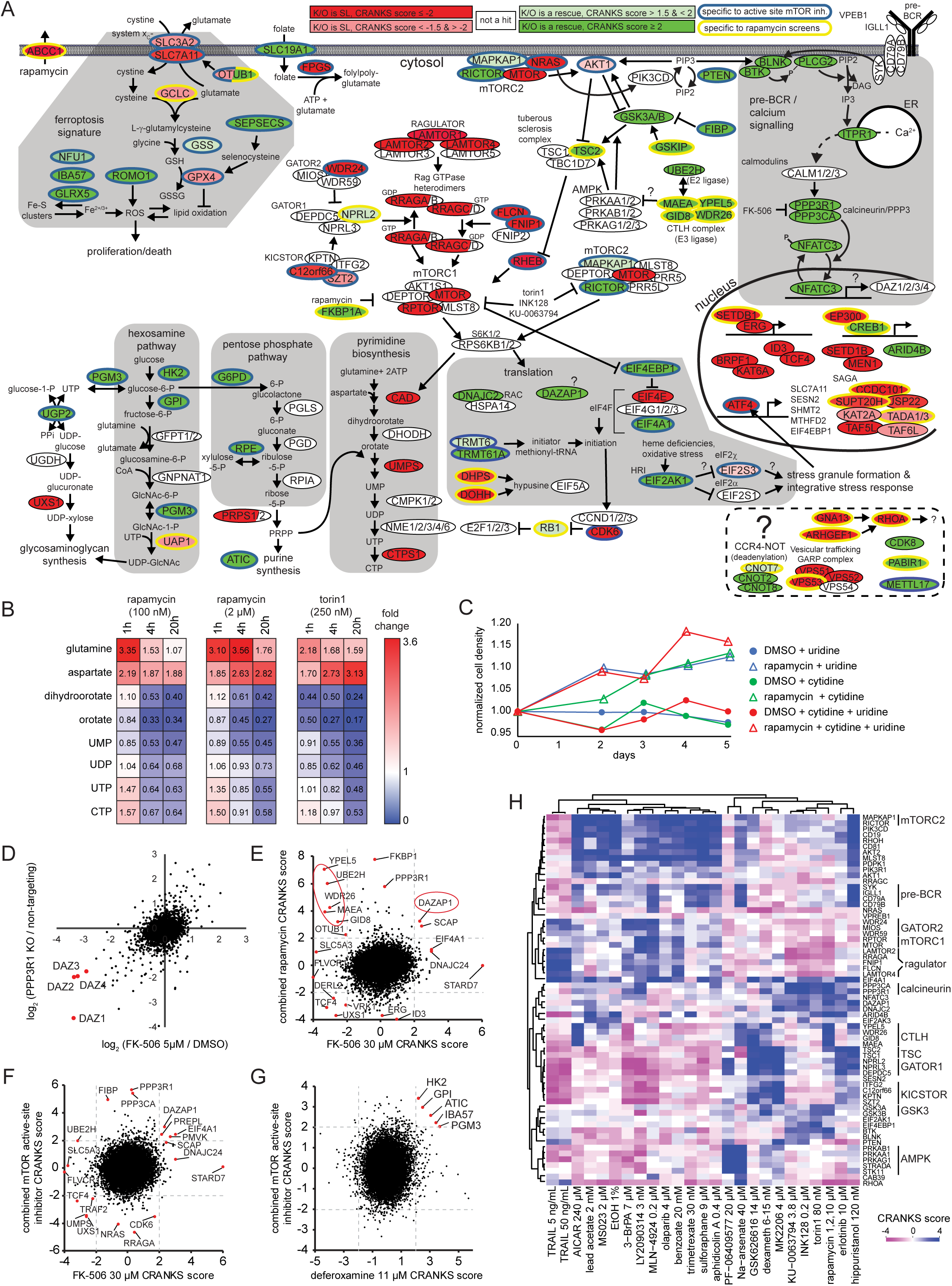
mTOR network. (A) Schematic of mTOR network indicating hits in diverse processes uncovered by CGI profiles of mTOR inhibitors. Hits specific for rapamycin are circled in yellow and for mTOR active-site inhibitors (INK128 0.2 µM, Torin1 80 nM, KU-0063794 3.8 µM) in blue. (B) Metabolomic analysis of pyrimidine biosynthesis metabolites in response to the indicated mTOR inhibitors. (C) Partial rescue of proliferation inhibition caused by rapamycin by supplementation with uridine and/or cytidine. Cell density was normalized to PBS-treated control cultures. See also Figure S8B. (D) RNA-seq analysis of PPP3R1 knockout versus WT NALM-6 cells (log_2_ fold change versus non-targeting guides) treated for 20 h with 5 µM calcineurin inhibitor FK-506 (log_2_ fold change versus DMSO). CTLH complex subunits are circled in red. (E) Comparison of CGI profiles for the calcineurin inhibitor FK-506 (5 µM) and triplicate independent rapamycin screens (1 µM, 2 µM, 10 µM). (F) Comparison of CGI profiles for the calcineurin inhibitor FK-506 and 3 combined active-site mTOR inhibitor screens (Torin1 80 nM, INK128 200 nM, KU-0063794 3.8 µM). (G) Comparison of CGI profiles for the iron chelator deferoxamine and 3 combined active-site mTOR inhibitor screens (Torin1 80 nM, INK128 200 nM, KU-0063794 3.8 µM) uncovers a shared hexosamine dependency. (H) Heatmap of compound CGI profiles that exhibit mTOR genetic dependencies. See also Figures S8 and S9.

Numerous CGIs revealed previously unknown aspects of mTOR signaling. Many screen hits mapped to metabolism, including the pentose phosphate, pyrimidine and hexosamine biosynthesis pathways (Figure 5A). CAD, a rate-limiting enzyme in pyrimidine biosynthesis downstream of mTOR,^92,93^ sensitized cells to all mTOR inhibitors. We validated these findings by measuring the levels of pyrimidine biosynthetic intermediates by GC/MS and found inhibitor treatment caused accumulation of intermediates upstream of CAD and reduction of intermediates downstream of CAD (Figure 5B). These data suggested reduced pyrimidine biosynthesis through suppression of CAD activity may at least in part mediate the anti-proliferative effects of mTOR inhibitors. Consistently, supplementation with cytidine, uridine or a nucleoside mixture partially alleviated anti-proliferative effects of rapamycin (Figures 5C and S8B).

The mRNA translation machinery showed both expected and unexpected interactions with mTOR inhibitors (Figure 5A; Tables S12B and S12C). eIF4E, the mRNA cap-binding subunit of eIF4F, sensitized to all mTOR inhibitors. EIF4EBP1, an eIF4E-binding protein that inhibits translation, rescued active-site mTOR inhibitors but not rapamycin, consistent with the inability of rapamycin to inhibit 4E-BP1 phosphorylation^94^ and 4E-BP1-dependent inhibition of proliferation by active-site mTOR inhibitors.^95^ DHPS and DOHH, which catalyze hypusination of eIF5A^96^, also sensitized to mTOR inhibitors. Conversely, the DEAD-box helicase eIF4A1 selectively suppressed active-site mTOR inhibitors. This result was surprising as mTORC1 is thought to indirectly stimulate eIF4A1 activity.^97,98^ However, eIF4A1 depletion may also stimulate mTORC1 by increasing intracellular amino acid levels due to reduced protein synthesis.^99,100^ The genetic bypass of mTOR inhibition by EIF4A1 may also reflect other roles of EIF4A1 such as in RNA stress granule formation.^101^

Surprisingly, the heme-regulated inhibitor (HRI) kinase EIF2AK1 sensitized cells to active-site mTOR inhibitors (Figure 5A; Tables S12B and S12C). HRI phosphorylates eIF2α to concomitantly downregulate global protein synthesis and translationally activate mRNAs with inhibitory upstream open reading frames, including ATF4 and CHOP.^102^ This CGI did not arise with other eIF2α kinases and suggested specific crosstalk between mTOR and HRI. ATF4 also sensitized to active-site mTOR inhibitors, consistent with previous links between mTORC1, ATF4, and the integrated stress response.^100,103,104^

mTOR inhibitor screens uncovered previously unappreciated connections between mTOR and calcium signaling (Figures 5A and S8A). Resistance to mTOR inhibitors was conferred by loss of several calcium signaling components, including subunits of the calcineurin phosphatase (PPP3CA, PPP3R1) and the transcription factor NFATC3 that localizes to the nucleus upon dephosphorylation by calcineurin. RNA-seq analysis of cells deleted for calcineurin subunits or treated with the calcineurin inhibitor FK-506 revealed that the deleted in azoospermia (DAZ) genes (DAZ1/2/3/4) located on the Y chromosome were downregulated (Figures 5D and S8C). Conversely, loss of DAZAP1, an RNA-binding protein that binds DAZ proteins^105^ conferred resistance to all mTOR inhibitors and also the calcineurin inhibitor FK-506 (Figures 5E and 5F). Intriguingly, deletion of DAZ genes and rapamycin treatment both cause male infertility^106,107^ and NALM-6 cells are derived from a male patient. Lifespan expansion by rapamycin is stronger in female versus male mice^108^ and female but not male RPSKB1 knockout animals exhibit lifespan extension.^109^ These results suggested that calcium signaling may interact with the mTOR network in a sex-specific manner.

mTOR inhibitor screens also revealed genetic connections with ferroptosis, a form of cell death triggered by insufficient glutathione (GSH) levels and iron-induced lipid peroxidation,^110^ which can be suppressed by reduction of mTOR activity.^111^ GPX4, a glutathione-dependent peroxidase that removes lipid peroxides, and components of the X_C_- antiporter that couples glutamate export to cystine import, exacerbated the anti-proliferative effects of mTOR inhibitors (Figure 5A). Genes implicated in Fe-S cluster assembly (GLRX5, IBA57, NFU1) also rescued active-site mTOR inhibitors, suggesting a role for free iron, possibly via effects on mTORC1 subunit expression.^112^ Given that mTORC2 is linked to iron metabolism,^113^ we screened the iron chelator deferoxamine and observed suppression by hexosamine pathway genes (HK2, GPI, PGM3), as also seen for mTOR inhibitor screens (Figure 5G). We speculate that free iron may alter heme levels, which in turn activate EIF2AK1 kinase to repress protein synthesis.

To explore the global patterns of mTOR genetic interactions across the entire chemogenomic dataset, we selected a set of 64 genes in the mTOR network and clustered all screens with hits in at least 9 genes within this set (Figure 5H). Many known mTOR subunits or regulators that scored only weakly with mTOR inhibitors scored strongly in other clustered screens including MLST8, GATOR1, GATOR2, KICSTOR, TSC1, AMP-dependent protein kinase (AMPK) subunits, and the pre-BCR/calcium signaling pathway.^114^ Phosphorylation of TSC2 by AMPK is thought to prime subsequent phosphorylation by GSK3^115^ and consistently, AMPK subunits strongly sensitized to the GSK3 inhibitor LY2090314 (Figures 5H and S3E). AMPK subunits rescued the AMPK activator PF-06409577 and arsenate, even though the latter is claimed to inhibit AMPK.^116^ Despite the shared AMPK subunit rescues, many other genes had opposite scores in the PF-06409577 and arsenate screens (Figure 5H). The CTLH ubiquitin ligase complex, noted above for its interactions with many compounds, has been proposed to regulate AMPK,^117^ and both CTLH subunits and TSC1/2 were strong rescues in rapamycin but not mTOR active site inhibitor screens (Figure 5A and 5H). CTLH genes were also potent hits in HMGCR inhibitor (i.e., statin) screens, in which the TSC1/2 genes were linked to ER-associated protein degradation (ERAD; Figure S9; Table S12C). Collectively, these genetic data support a CTLH-AMPK-TSC axis.

High concentrations of the nucleoside biosynthetic intermediate AICAR impair cell growth and survival, previously thought to be due to ADK-dependent conversion of AICAR into ZMP and activation of AMPK.^118^ However, AMPK subunits did not rescue in the AICAR screen (Figure S5C; Table S12D). Instead, one-carbon metabolism and purine biosynthesis genes were strong rescues, as well as CTLH subunits, mTORC1/mTORC2 subunits, and TP53 together with downstream apoptosis effectors (Figures S8D-S8F). Conversely, uridine-cytidine kinase UCK2, which increases pyrimidine nucleotide pools by salvage of pyrimidine nucleosides, and NT5C2, which converts purine nucleotides into nucleosides, were the strongest SSLs. These genetic interactions supported a recent model whereby AICAR causes purine-pyrimidine nucleotide imbalance, DNA replication stress, and apoptosis.^119^ This model also explains the potent rescue effects of ADK, AK2, and NUDT2 (Figure S8F). Given that mTOR appeared to activate *de novo* pyrimidine biosynthesis (see above), we were surprised that reduced mTORC1 and mTORC2 function rescued the effects of AICAR, as did CTLH subunits. These genetic interactions suggest the existence of additional regulatory circuits that control nucleotide balance.^120^

### A DNA damage response cluster uncovers mechanisms of DNA damage

The DNA damage response (DDR) mediates the detection, repair and downstream consequences of various forms of DNA damage.^121^ We screened 34 compounds with a known DDR-related MOA, from which we manually curated a list of 25 screens with a strong DDR signature (see Methods). We then generated a DDR heatmap from an unbiased list of 126 DDR-associated genes that exhibited CGIs with these 25 compounds (see Methods; Figure 6A). This CGI network recapitulated most of the known functional relationships in the extensive DDR network (Figure S10A) and provided new insights into compound MOA, as highlighted below.

**Figure 6.**
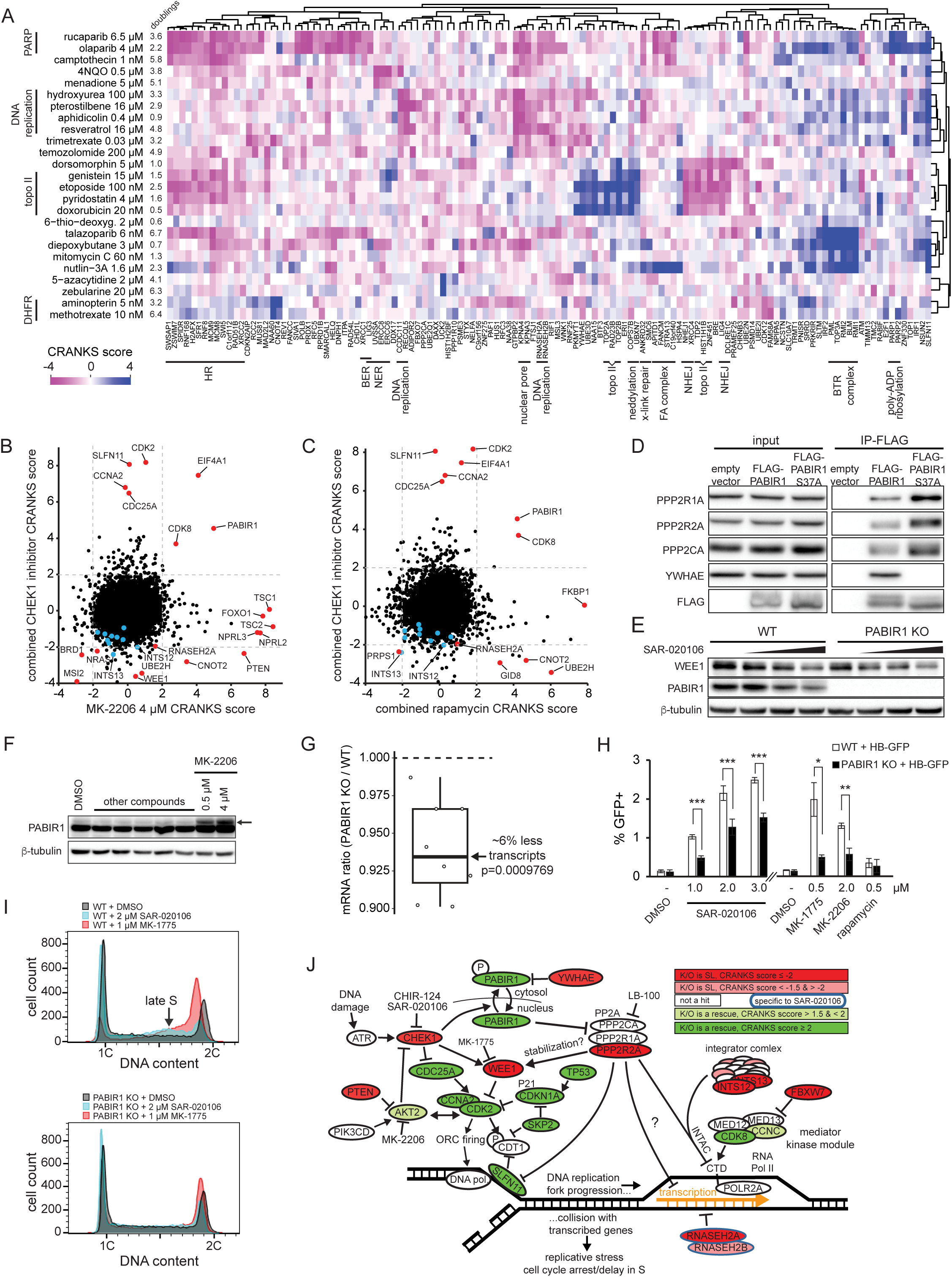
DNA damage response network. (A) Heatmap of CGI profiles for 25 DNA damaging agents and 126 DRR genes. An unbiased list of 126 DDR-related genes was compiled from a core set of screens (see Methods). Subclusters that reflect different aspects of the damage response are indicated. The DDR cluster unexpectedly included the putative AMPK inhibitor dorsomorphin. (B) Comparison of chemogenomic profiles for two combined CHEK1 inhibitor screens (SAR-020106 2.2 µM), CHIR-124 0.142 µM) versus the AKT1/2 inhibitor MK-2206 4µM. Blue dots indicate 10 of the 15 known integrator complex subunit genes. (C) Comparison of chemogenomic profiles for two combined CHEK1 inhibitor screens (SAR-020106 2.2 µM, CHIR-124 0.142 µM) versus three combined rapamycin screens. Blue dots indicate 10 of the 15 known integrator complex subunit genes. (D) Co-immunoprecipitation of PABIR1 with PP2A subunits and YWHAE. The PABIR1^S37A^ mutation abrogated YWHAE interaction and enhanced PP2A interaction, consistent with phospho-dependent retention of PABIR1 in the cytosol by YWHAE. (E) Wild type NALM-6 or a PABIR1 knockout clone treated with SAR-020106 (0, 0.125, 0.5 and 2 µM; 20 hours) did not exhibit increased WEE1 expression. (F) Induction of a high molecular weight form of endogenous PABIR1 (indicated by arrow) caused by treatment with the AKT1/2 inhibitor MK-2206. Other compounds with no effect were included on the same blot as incidental controls (the CDK8 inhibitors SEL-120-34A at 0.1 µM, BRD-6989 at 1 µM and BI-1347 at 0.01 µM; CDK1 inhibitors RO-3306 at 2 µM and 4 µM). (G) RNAseq-based ratio of total mRNA versus ERCC spike-in control RNA revealed that PABIR1 knockout had lower general transcription levels than wild type NALM-6. (H) CHEK1 (SAR-020106), WEE1 (MK-1775) and AKT1/2 (MK-2206) inhibitors increased R-loop formation as indicated by HB-GFP positive cells. Rapamycin served as a proliferation control. (I) The CHEK1 (SAR-020106 2.2 µM) or WEE1 (MK-1775 320 nM) inhibitors slow cell cycle progression in late S phase in a PABR1-dependent manner. DNA content distribution determined by propidium iodide FACS analysis. (J) Model of how general transcription instigates DNA replication stress. Hits from two combined CHEK1 inhibitor screens (SAR-020106 2.2 µM, CHIR-124 0.142 µM) are indicated. See also Figures S10 and S11.

The G-quadruplex (G4) stabilizer pyridostatin and the AMPK inhibitor dorsomorphin clustered with the topoisomerase II (TOP2) poisons etoposide, doxorubicin and genistein (Figure 6A). TOP2A and TOP2B were the strongest rescues of TOP2 inhibitors and pyridostatin, with very similar overall genetic signatures between the different compounds (Figure S10B; Table S12E). These results are consistent with recent studies that show pyridostatin and other G4 ligands may trap TOP2 in toxic G4 complexes, which subsequently cause transcription-dependent double strand breaks.^21,122^ The strong rescue of dorsomorphin by TOP2A/B suggested that dorsomorphin may also trap toxic TOP2-DNA complexes. Intriguingly, pyridostatin and dorsomorphin both contain quinoline-related moieties and other quinoline-containing compounds both inhibit TOP2 and stabilize G4 structures.^123^ TOP2 inhibitors appear to activate AMPK^124^ and dorsomorphin has AMPK independent anti-cancer activity.^125^ Dorsomorphin may thus directly inhibit TOP2 and indirectly activate AMPK.

Inhibition of dihydrofolate reductase (DHFR) by folate analogs potently disrupts all aspects of one-carbon metabolism.^126^ The analogs aminopterin and methotrexate are actively transported into cells, whereas the lipophilic analog trimetrexate is thought to enter cells by passive diffusion.^127^ The three analogs had different profiles (Figure 6A; Table S12F). Intriguingly, two genes that remove abnormal bases and process apurinic/apyrimidinic sites, the uracil DNA glycosylase UNG and the endodeoxyribonuclease APEX1 were specifically SSL with trimetrexate; another SSL gene, APEX2, encodes a nuclease that processes DNA 3’ blocks produced upon uracil excision (Figure S10C). These CGIs were consistent with misincorporation of uracil caused by depletion of thymidine upon DHFR inhibition, failure of which leads to genomic instability.^128,129^ Conversely, the organic anion transporter ABCC4 was a trimetrexate-specific strong rescue. These observations could inform clinical deployment of different DHFR inhibitors.^130^ We also observed that the transmembrane protein N-acetyltransferase NAA60^131^ and the folate importer SLC19A1 were strong rescues for aminopterin and methotrexate but potent SSLs with trimetrexate (Table S12F). We propose that SLC19A1 activity requires acetylation by NAA60 and that loss of SLC19A1 activity prevents aminopterin and methotrexate import but sensitizes cells to trimetrexate because of reduced folate import.

The PARP inhibitors rucaparib and olaparib generate toxic trapped PARP complexes on DNA^132^ and were rescued by PARP1, and to a small extent by PARP2, as well as by p53 and other DDR/apoptotic mediators, the ADPRHL2 mitochondrial PAR hydrolyzing enzyme, and subunits of the mTORC2 complex (Figure S4C; Table S12G). Sensitizers included the MCM8/9 DNA helicase that mediates repair of collapsed replication forks, the DNPH1 deoxyribonucleoside 5’-monophosphate glycosidase, the PRDX1 peroxide reductase, and various genes implicated in HR including the Shu complex. A recently characterized gene, C1orf112 (a.k.a. FIRRM), was SSL with PARP inhibitors, and has recently been shown to cooperate with FIGNL1 to stimulate RAD51 filament disassembly in homologous recombination.^133^ These data reveal the complex genetics of PARP inhibition. A third PARP inhibitor, talazoparib, shared many sensitizers but was not suppressed by loss of PARP1 but rather by all subunits of the BTR complex (BLM, TOP3A, RMI1/RMI2; a.k.a. the BLM dissolvasome), as well as by the PML nuclear body scaffold and multiple SUMOylation pathway enzymes (UBA2, SAE1, SENP1, SUMO2, UBE2I), all of which may impact HR.^134,135^ We reasoned that the difference between the PARP inhibitors was due to the lower dose of talazoparib used. Indeed, a screen at higher talazoparib dose recovered PARP1 as the top suppressor and concomitantly removed the suppressive effects of BTR, PML and SUMOylation genes (Figure S10D; Table S12G). We speculate that at low doses of talazoparib, Holliday junction migration catalyzed by the BTR complex stimulates collisions with trapped PARP and toxic strand breaks. Furthermore, since PML bodies process some forms of DNA damage and require SUMOylation of PML and other proteins,^136^ these steps may generate toxic branch migration structures. At higher doses of PARP inhibitor, the preponderance of trapped complexes may render BTR-mediated toxicity inconsequential.^137^ These results illustrate the complex nature of dose-response relationships and may be of importance in clinical applications of PARP inhibitors.

The CHEK1 kinase imposes cell cycle arrest upon activation of the DDR. CHEK1 inhibitors, such as SAR-020106 and CHIR-124, cause unregulated CDK activation that leads to unrestrained ORC firing, acute depletion of nucleotide pools, DNA replication stress and DDR-induced proliferative defects. SAR-020106 and CHIR-124 elicited highly similar signatures (Table S12H) with CDK2 and its cyclin CCNA2 as top rescues and the rate-limiting nucleotide synthesis enzyme PRPS1 as a strong SSL (Figure 6B and 6C). CDK2/CCNA2 also rescued MK-1775, an inhibitor of the CDK inhibitory kinase WEE1 (Figure S3B). PABIR1 (formerly FAM122A), which encodes an inhibitor of the PP2A phosphatase^138^, was also a strong rescue in the SAR-020106, CHIR-124 and MK-1775 screens (Figures 6B, 6C and S10E). As predicted, PABIR1 was a rescue in the PP2A inhibitor LB-100 screen (Figures S10E and S10F). PABIR1 is bound by 14-3-3 epsilon encoded by YWHAE^139^, which was SSL in the CHEK1 inhibitor screens. PABIR1 is phosphorylated by CHEK1 on S37, resulting in cytosolic PABIR1.^140^ We found that PABIR1 specifically interacts with PP2A subunits and YWHAE but that a PABIR1^S37A^ mutant lost the YWHAE interaction and bound PP2A subunits more strongly (Figure 6D). Unlike in A549 cells,^140^ we did not observe any alteration of WEE1 abundance in PABIR1 KO cells after treatment with SAR-020106 (Figure 6E). We observed a prominent higher MW species of PABIR1 upon treatment with the AKT inhibitor MK-2206 (Figure 6F), consistent with the observation that AKT phosphorylates and inactivates CHEK1^141,142^ and with the rescue of the AKT inhibitor MK-2206 by PABIR1 (Figure 6B).

PABIR1 strongly correlated with the transcription-activating kinase CDK8 across many screens, including as a rescue for the AKT1/2 inhibitor MK-2206 and rapamycin (Figures 6B and 6C). We posited that PABIR1 may prevent the dephosphorylation by PP2A of the C-terminal domain (CTD) of RNA polymerase II (POLR2A), which is phosphorylated by CDK8 to activate transcription.^143^ We further hypothesized that loss of CDK8 or increased PP2A activity may elevate nucleotide pools through reduced general transcription rate and thereby increase the pool of nucleotides to overcome lower CAD activity in the presence of rapamycin (see mTOR section above). PP2A controls general transcription through dephosphorylation of the CTD via the PP2A-integrator complex INTAC.^144^ INTS12 and INTS13 scored as SSL in the CHEK1 inhibitor screens (Figure 6B and 6C), consistent with a transcription suppression model for rescue of replication stress. RNA-seq profiles revealed a decrease in general transcript levels in PABIR1 knockout cells compared to WT (Figure 6G) although we were unable to detect gross changes in phosphorylation of the CTD upon CHEK1 inhibition or PABIR1 knockout (Figure S10G). Lower transcription rates may suppress CHEK1 inhibitor-induced replication stress by reducing RNA Pol II-replication fork collisions that generate R-loops and consequent stalled/collapsed replication forks.^145^ Using the hybrid binding (HB) domain of RNase H1 fused to GFP as an R-loop probe^146^ we found that indeed CHEK1 inhibition induced R-loop formation whereas the PABIR1 knockout reduced R-loop accumulation (Figures 6H, S10H and S10I). Consistently, both subunits of RNase H2, which degrades RNA:DNA hybrids, were SSL in the SAR-020106 screen (Table S12H). Strikingly, the PABIR1 knockout completely reversed the delay in late S phase caused by MK-1775 or SAR-020106 (Figure 6I). Finally, the DDR executioner SLFN11, which mediates sensitivity to many DNA damaging agents (e.g., as a rescue in many DDR screens above) and is thought in part to block and disassemble stalled replication forks,^147,148^ was a strong rescue in CHEK1 inhibitor screens (Figures 6B and 6C). Intriguingly, PP2A-mediated dephosphorylation of SLFN11 may stimulate its ssDNA-binding activity,^149^ and potentially be antagonized by PABIR1. In summary, we propose that PABIR1 inhibits PP2A thereby increasing general levels of transcription and exacerbating DNA replication stress through transcription-replication fork collision (Figure 6J).

### Conservation and cell type-specificity of genetic architecture

To explore how well our results in the NALM-6 cell line might generalize to other human cell types, we compared our DDR network to a CRISPR screen dataset generated with 27 different DNA damaging agents in the non-transformed TP53-null retinal epithelial RPE-1 cell line using the TKOv3 sgRNA library, as published by Olivieri et al.^21^ This previous study used lower compound doses and more population doublings to enrich for SSL interactions, as well as a different scoring scheme.^26^ Of the total number of hits across both datasets, 217/1581 overlapped (24% of either dataset) despite differences in DDR agents, drug concentrations, and cell line genotypes between the studies. The Olivieri dataset contained far more SSLs relative to our dataset (97.3% vs 26%, respectively). Notably, 30 out of the 31 screens in the Olivieri dataset were most similar to one of the 26 DDR screens in our dataset (p=2.8e-49, t-test, Figure S11A), again illustrating the power of CGI profiles to predict compound MOA and validating the Jaccard index for screen comparisons. Gene set enrichments unique to each dataset reflected compound and concentration differences between the datasets (Figure S11B). Gene expression differences between NALM-6 and RPE-1 explained why some hits failed to reproduce across the two cell lines (Figure S11C; Table S1D).^150^ A total of 9 compounds were common between the two datasets, allowing a direct comparison of 1,460 statistically significant chemogenomic interactions in aggregate (Figure S11D; Table S13). Despite substantial experimental differences noted above, most interactions were conserved in terms of the direction of the effect (1027/1,460; 70%). The correlation of these 1,460 pairs of scores was very significant (R=0.53, p=2.3e-105) and positive for each compound (R≥0.24). However, the magnitude of individual CGIs did not reproduce as well, with only a small fraction of significant hits shared (33/1,460; 2.3%). The wild type status of TP53 in NALM-6 but not RPE-1 cells, as well as the absence of SLFN11 expression in RPE-1 cells,^151^ explained the strong differential apoptotic signature between the studies and the paucity of SLFN11 interactions in the Olivieri dataset (Figure S11D). While false negatives may partly account for minimal overlap, the much greater differences than between NALM-6 replicate screens (Figure 1F) suggests that cellular context strongly influences the magnitude of CGIs. The qualitative conservation of chemogenomic interactions suggests that the structure of the DDR genetic network is similar in different cell types, while the relative dependence on different elements of the network appears to vary by cell type.

### Prediction of chemical synergism from CGIs

Chemogenomic profiles represent a potential input for prediction of synergistic compound interactions for drug discovery.^152^ To explore this concept with the CGI dataset, we first combined one compound with a known target and a second compound predicted to sensitize the loss of the same target according to its chemogenomic profile. We combined vemurafenib, which at the high concentration screened elicited a PLD signature, with different Golgi, autophagy, or vesicle trafficking inhibitors (golgicide A, bafilomycin A1, brefeldin A, wortmannin, myriocin, monensin) in 5x5 dose combination matrices. The Bliss model, which is based on the assumption that the combination of two compounds that act independently will exhibit a multiplicative effect on cell proliferation, was used to asses synergy.^153^ In 3-day viability assays, synergism occurred at specific dose combinations between vemurafenib and bafilomycin A1, myriocin, or monensin (Tables S14A-S14D). The effects for monensin were observed at all pairwise combinations (Figure 7A) with some combinations producing 50-fold lower viability than expected from Bliss independence. This strong synergistic effect was also manifest in melanoma cells (Figure S12A; Table S14E). We then selected 5 compounds with profiles most similar to that of vemurafenib (clomiphene, BH1, Q15, ETC159, bafilomycin A1) and tested for interactions with monensin in 5x5 dose combinations. This set of combinations produced significantly greater absolute Bliss scores (i.e., the log_2_ ratio of the observed viability over that expected for Bliss independence) than a control set of 11 non-specific combinations (Figure 7B; Tables S14C, S14D and S14F). These observed synergies may be mediated by a compromised PLD response in the presence of a potent vesicle trafficking inhibitor, analogous to the SSL dependencies on lipid metabolism and membrane-trafficking for PLD inducers. To systematically test this target-based approach, we used the list of target genes for known drugs used in our screens (Table S3), compiled a list of predicted synergistic and antagonistic drug combinations covering a broad range of mechanisms, and tested 15 of these predictions in 5x5 dose combinations (Figure 7B; Tables S14C, S14D, S14F and S14G). The observed absolute Bliss scores were significantly greater on average than those of the non-specific combinations, demonstrating the generalizability of this simple predictive approach for combination therapies and adverse drug interactions.

**Figure 7.**
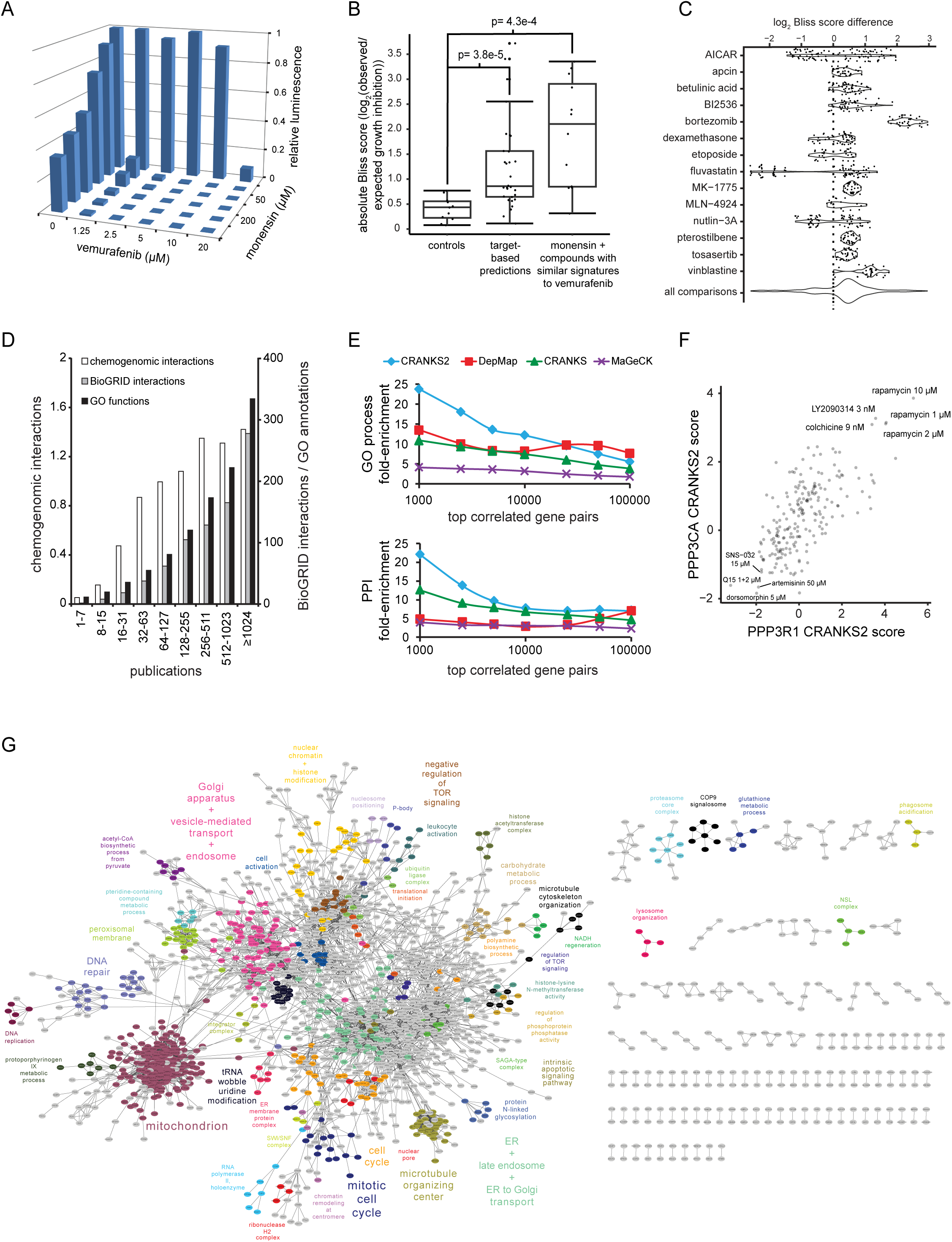
Prediction of compound synergy, protein-protein interactions and gene function. (A) Synergism between vemurafenib and monensin in NALM-6 cells. (B) Bliss scores of predicted interacting compound pairs. See main text for details. (C) Observed Bliss score differences between pairs of predicted antagonistic-synergistic pairs with the indicated training set compounds on the y-axis. See main text for details. (D) The CGI dataset was not biased against the under-studied genome, unlike protein-protein interaction (PPI) datasets and GO functional annotation. Number of publications from NCBI gene2pubmed were used as a proxy for the extent to which genes have been studied. (E) Prediction of shared GO biological process annotations (top) and protein-protein interactions (PPIs, bottom). CRANKS2, CRANKS and MAGeCK analysis of the CGI dataset were compared to the DepMap CRISPR gene essentiality screen dataset^6^. (F) Example of a strong correlation in CRANKS2 scores between two genes (PPP3CA and PPP3R1, the catalytic and regulatory subunits of protein phosphatase 3, respectively) indicative of a functional association. (G) Network of top 10,000 gene pairs with most correlated CRANKS2 scores across the CGI dataset. Annotated genes in different clusters are colored the same as the label. Highly correlated pairs of genes used to generate the network are provided in Table S15B. See also Figure S13A.

### Machine learning-based prediction of chemical synergism

We then used machine learning to explore the richness of chemogenomic profiles for synergy prediction without prior knowledge of compound MOA. We first quantified proliferation inhibition for a 15x15 compound pairwise single dose matrix (at approximately IC30 values; Table S14H) and calculated the Bliss score for each pair. We trained a multi-variate linear regression model based on CRANKS scores for each compound to predict Bliss scores with other compounds by multiplying learned gene-specific weights by all gene scores with absolute value ≥1 in the screens for the other compounds. Fourteen of the 15 models had significant predictive power based on internal cross-validation. For each of these models, we selected new compounds predicted to be either synergistic or antagonistic based on their chemogenomic profiles, excluding compounds with similar structures or reported MOAs to those in the training set. Viability assays were performed for each compound individually and in combination at the doses used in chemogenomic screens (Table S14I). We identified pairs of compounds tested in combination with the same query compound with a clear difference both between the two predicted scores (≥10% in relative rank) and the two observed scores (≥0.3), and plotted the difference between the observed Bliss score of the predicted antagonist minus that of the predicted synergizer (Figure 7C). Positive values indicated that the model correctly identified which of the two compounds were synergistic or antagonistic with the training set compound. These results showed that machine learning models can leverage large-scale CGI datasets to predict chemical synergism *de novo*.

### Insights into the unknome

The primary literature is heavily biased towards the study of a relatively small subset of human genes.^154-156^ The unbiased nature of our genome-wide screens revealed CGIs for much of the understudied protein coding genome (Figure 7D), thus providing entry points to explore new human biology. The least studied genes nevertheless produced fewer chemogenomic interactions on average, perhaps due to a relative lack of involvement in critical cellular processes or low expression in NALM-6 cells. The EKO library used in this screen is unique in its coverage of >20,000 alternatively-spliced exons and of >3,800 hypothetical genes annotated only as on-average short open reading frames (ORFs).^24^ Exons with CGIs were more highly expressed (Figure S12B) and skipped by fewer exon-exon junction reads (Figure S12C) in NALM-6 cells compared to exons with no CGIs. Conversely, the latter more often overlapped with known alternative splicing events (Figure S12D, p=2.6e-7, U test), consistent with tissue-specific functions not manifest in NALM-6 cells. Finally, hypothetical genes with likely CGIs were better supported by mass spectrometric evidence for protein expression (Figure S12E, p=3.8e-6, U test). Overall, these results align with gene essentiality properties of alternatively spliced exons and hypothetical genes and show how CGIs can illuminate gene structure and function.

### Accounting for proliferation inhibition improves predictive power

Strong correlations between gene scores across CRISPR viability screens are predictive of protein-protein interactions (PPIs) and gene functions.^157-159^ We compared different datasets and scoring algorithms for overlap between the most strongly correlated gene pairs at various cut-offs with known PPIs and shared GO biological process annotations (Figure 7E). Gene pairs with the most highly correlated CRANKS scores were significantly more predictive of PPIs and shared processes than those with highly correlated MAGeCK scores.^25^ Because CRANKS accounts for the fitness effects of essential genes that drive basal sgRNA depletion in pooled screens, we expected CRANKS to be superior to other methods when applied to our dataset. However, CRANKS assumes that essentiality rank alone can effectively model gene essentiality and implicitly that essential gene sgRNAs deplete linearly with population growth. We observed that the CRANKS scores of many essential genes showed weak but significant correlation with the number of population doublings (Figure S12F), indicating that residual fitness effects may bias scores. CRANKS scores for essential genes with sgRNAs that depleted relatively quickly from the pool (∼7days) were often over-corrected and tended to correlate positively with population doublings, whereas scores for essential genes with slowly depleted sgRNAs often had under-corrected scores and tended to correlate negatively with proliferation (Figures S12F and S12G). As we recorded proliferation inhibition for all screens, we computed for each gene the correlation between CRANKS score and proliferation inhibition, and then used this correlation as a second dimension to select a set of control sgRNAs for each gene, which mainly affected essential gene scores. We called this modified algorithm CRANKS2 and used it to recalculate all gene scores in our dataset (Table S15A). Control of residual bias for essential genes with CRANKS2 improved prediction of PPIs and shared GO biological processes (Figure 7E). The CGI dataset rescored with CRANKS2 outperformed a benchmark DepMap dataset of 769 cell line CRISPR screens (Figure 7E).^6^ In an illustrative example, scores for PPP3CA and PPP3R1 subunits of the calcineurin phosphatase strongly correlated over all screens with ≥10 hits and yet this correlation was not captured in the DepMap dataset (Figure 7F).

### Chemical genetic interactions reflect the genetic landscape of a human cell

Gene-gene correlation networks across large-scale genetic interaction or chemical-genetic datasets in budding yeast reveal the modular organization and hierarchy of cellular processes.^3,20^ We found that the top 10,000 gene pairs with the most correlated CRANKS2 scores in the chemical-genetic interaction dataset (PCC > 0.3882; Table S15B) formed a highly connected network of 2006 genes, with 1707 genes (85%) and 9779 edges (98%) residing in a single giant cluster. Many discrete sub-clusters within the network were enriched for specific GO (Figure 7G; Table S15B). Large clusters were annotated to mitochondrial, ER/Golgi/endosome, cell cycle/mitosis, chromatin/histone modification, and microtubule functions, while small clusters included those annotated to various aspects of DNA replication/repair, proteasome, COP9/signalosome, peroxisome, mTOR signaling, and metabolic processes. This genetic correlation map also contained dense regions without dominant GO annotations, potentially due to pleiotropic effect genes that coordinate process responses across different chemically-induced stresses. These features were robust to the correlation threshold, GO terms selected, and visualization method (Figure S13A).^160^ Despite the relative sparsity of the human gene correlation network compared to the exhaustively mapped yeast network,^3^ the clusters of GO terms were similar in the human and yeast networks (Figure S13B and S13C). Analogous to previous results in budding yeast, we found that essential genes were 3.2-fold enriched for genetic interactions compared to non-essential genes,^161^ and form the backbone of the gene-gene correlation network (42% of genes).^3^ These results underscore the overall conservation of genetic architecture in eukaryotes.

## Discussion

The CGI dataset reported here is the largest set of isogenic genome-wide CRISPR screens in human cells to date. This resource provides a valuable asset for interrogating the MOAs of known drugs and uncharacterized bioactive compounds, understanding gene function, and elucidating the genetic architecture of human cells. As first demonstrated in budding yeast, extensive chemical-gene interaction networks reflect the functional organization of the cell by virtue of correlated genetic response patterns to myriad perturbations.^20^ As we show, the CGI matrix provides new insights into cellular functions including lipid and nucleotide metabolism, mitotic regulation, mTOR signaling, and the response to DNA damage, as well as many other processes. These data complement recent efforts to directly map the genetic interaction landscape of human cells.^162^

The chemogenomics approach can inform and accelerate drug discovery through the systematic identification of compound on- or off-target effects and prediction of drug resistance mechanisms. Our chemogenomic screens identified many compound MOAs that contradicted assigned mechanisms in the literature. The magnitude of this problem in drug discovery has been uncovered through other systematic approaches^163,164^ and may be particularly acute for phenotype-driven screens.^165,166^ Chemogenomics represents a powerful strategy to guide structure-activity relationship (SAR) studies at all stages of the drug discovery pipeline. Hallmark chemogenomic profiles may be used to rule out less interesting hits from phenotypic screens and may also uncover new opportunities for drug repurposing.^167^ The integration of CGI data with patient genome sequence data also has the potential to prospectively minimize adverse drug interactions in patient sub-populations. Finally, as we show here, chemogenomic profiles enable prediction of drug synergies.^152,168^

A strength and at the same time a limitation of the CGI dataset is that our studies were carried out in a single human cell line. Isogenic screens obviate issues in comparing highly divergent cancer cell lines, and when coupled with a scoring approach that allows robust detection of genetic interactions of essential genes, yields data with a strong signal-to-noise ratio. However, some CGI profiles in our dataset undoubtedly reflect cell-type specific genetic networks and/or genetic aberrations specific to the NALM-6 cancer cell line used in these studies. While our analysis shows that judicious choice of compound concentrations can maximize recovery of CGIs for single screens, dose effects also warrant screens at two or more concentrations to ensure complete recovery of genetic rescues and sensitizers. A further limitation of our dataset is the nature of compounds selected for screens, which were inherently enriched for inhibitors of biological functions relevant to NALM-6 proliferation and/or survival. Despite these limitations, our CGI matrix outperformed other datasets for prediction of gene function. Our analysis also shows that while different cell lines may exhibit genetic differences in compound responses, most differences are in magnitude only and reflect an underlying core genetic architecture. Extension of the CGI matrix to new compounds that target additional diverse processes, multiple sentinel cell lines, and more complex phenotypic readouts should assign function to many less-well characterized genes and reveal how cell type-specific genetic landscapes dictate responses to chemical perturbation.

## STAR METHODS

### KEY RESOURCES TABLE

See attached table.

### RESOURCE AVAILABILITY

#### Lead contact

Further information and requests for resources and reagents should be directed to and will be fulfilled by the corresponding author Mike Tyers.

#### Materials availability

The unique reagents generated in this study are available from the lead contact Mike Tyers.

#### Data and code availability

- All supplementary tables are available through Mendeley Data at http://doi.org/10.17632/zdnrtmyfgh.1
- All sequence data and read counts are deposited in the NCBI Gene Expression Omnibus (GEO) repository under accession numbers: GSE315547 (CRISPR screens) and GSE316956 (RNA-seq).
- Original code generated in this study is available on GitHub at https://github.com/JCHuntington/CRANKS.
- All screen metadata, including compound suppliers, compound structures and characterization, dose-response curves, population doublings, CRANKS scores, CGI profiles, hit annotation, GO term enrichment, and screen comparison plots are available at https://crispr.tyerslab.com.
- Any further information required to reanalyze data is available upon request from the lead contact.

### EXPERIMENTAL MODEL AND STUDY PARTICIPANT DETAILS

#### Cell lines

The NALM-6 pre-B lymphocytic human cell line was kindly provided by Steve Elledge (Harvard Medical School). The generation of a NALM-6 clone (#20) expressing SpCas9 under a Tet-ON promoter was described previously.^24^ This clone of NALM-6 was verified for correct identity by whole genome sequencing and karyotype analysis. NALM-6 clone #20 was TP53^+/+^, MSH2^-/-^, NRAS^A146T^ and partially trisomic for Chr15. HEK293T and Jurkat (clone E6-1) cells were obtained from ATCC (catalog numbers CRL-3216, TIB-152). RPE-1 and RPE-1 cells expressing H2B-GFP and α-tubulin-mRFP were kindly provided by Helder Maiato (University of Porto). Cells were grown in 10% FBS (Sigma) RPMI (NALM-6, Jurkat) or DMEM (HEK293T, RPE-1) at 37°C in a 5% CO_2_ incubator unless indicated otherwise. A hypoxia screen at 5% O_2_ was performed in a BioSpherix chamber (screen 191). Screens at different temperatures were performed under standard conditions but at 35°C, 39°C and 42°C (screens 302, 303, 539). All cell culture was performed with the same lot of FBS (Sigma) except for a screen in which a suboptimal lot from Wisent was used (screen 426).

### METHOD DETAILS

#### EKO sgRNA library generation

The EKO pooled library of 278,754 different sgRNAs was used as previously described.^24^ The EKO library targets 19,084 RefSeq genes (∼10 sgRNA per gene), an additional 3,872 known or hypothetical protein-coding genes (∼10 sgRNA per gene), as well as 20,852 alternatively spliced exons derived from 8,744 genes (∼3 sgRNA per exon). A set of 2,043 randomly generated non-targeting sgRNA sequences with no detectable match to the human genome is also included in the EKO library. The entire set of 278,754 sgRNAs was synthesized as three 92K sub-library pools (CustomArray), amplified by PCR, cloned by Gibson assembly into the pLX-sgRNA-2XBfuA1-AAVS1 plasmid,^24^ transformed into ultracompetent DH5α *E. coli* at a representation of at least 100 colonies per sgRNA, and amplified on 15 cm LB+ampicillin plates at ∼50,000 colonies per plate. Plasmid DNA was recovered by maxiprep DNA purification.

#### Library transduction across screening rounds

The 428 screens reported in this study were carried out in 9 screening rounds over a period of 4 years (rounds 0 to 8) in which sets of compounds were screened in batches against the same thawed NALM-6 library pool (except in round 0,1 and 2 where we used freshly infected cells following blasticidin selection without freeze/thaw). For round 0, we used the identical NALM-6 library pool published previously.^24^ Upon thawing these cells for round 1, we observed a ∼50% mortality rate and thus discarded this batch and restarted screens with a new EKO library pool infection. From this second infection, we generated a new pool which we used for screening round 1. However, after completion of round 1, we noticed a slight library under-representation that yielded similar but less complex signatures for 24 compounds (which were re-screened in later rounds). Analysis of cell counts at each stage of pool generation showed that the second pool suffered from higher than desirable cell mortality 2 days after lentivirus addition. We therefore generated a third EKO library pool in the same manner but changed media 24 h after lentivirus addition to mitigate toxicity of HEK293T producer cell debris toward NALM-6 cells. Each batch of newly thawed cells from this third EKO library infection had minimal mortality and excellent representation. This library pool was used for round 2 (following blasticidin selection without freeze/thaw) and all subsequent rounds using thawed aliquots. Individual follow-up screens to test specific hypotheses also used the latter library pool using identical protocols to round 8.

#### EKO library lentivirus production

A three-plasmid expression system was used for lentiviral production in HEK293T cells. Lentiviral particles were generated by co-transfection of the two packaging plasmids psPAX2 (Addgene #12260) and pCMV-VSV-G (Addgene #8454) along with the EKO plasmid library into HEK293T cells, as previously described.^24^ Briefly, plasmids psPAX2 (151 μg), pCMV-VSV-G (82 μg), equal amounts of each of the three EKO sub-library pools (78 μg each) and 1.4 mg polyethyleneimine (1 mg/mL in water) were combined with water in a final volume of 16 mL. The resulting mixture was added to 550 mL of DMEM + 10% FBS and overlaid on 90% confluent HEK293T cells grown in a Corning HYPERFlask. After 16 h, media was changed to DMEM + 2% FBS and after 32 h lentivirus-bearing supernatant was collected, passed through a 0.45 μm filter and mixed with a 0.22 µm filter-sterilized concentrated stock solution to a final concentration of 5% sucrose, 2 mM MgCl_2_ and 10 mM HEPES pH 7.2. Harvested lentiviral preparations were stored at -80°C and used for three separate EKO library transductions of the inducible NALM-6 Cas9 clone #20,^24^ which were used for screens in rounds 0, 1 and 2-8 (see note on screening rounds above).

#### EKO library transduction

To transduce the EKO library into NALM-6 cells, 800 million cells of the inducible-Cas9 clone #20 were grown at 2 million cells per mL in a spinner flask supplemented with 10 µg/mL protamine sulfate 15 min prior to lentivirus addition at an MOI of 0.3. The next day, 100,000 cells were recovered to assess MOI by Q-PCR, as previously described.^24^ Briefly, genomic DNA was extracted from the library aliquot and, as a standard, from cells bearing pLX-sgRNA insertions of known copy numbers using the prepGEM DNA extraction kit (ZyGEM). This DNA was used as a template for Q-PCR reactions with primers q-PCR FWD 1 BLAST probe and q-PCR REV 1 BLAST probe matched with Roche’s Universal Primer Library #89 and reactions were performed on a ViiA7 Real-Time PCR system using the ViiA™ 7 software v1.2 (Applied Biosystems, Foster City, California, USA). Libraries were used only if the MOI was within 0.2 to 0.5. Then, cells were resuspended in new media (or not; see screening round note above) by centrifugation (400 x g, 15 min) and re-cultured in spinner flasks at 500,000 cells per mL. After 24 h, blasticidin (10 µg/mL) was added for library selection and expansion for a total of 6 days. After this period, approximately 6 billion transduced cells were obtained. Cells were pelleted and resuspended at 5 million cells per mL in freezing media (round 0: 20 % FBS, 10% DMSO & 70% RPMI, resulting in high mortality rates; round 1-8: 50 % FBS, 10% DMSO & 40% RPMI, resulting in low mortality rates) and 4.5 mL aliquots frozen slowly in 5 mL cryovials placed in thin-walled styrofoam-insulated boxes at -80°C overnight before transfer to liquid N_2_ for long term storage.

#### Genome-wide CRISPR knockout screens

A minimum of 70 million cells (250 cells per sgRNA) of the transduced library pool immediately following blasticidin selection (rounds 0,1, 2) or 1 day post-thawing when starting from frozen N_2_ aliquots generated during round 2 (rounds 3-8) were diluted to 400,000 cells per mL and supplemented with 2 µg/mL doxycycline to induce Cas9 expression starting at day -7 (see Figure 1A). Cells in culture were diluted back to 400,000 cells per mL on day -5 and day -3 with 10% FBS RPMI supplemented with 2 µg/mL doxycycline. On day 0, for each round, a minimum of 70 million cells (250 cells per sgRNA) were collected as day 0 post-doxycycline samples. The remaining cells were diluted with fresh media without doxycycline to 400,000 cells per mL and split into T-175 (175 mL, 70 million cells, 250 cells per sgRNA, round 0 and controls for all rounds) or T-75 (70 mL, 28 million cells, 100 cells per sgRNA, rounds 1-8) flasks for each screen. Flasks were then treated with the indicated concentration of each compound, as chosen to cause an estimated IC30 for proliferation inhibition (a.k.a. GI30), based on 384-well dose response curves for each compound. Most compound stocks were dissolved in DMSO, but a few (depending on compound solubility) were dissolved in PBS, water or ethanol. Unless indicated otherwise, concentrations of solvent used for each screen were adjusted to reach 0.1% (v/v) DMSO, 0.1% (v/v) water/PBS or 0.2% (v/v) ethanol. Exceptions are listed under technical notes in metadata column of Table S1A. For each round, a minimum of 70 million cells were left untreated as controls and a minimum of 28 million cells were treated with each solvent used for each round. As further controls, higher concentrations of DMSO and ethanol (at 0.3% v/v and 1% v/v) were also screened alone to identify potential specific CGIs for each solvent. DMSO at 0.3% v/v yielded a weak signature; DMSO at 1.0% slowed proliferation (4.13 population doublings) and yielded several significant hits; ethanol at 1% v/v generated a strong apoptotic signature. NALM-6 cells divide about once per 24 h and begin to saturate at a density of approximately 2 million cells per mL. Cell concentrations in each flask were monitored every 2 days (day 2, 4, 6 and 8) by reading 1 mL of the cell suspension with a Beckman Coulter Z2 particle counter/size analyzer (Beckman Coulter, Brea, California, USA) following a 1:10 dilution in Isoton II diluent. Every 2 days, a minimum of 70 million untreated cells were collected, washed in PBS and frozen as population drift controls (for rounds 2-8). On day 2, 4 and 6, whenever cell concentration exceeded 800,000 cells per mL for any condition, excess culture was discarded to leave 70 million (round 0) or 28 million (rounds 1-8) cells in the flask and volumes brought back respectively to 175 mL and 70 mL with fresh medium (i.e., to adjust cell concentration back to the starting density of 400,000 cells per mL). When cell concentration was below 800,000 cell per mL at each sampling day, incubation was continued for a further 2 days. For each diluted culture, compounds were added in the requisite amount to maintain the starting (IC30) concentration after dilution. Unless otherwise stated in the technical notes column of Table S1A, we assumed that added compounds were stable in cell culture. The number of population doublings obtained after 8 days of culture was calculated based on dilution steps (untreated cultures underwent between 7 and 8 population doublings in each round). At the end of day 8, all cells were pelleted, washed with PBS and frozen. All chemical and stress treatments were carried out for a total of 8 days with the exception of culture under hypoxic conditions (5% O_2_), which lasted for 12 days. In some cases, when compounds failed to cause sufficient inhibition of proliferation, we increased the concentration on day 4 (e.g., Q15 1-2 μM was screened at 1 μM on day 0 to 4 and 2 μM until day 8), as noted in Table S1A.

#### Genomic DNA extraction

Genomic DNA extractions for round 0 samples were carried out using a custom genomic DNA extraction protocol, as previously described.^24^ Briefly, frozen cell pellets were resuspended in 1.4 mL TE (10 mM Tris-HCl pH8.0, 1 mM EDTA) and lysis buffer (10 mM Tris-HCl pH8.0, 10 mM EDTA, 0.5% w/v SDS, 0.2 mg/mL proteinase K) was added to a final volume of 14 mL. Tubes were incubated for 3 h in a 55°C water bath with occasional vortexing. 5M NaCl was added to extracts to a final concentration of 0.2M, lysates were extracted twice with equal volumes of pH 7.5 phenol/chloroform/isoamyl alcohol (25:24:1) followed by chloroform extraction using phase-lock tubes. RNAse A (final concentration 50 μg/mL) was added and an overnight digestion carried out at 37°C. After phenol/chloroform and chloroform extraction, DNA was recovered by precipitation with 2.5 volumes ethanol and 1/30 volume 3M sodium acetate (pH 5.2). The pellet was washed with 70% ethanol, briefly dried and resuspended in 1 mL TE in a 55°C dry block, and finally sheared by several passes through a 27-gauge needle. Genomic DNA extractions of samples generated in subsequent rounds were performed using either the QIAamp DNA Blood Maxi kit (Qiagen) (rounds 1-2) or the Gentra Puregene Cell kit (Qiagen) (rounds 3-8) according to the manufacturer’s instructions.

#### Next generation sequencing

The sgRNA library was amplified in a first round of PCR from 462 µg of genomic DNA for samples of 70 million cells (i.e., given that a human diploid cell contains 6.6 pg DNA) in 575 µL 10X PCR buffer, 115 µL 10 mM dNTPs, 23 µL 100 µM primer Outer-1, 23 µL 100 µM primer Outer-2, 115 µL DMSO and 145 units of GenScript Green Taq DNA polymerase in a total volume of 5.75 mL. For samples of 28 million cells, 185 µg of DNA genomic template in the same PCR cocktail was used in 2.5X smaller volumes. Whenever the amount of gDNA for a sample was insufficient, all the gDNA was used and the number of PCR cycles increased accordingly to compensate. PCR reactions in 100 µL aliquots were setup in 96-well format on a T100 thermal cycler (BioRad, Hercules, California, USA) with an initial incubation of 95°C for 5 min, followed by 26 cycles of 35 sec at 94°C, 50 sec at 52°C and 40 sec at 72°C, and a final step of 10 min at 72°C after the last cycle. Completed reaction mixes for each sample were combined into one tube and 1.5 mL concentrated 15-fold by ethanol-precipitation using a 2.5X (v/v) EtOH 100%-PCR product ratio. Concentrated PCR products were loaded on a 1% agarose gel and the 475 bp amplicon extracted with a EZ-10 Spin Column DNA Gel Extraction Kit (Bio Basic) (round 0-6). Purification of PCR products was not carried out for rounds 7-8 as we determined this step was unnecessary. A second PCR reaction was performed to add Illumina sequencing adapters and 6 bp indexing primers using 250 ng PCR1 purified template for round 0-6 or 10 µL of 1:20 dilution of unpurified PCR1 product for round 7-8, 5 µL 10X buffer Takara (rounds 0-4) or 10 µL 5X buffer Kapa (rounds 5-8), 5 µL 2.5 mM dNTPs, 1 µL PAGE-purified equimolar premix 100 µM TruSeq Universal Adapters -2/0/+2/+5/+7 to shuffle the constant DNA region to be sequenced prior to the guide RNA sequence (round 0-3) or TruSeq Universal Adapter 0 (round 4-8), 1 µL of 100 µM PAGE-purified TruSeq Adapter with appropriate index, 1 µL DMSO and 5 units Takara Taq (rounds 0-4) or 5 units Kapa HiFi HotStart DNA polymerase (rounds 5-8) with volumes brought to 50 µL total in all cases. Reaction cycles were 5 min at 95°C, 5 cycles of 15 sec at 95°C, 30 sec at 50°C and 30 sec at 72°C, 5 cycles of 15 sec at 95°C, 30 sec at 56°C and 30 sec at 72°C, followed by 5 min final step at 72°C after the last cycle. A 236-245 bp amplicon (round 0-3) or 238 bp amplicon (round 4-8) was gel extracted from a 2% agarose gel (round 0-6) or purified using solid-phase reversible immobilization (SPRI) beads (AxyPrep FragmentSelect-I Kit) using a 1:1 ratio of SPRI beads and PCR product (round 7-8). Purified PCR products were quantified using the Infinite M1000 PRO microplate Reader (Tecan, Männedorf, Switzerland) and sequenced on an Illumina HiSeq 2000 with a read length of 50 nucleotides (SR50) (McGill University and Génome Québec Innovation Centre, Montréal, Canada; rounds 0-3) or on an Illumina NextSeq 500 in a 75 cycle single end high output flow cell, reading 43 bp with the 23 first bp read in dark cycles (IRIC Genomics Platform, Montréal, Québec, Canada; rounds 4-8). Average sequencing coverage was estimated at a minimum of 50 reads per sgRNA.

#### Data processing

Illumina sequencing reads were aligned to the designed EKO library^24^ using Bowtie 2.2.5 with default parameters and total read counts per sgRNA, including imperfect matches, were tabulated for each sample. For each round separately, read counts across all untreated or 0.1% DMSO or ethanol-treated control samples were added together to generate a single background sgRNA frequency distribution averaged across the 8-day screen. A small number of screens were split into separate cultures partway through the 8 day time course and then sequenced as two or more separate samples (Tables S1A and S1B). In these cases, the reads of all co-cultured samples were summed to generate a single sgRNA frequency distribution representing a single independent screen (Table S1B). The condition-specific RANKS algorithm (CRANKS) was used to generate gene scores and FDR values, comparing the read counts in each compound-treated sample to that in the background distribution. The minimum number of reads per sgRNA used in CRANKS was set between 20 and 200, depending on the depth of coverage of the pooled background distribution. Specifically, the minimal read threshold was set to 30, 20, 50, 50, 75, 95, 40, 100, 200 and 95 for rounds 0 through 8, and later individual screens, respectively. Otherwise, CRANKS was run under default parameters, including a minimum of 4 sgRNAs per gene passing the minimal read threshold. A few processed screens had inadequate library representation, in some cases because of excessive cell death caused by the compound treatment and in other cases due to technical reasons, such as poor DNA extraction or PCR amplification. To identify screens with insufficient library representation, we counted the total number of unique sgRNAs with at least one aligned read and applied a threshold of 260,000 sgRNAs (out of 278,754), corresponding to a maximum acceptable sgRNA dropout rate of ∼6.5%, below which we considered that the library was not adequately represented for sgRNA fold-changes to be reliably estimated. Based on this threshold, we discarded the results of 8 screens, which were not included in the final dataset. All but one of the compounds from these screens were re-run at lower doses to achieve acceptable quality screen data. The dataset used for all global analyses (e.g. Figures 1B, 1C, 1E, 2A, 2D, 2E, 3A, 3B, 3C, 3E, 4A, 5G, 6A and 7A-7G) was restricted to screens reported in rounds 0-8. 5 additional screens (cytarabine 16 nM, SAR-020106 2.2 μM, CHIR-124 142 nM, CHIR-99021 3.81 μM, talazoparib 55.2 μM; see Tables S1A and S1B) were run after round 8 to validate specific hypotheses.

#### CRANKS: Condition-specific Robust Analytics and Normalization for Knockout Screens

CRANKS is an extension to the RANKS algorithm, which has been described in detail previously.^24^ Briefly, RANKS estimates individual sgRNA p-values as the frequency of non-targeting control sgRNAs in the EKO library with a read frequency fold-change more extreme than itself and calculates the average log_e_ p-value across sgRNAs (i.e., the geometric mean) to generate the gene score. The newer version of RANKS, called RANKS-V2.0, combines sgRNA depletion and enrichment into a single score by averaging log(d)-log(e), where d is the fraction of control guides which are more depleted and e is the fraction which are more enriched (see https://github.com/JCHuntington/RANKS/tree/RANKS-V2.0). If a fraction equals to zero, d or e is instead set to 1 divided by the number of control guides. In CRANKS, control sgRNAs were instead defined as those targeting the 500 genes immediately more essential than the scored gene, thereby controlling for the proliferation-dependent dropout and greater variance in fold-changes of essential gene-targeting sgRNAs. For genes within the top 500 most essential, all genes scored as more essential than the gene itself were used to generate the control distribution. For genes within the top 50 most essential, all 49 other most essential genes were used. Gene essentiality ranking in NALM-6 was determined by applying RANKS on our previous data,^24^ comparing the day 0 to the day 15 sample, using the non-targeting control sgRNAs in EKO as the control distribution and the default 20-read minimal sgRNA coverage, including genes with as few as a single sgRNA passing the read coverage threshold. By controlling for the interaction between cell proliferation and essential gene knockouts, CRANKS can therefore score essential genes proportionally to their actual interactions with the compound/condition. A few genes (e.g., MYC) were extremely essential and generally had too few reads in both test and control samples to pass the critical thresholds to calculate a score. The CRANKS software and source code are available at https://github.com/JCHuntington/CRANKS. One ranking was generated for the core set of genes in the EKO library present in RefSeq and a separate ranking was generated for the extended sgRNA set including alternatively spliced exon targets and hypothetical genes. The core set ranking was used to generate the reported gene CRANKS scores while the extended set ranking was used only to generate alternative exon scores and hypothetical gene scores.

#### Compound sources and validation

Compounds were obtained from a combination of commercial suppliers, collaborators, and in-house chemical synthesis, as detailed in supplemental compound source and validation Table S1C. Chemical identifier information includes the chemical supplier product numbers, the Chemical Abstracts Service (CAS) registry numbers (https://doi.org/10.1021/ci60006a016) and PubChem Compound ID (CID) numbers (https://doi.org/10.1093/nar/gkv951). Chemical structure information is provided in Table S1C as SMILES (https://doi.org/10.1021/ci00057a005) and InChI string (https://doi.org/10.1515/ci.2006.28.6.12) notation. Concentrated compound stock solutions were prepared using DMSO, double-distilled water, or PBS1X as the solvent. The chemical identities of compounds were confirmed through a combination of high-resolution mass spectrometry (HRMS), HPLC-MS (LCMS), UV absorption, X-ray powder diffraction, and NMR. Analytical data used to confirm chemical identities is provided in Table S1C.

HRMS accurate mass measurements were recorded on an Agilent system (LC/MSD TOF model 61969A or an LC-TOF model 6224) equipped with an electrospray ionization source. Assay compound stock solutions were diluted with water (5%) in methanol to approximately 0.3 mg/mL concentration and injected into the mass spectrometer using a 0.5 mL/min flow of water (50%) in methanol. The capillary voltage was set to 3000 V and mass spectra were acquired from 100 to 3000 m/z. Protonated molecular ions (M+H)^+^ and/or sodium adducts (M+Na)^+^ were used for empirical formula confirmation in positive mode and deprotonated molecular ion (M-H)^−^ was used in negative mode.

Analytical HPLC-MS (LCMS) chromatograms were recorded on an Agilent system (HPLC model 1260, Quadrupole MS model 6120) equipped with an Atmospheric-Pressure Chemical Ionization (APCI) chamber and a reversed-phase column (Agilent Poroshell 120, EC-C18, 2.7 µm, 2.1 x 30 mm) using strong solvent mixture “B” [MeOH/ H_2_O/ AcOH (95: 5: 0.1)] in weak solvent mixture “A” [H_2_O/ MeOH/ AcOH (95: 5: 0.1)] as mobile phase, a flow rate of 1.50 mL/ min, and a linear gradient (0.0-0.5 min, 0 to 100% “B” in “A”; then 0.5-2.0 min, 100% “B”).

X-ray powder diffraction (PXRD) diffractograms^169^ were recorded using a Bruker Smart Diffractometer equipped with a Microfocus source delivering CuKalpha radiation (wavelength of 1.54178 Å) and a two-dimensional APEX 2 CCD detector. The collected data was summarized by listing the 2-theta angles and relative intensity of the principal peaks, defined as having a relative intensity of at least 10% compared to the major peak and 2-theta angles of ≤ 40°. Compound identity was confirmed by comparison of the patterns calculated from published crystal structure data or by a search-match procedure using pattern records from the Powder Diffraction File (PDF) database.

NMR spectra were recorded on a Varian 400-MR spectrometer tuned for hydrogen (^1^H) and carbon (^13^C) atom detection. Chemical shifts (δ) are reported in parts per million (ppm) relative to tetramethylsilane (0 ppm). Compounds were dissolved in deuterated solvents (D_2_O or CDCl_3_) and trace proteo solvent peaks were used as internal reference.^170^

#### Compound dose-response curves

To maximize recovery of genetic interactions, i.e., both rescue and SSL hits in single screens, we screened at compound doses estimated to cause 30% inhibition of proliferation (IC30, a.k.a. GI30) of the parental NALM-6 cell line. Prior to screens, all compounds were assessed for effects on proliferation in a 10-point titration curve. Panels of dilutions in steps of between 1:2 and 1:5 in quadruplicate with constant solvent concentration (0.1%) across all wells were performed at the IRIC High-Throughput Screening Platform (Montréal, Québec, Canada). A Beckman Coulter BioMek FX robot (Beckman Coulter, Brea, California, USA) (round 0-6) or an Echo 555 Acoustic liquid handler robot (Labcyte, San Jose, California, USA) (round 7-8) were used to distribute compounds in 384-well plates and either NALM-6 or NALM-6 Cas9 clone #20 (there was no phenotypic difference between the two lines) were added (50 µL per well at 200,000 cells per mL). Cells were grown for 72 h in a high humidity chamber installed in a 37°C 5% CO_2_ incubator. Cell proliferation was assessed by removal of 25 µL medium (taking care not to remove suspension cells that had settled at the bottom of wells) and addition of 25 µL CellTiter-Glo Luminescent Cell Viability Assay (Promega, Madison, Wisconsin, USA) to the remaining 25 µL of culture. Luminescence was read on a BioTek Synergy/Neo microplate reader (Promega) and statistical analysis was performed using IDBS Activity Base software (IDBS, Boston, Massachusetts, USA).

#### Single sgRNA-mediated knockout cell lines

Individual sgRNAs used in follow-up experiments were cloned into the LentiCRISPRv2 (Addgene #52961) or LentiCRISPRv2GFP (Addgene # 82416) plasmids in accordance with the Zhang lab protocol available at Addgene. Briefly, plasmids were digested with BsmBI and gel-purified using the EZ-10 Spin Column DNA Gel Extraction Kit (Bio Basic, BS654). sgRNA sequences were ordered together with the reverse-complement sequence as oligonucleotides (see Key Resources Table) from Integrated DNA Technologies (IDT, Coralville, Iowa, USA). Both forward and reverse oligonucleotides encoding sgRNAs were phosphorylated with T4 Polynucleotide Kinase (PNK) and annealed in a T100 thermocycler (BioRad, Hercules, California, USA). Annealed oligos were ligated with digested plasmids using T4 DNA ligase and transformed into Stbl3 competent cells. After amplification and DNA isolation of bacterial clones using a PureLink HiPure Plasmid Filter Midiprep Kit (ThermoFisher Scientific), sgRNA sequence insertions were confirmed by Sanger sequencing at the IRIC Genomics Platform. For single sgRNA lentivirus production, 6 μg psPAX2, 3 μg pCMV-VSVG and 9 μg LentiCRISPRv2 were mixed in 666 μL water with 53 µg polyethyleneimine. The transfection mixture incubated for 15 min at room temperature (RT) and then overlaid dropwise on top of 90% confluent 293T cells in 10 cm dishes in 10 mL of media. After 16 h, media was changed to 2% FBS-DMEM and after 32 h, lentivirus-bearing supernatant was collected, passed through a 0.45 μM filter and adjusted to a final concentration of 5% sucrose, 2 mM MgCl_2_ and 10 mM HEPES pH7.2. NALM-6 Cas9 clone #20 cells were transduced with lentiviral particles and selected as above to yield polyclonal knockout pools for each gene of interest. To generate scarless PABIR1 KO clones in WT NALM-6 cells were nucleofected (4D-Nucleofector, Lonza) with plasmid LentiCRISPRv2GFP encoding sgRNA 3B03 (GGAGTGGCTTATCTGCATTG) using the SF solution kit and recommended settings. 24 h post-nucleofection, single GFP+ cells were sorted into 96-well plates at the IRIC FACS Platform using a BD FACSAria IIu sorter and individual clones expanded over 4 weeks. Clones were assessed for PABIR1 status by immunoblot and candidate knockouts confirmed by RNA-seq (1 copy of the PABIR1 gene had a T insertion, the other an AAGAC insertion).

#### TRAIL hit validation

For validation of hits in the TRAIL screen, 1 million cells in 1 mL medium in 12-well plates were supplemented with 10 µg/mL protamine sulfate 15 min prior to addition of lentiviral particles made from LentiCRISPRv2 constructs (100 μL lentivirus preparation diluted to 1 mL with media). The next day, 2 mL media was added and 48 h post-infection puromycin selection (1 μg/mL) started for a period of 6 days. Cells were counted on a Beckman Z2 Coulter counter, diluted to 400,000 cells per mL and 25 μL of culture added to each well of a 96-well plate. A gradient of TRAIL concentrations in 25 μL medium was added to each well and viability measured 3 days later by CellTiter-Glo as described above.

#### Screen similarity and chemical similarity

The similarity between two screens was defined as the Jaccard index (i.e., dataset overlap over the dataset union) between the sets of genes with CRANKS scores above 2, below -2, or with an FDR<0.05, considering only genes with scores of the same sign in both screens as part of the overlap (Table S5). We used Tanimoto similarity as the measure of chemical similarity, which is defined as the Tanimoto (a.k.a. Jaccard) index between the sets of all chemical substructures encoded in each molecular fingerprint. Tanimoto similarities were calculated using Open Babel 2.4.1^171^ for all pairs of compounds that could be assigned a PubChem chemical ID or for which a chemical SMILES string could be generated. Similar pairs, defined as those with Tanimoto similarity of ≥0.8, are indicated in Table S5.

#### Prediction of compound physico-chemical properties

The SwissAdMe cheminformatics resource^172^ was used to generate quantitative predictions of 29 physico-chemical properties for each compound. Dose and calculated dose-to-solubility ratio (dose/ESOL) for each screen were also used as properties. The MolGPka web-tool^136^ was used to predict the pKa of each proton donor/acceptor group of each compound and only the most basic (i.e., highest pKa value) is reported in Figure S6G and Table S9.

#### Phospholipidosis signature

In the network of compounds with similar CGI profiles (i.e., Figure 2E), the cluster of screens connected directly or indirectly to clomiphene or vemurafenib (except through cannabidiol) was defined as the phospholipidosis (PLD) cluster based on the similarity of signatures and complete overlap with known PLD inducers. Cannabidiol was excluded since its profile is much more similar to that of THC, which is not directly connected to the cluster. We defined the average PLD signature as the average gene scores across all 28 screens in the cluster (Table S10). We used the Gene Ontology (version 2019-03-19) to define genes as directly and exclusively involved in either lipid biosynthesis or catabolism. We considered all genes part of GO biological process terms containing either the phrase “lipid biosynthetic process” or “lipid catabolic process” and excluded the term “regulation” to remove indirect effects. We also excluded genes shared by both lists so that the final gene sets would be exclusively associated with either anabolic or catabolic activity. Finally, we used the HUGO Gene Nomenclature Committee (HGNC) resource to map unmapped GO gene symbols to identifiers used in EKO. To assess whether biosynthetic genes were significantly enriched in the top SSL hits from the average PLD signature relative to catabolic genes, we compared the SSL ranks of the top 10% of genes from each category using the Mann-Whitney U test, which in the absence of enrichment, should produce a non-significant p-value, consistent with equivalent distributions.

#### Analysis of transporter genes

We defined transporter genes as the union of genes either annotated in Gene Ontology (version 2023-07-27) with any term containing “transporter_activity”, listed in the TransportDB 2.0 database (retrieved 2023-10-10),^173^, or with a gene symbol beginning with “ABC” or “SLC”. Reported small molecule-transporter pairs were retrieved from the VariDT 4.0 database (dated 2025-11-28).^174^ For Figure S6C, only genes and screens with compounds represented in the VariDT database were included, with genes that had an expression level <3 log_2_(reads-per-million) and screens with zero hits excluded.

#### LipidTOX Red assays and analysis

NALM-6 cells were grown for 24 h in 96-well plates with 10 doses of each compound in a series of 2-fold increments centered approximately on the screened (i.e., IC30) dose. Each treatment dose was tested in duplicate in separate plates with 40 untreated control wells. Cells were stained with LipidTOX Red dye (ThermoFisher H34351, 1:1000 dilution) and Hoechst DNA stain (0.02 µg/mL) then imaged in two channels (excitation laser 561 nm with a 640/40 filter; excitation laser 365 nm with a 390 nm UV excitation filter and 450/5 UV bandpass filter) on an Opera CHKN/QEHS ver.2.0.1.14101 confocal microscope (Perkin Elmer). Acapella 2.6 analysis software was used to identify cell nuclei (method M), remove border objects, assess which nuclei had appropriate area (25-80 µm^2^) and roundness (>0.8), estimate cell area (defined as the area centered on the nucleus with 60% greater surface), and quantify the average LipidTOX Red signal per cell. Based on the distribution of LipidTOX Red signal intensity values found in untreated wells (mean intensity 159.8, standard deviation 4.4) we applied thresholds 7 points above and below this average to classify compounds as causing a significant increase or decrease in LipidTOX Red signal, respectively. Both duplicate wells had to be above or below the threshold, which translates to a p-value of ≤0.0031 assuming a Gaussian null distribution, and a Bonferroni-corrected p-value of ≤0.031, given each compound was tested at 10 different concentrations. Cannabidiol was tested at 20 different doses and thus required a slightly stricter threshold, which it failed to reach despite marginally passing the usual threshold for lower intensity at a single dose (20 µM). A few compounds showed non-linear effects including both increased and decreased LipidTOX Red signal depending on dose. In these cases, we classified the compounds according to the largest absolute variation about the mean considering all doses tested. The complete list of measurements performed and the LipidTOX Red signal classification for each compound are in Table S11. Unlike control wells, treated wells tended to show greater variability in LipidTOX Red intensities, with most compounds producing extreme effects, especially at higher doses. We considered LipidTOX Red readings between 100 and 180 as normal, with low and high bins representing readings well outside the normal range. Cell viability was defined by the number of nuclei classified as viable for analysis compared to the total number of nuclei detected, with a maximum of 700 nuclei considered per condition. Only compounds with greater than 10% inhibition (<630 nuclei) in at least one well were considered for the analysis. To account for the different levels of compound toxicities, we expressed cell viability as the log_2_ ratio of viability for each well divided by the lowest viability observed for the compound at any dose.

#### Mitotic cell imaging

RPE-1 cells expressing H2B-GFP and α-tubulin-mRFP were plated at a density of 25,000 cells/well in 24-well plates containing poly-L-lysine (Sigma-Aldrich) coated round coverslips in each well. The following day, cells were treated with the indicated concentrations of compound for 17 h. Coverslips were then washed in PHEM buffer (60 mM Pipes, 25 mM HEPES, 10 mM EGTA, 4 mM MgSO_4_, pH6.9) pre-warmed at 37°C before fixation in PHEM buffer containing 4% formaldehyde (Sigma-Aldrich) for 20 minutes at 37°C. Coverslips were then washed 4 times (1 fast wash and 3 x 5 min washes) in TBS-tween 0.1% and mounted on slides using Vectashield media containing DAPI (Vector Laboratories). Images were acquired on an AxioImager microscope (Carl Zeiss) with a 63x oil objective (NA 1.4 DICIII) and an AxioCam HRM camera (Carl Zeiss) using AxioVision software (Carl Zeiss). To measure the mitotic index, RPE-1 cells were seeded at a density of 8,000 cells/well in 96-well, black, glass flat-bottom plates (SensoPlate, Greiner bio-one) in triplicates. The following day, cells were incubated for 17 h with compound diluted in DMSO at indicated concentrations or DMSO alone control. Cells were then fixed with 4% formaldehyde at 37°C for 20 min. Cells were washed 4 times as above in TBS-tween and then incubated in PHEM buffer containing 2% BSA (Sigma-Aldrich) and 0.1% triton for 1 h at RT for blocking and permeabilization. Cells were then incubated with anti-phospho-Histone H3 (pHH3, Ser10) (EMD Millipore) diluted 1:300 in PHEM buffer containing 2% BSA. Plates were incubated for 2 h at RT and then washed 3 times with TBS-tween. Alexa Fluor® 488-coupled anti-rabbit (Invitrogen) diluted 1:200 in PHEM buffer containing 2% BSA was added to each well and plates were incubated for 1.5 h at RT in the dark, followed by 3 washes with TBS-tween. Finally, a solution of Mowiol-containing DAPI was added and plates were kept at 4°C in the dark until imaging using a 60x water immersion objective on an Opera CHKN/QEHS ver.2.0.1.14101 confocal microscope (Perkin Elmer). DAPI and 488 nm fluorescence signals were acquired for 50 fields/well with 4 z-stacks. Using the Acapella 2.6 software, maximum intensity projections of the 4 planes were first generated and method C was used to select the nuclei population. The mitotic index corresponds to the percentage of pHH3-positive cells at 488 nm relative to the total number of DAPI-stained nuclei. Experiments were performed in triplicate. In vitro microtubule polymerization assays were performed as previously described.^175^

#### Microtubule polymerization assay

Protein preparation, polymerization and centrifugation-based sedimentation assays were performed essentially as described previously.^176^ Briefly, a 2X BRB80-based premix was prepared with 5X BRB80 (400 mM PIPES, 5 mM MgCl_2_ and 5 mM EGTA, adjusted to pH 6.8 with KOH), 2 mM DTT, 2 mM GTP and 20% DMSO. 5 µL of the mixture was added to separate microcentrifuge tubes with 1 µL of a 10X working dilution of each compound in DMSO for each concentration assayed. Recycled tubulin obtained from bovine brain (100 µM stock)^177^ was thawed on ice, 1 µL added to each tube, and the total volume adjusted to 10 µL with water. After brief vortexing, polymerization reactions were carried out in a circulating precision water bath at 37°C for 10 min. Samples were then diluted to 25 µL with 1X BRB80 buffer and polymerized microtubules pelleted by ultracentrifugation at 279,000 x g (80,000 rpm in a Sorvall TLA-100 rotor) for 5 min at 25°C. Supernatants and pellets were boiled separately for 5 min in equal volumes of 1X Laemmli buffer and resolved by 12% SDS-PAGE, followed by Coomassie Blue staining.

#### RNA-seq analysis

Following compound treatment or lentivirus infection plus selection, 1 million NALM-6 cells collected from an exponentially growing culture (800,000 cells per mL) were centrifugated for 5 min at 2,300 x g at 4°C, resuspended in 1 mL TRIzol (Ambion, Life Technologies) and samples processed by the IRIC Genomic platform. In the case of PABIR1 knockout versus wild type cells, cell concentration was measured using a Beckman Z2 Coulter counter in triplicate. ERCC RNA loading control Spike-in mix (Ambion Ref. 4456740) was diluted 1:20 in water and 20 µL per mL spiked in Trizol prior to addition of 1 mL over each cell pellet. Total RNA was isolated using RNeasy mini kit (Qiagen) according to the manufacturer’s instructions. RNA was quantified using Qubit (Thermo Scientific) and quality was assessed with the 2100 Bioanalyzer (Agilent Technologies). Transcriptome libraries were generated using the KAPA RNA HyperPrep (Roche) using poly-A selection (Thermo Scientific). Sequencing was performed on an Illumina NextSeq 500 at various read depths or on an Illumina Hi-Seq 2000 at a targeted depth of 50 million reads per sample.

To establish baseline expression levels, RNA-seq was performed on four NALM-6 clones expressing Cas9 and either an sgRNA targeting Azami green as a non-targeting control, the AAVS1 locus as a genome-cutting control, or the gene TLE4 that caused only minimal expression differences relative to wild type cells (Table S1D). Reads were aligned to Gencode V23 and RefSeq (January 2016) transcripts using Bowtie 2.2.5 with default parameters. Alignments with more than one inserted base, one deleted base or a total edit distance of >5 were discarded. Aligned read counts from the four samples were added together. Gencode alignments were used to estimate normal gene expression in NALM-6 cells (as log_2_ reads per million; Table S1D; data used in Figures 2A, 2D, S1C, S1D and S11C) while RefSeq, being the same reference used to annotate the core sgRNA set of the EKO library, was used for analyses on mutant strains, drug treatments, and alternative exon expression in NALM-6 (i.e., Figures S5B and S7A-S7D). The FK-506 treated sample gene read counts were compared to the DMSO-treated sample read counts, excluding genes with less than 100 reads in both samples (Table S16A). PPP3CA and PPP3R1 knockout cells were analyzed in the same manner, comparing each knockout population with a different sgRNA to a different wild type control (either wild type parental cells or a cell population targeted with an sgRNA against either the AAVS1 locus or the Azami-green sequence); the log_2_ fold-changes of the 3 comparisons for each gene were averaged together (Tables S16B and S16C). Prior to calculating log2 fold-change, read counts were divided by the 75^th^ percentile gene’s read count in the same sample, which is termed upper-quartile normalization, as suggested.^178^ Exon expression was defined as the fraction of reads aligned to the gene that overlap the exon. Exon-skipping was quantified by counting reads that overlap an exon-exon junction that excludes the exon. General transcription rate in PABIR1 knockout clones and matched controls was determined using the average ratio of reads mapped to ERCC spike-ins to the number mapped to RefSeq genes (Figure 6G). Given an equal number of cells and an equal amount of spiked-in RNA, this ratio should be inversely proportional to the amount of mRNA in an average cell. The transcription rate of each knockout relative to its matched control sample was therefore calculated as the ratio of the two ERCC/mRNA read ratios, i.e. mRNA^KO^/mRNA^WT^ = [ERCC-reads^WT^/mRNA-reads^WT^] / [ERCC-reads^KO^/mRNA-reads^KO^] (Table S16D).

#### Metabolomics

Experiments were carried on 30 mL of asynchronous NALM-6 cells at 400,000 cells per mL in T-75 flasks treated with 1000X DMSO solvent control, rapamycin (final concentrations of 100 nM and 2 µM) or Torin1 (final concentration 250 nM) for 20 h, 4 h or 1 h prior to sample collection. For GC-MS data acquisition, cells were centrifuged 7 min at 335 x g and washed three times with cold saline solution (9 g/L NaCl). Following centrifugation, cells were quenched on dry ice with 600 µL 80/20% HPLC-grade methanol/water that had been pre-chilled to -20°C. Samples were sonicated for 10 min, at 30 sec on/off intervals at 4°C on high setting (Diagenode Bioruptor, Cat#UCD-200 TM) and centrifuged at 13,000 rpm at 4°C for 10 min. The internal standard, myristic acid-D_27_ (750 ng, Sigma) was added to the extracts. Samples were dried in a cold (-4°C) vacuum centrifuge (Labconco) overnight. Dried samples were solubilized in 30 µL methoxyamine-HCL (10 mg/mL in pyridine) (Sigma, Cat#226904) through sonication (30 sec) and vortexing (30 sec), followed by centrifugation at 13,000 rpm for 10 min. Samples were transferred into autoinjection GC-MS vials and heated at 70°C for 30 min. Samples were derivatized with 70 µL *N*-*tert*-butyldimethylsilyl-*N*-methyltrifluoroacetamide (MTBSTFA) (Sigma, Cat#M-108) at 70°C for 1 h. A volume of 1 µL of the derivatized sample was injected inside an Agilent 5975 C GC-MS equipped with a DB-5MS+DG (30m x 250 µm x 0.25 µm) capillary column (Agilent J&W, Santa Clara, CA, USA). The injection was performed in the GC in splitless mode with an inlet temperature set to 280°C and electron impact set at 70 eV. Helium was used as the carrier gas at a flow rate at which myristic acid elutes at approximately 18 min. The quadrupole was set at 150°C and the GC-MS interface was set at 285°C. The oven program was set at 60°C for 1 min, with an increasing rate of 10°C/min until 320°C. Bake-out was set at 320°C for 10 min. Data acquisition was performed using scan mode (50-700 *m/z*). GC-MS metabolites and fragments are shown below:

| *Metabolite* | *Quantifier Ion* | *Qualifier Ion* | *Retention Time (min)* |
| --- | --- | --- | --- |
| Aspartate | 418 | 390 | 18.66 |
| Glutamine | 431 | 329 | 20.98 |

All metabolites were previously validated using authentic standards (Sigma) to confirm retention times and mass spectra. Data analysis was performed using Agilent ChemStation and Mass Hunter Quant software. The steady state level was determined for each metabolite and normalized to the peak intensity of myristic acid-D_27_ and cell number.

For LC-MS data acquisition, cells were centrifuged 7 min at 335 x g and washed three times with ice-cold ammonium formate (150 mM, pH 7.4) solution. Following centrifugation, cells were quenched on dry ice following the addition of 380 µL of 50/50% HPLC-grade methanol/water and 220 µL acetonitrile (both at -20°C). Cells were lysed by bead beating for 2 min at 30 Hz (Eppendorf Tissue-Lyser) using six 1.4 mm ceramic beads. Cellular lysate was partitioned into aqueous and lipid layers following the addition of 600 µL ice-cold dichloromethane and 300 µL cold water. Samples were incubated on ice for 10 min. The mixture was centrifuged at 4000 rpm for 10 min at 1°C. The aqueous layer was transferred to pre-cooled tubes and dried in a cold (-4°C) vacuum centrifuge (LabConco) overnight. The dried samples are dissolved in 50 µL UPLC-grade water and spun down at 13,000 rpm for 10 min at 1°C. Supernatants were transferred to LC-MS vials containing 200 µL Teflon inserts. A volume of 5µL was injected on an Agilent 6530 quadrupole time of flight (QTOF) mass spectrometer. The chromatography was performed using a 1290 Infinity ultra-performance liquid chromatography system (Agilent), which includes a vacuum degasser, autosampler, and a binary pump. The source-gas temperature was set at 150°C and the flow was set at 13 L/min. The nebulizer pressure was set at 45 psi, while the capillary voltage was set at 2000 V. Eluents were detected using negative ionization. The resolution of the metabolites was performed using a Zorbax Extend C18 column (1.8 μm, 2.1×150 mm^2^ with guard column 1.8 μm, 2.1×5 mm^2^, Agilent). The chromatographic gradient was initiated at 100% mobile phase A (97% water, 3% methanol, 10 mM tributylamine, 15 mM acetic acid, 5 μM medronic acid) for 2.5 min, followed with a 5 min gradient to 20% mobile phase B (methanol, 10 mM tributylamine, 15 mM acetic acid, 5 μM medronic acid), a 5 min to 5.5 min gradient to 45% mobile phase C (90% ACN), and a 7 min gradient to 99% mobile phase B at a flow rate of 0.25 mL/min. This was followed to 100% mobile phase B for a 4 min hold time. The restoration of the column was performed by backwashing with 99% mobile phase C for 3 min at flow rate of 0.25 mL/min. The flow rate was increased to 0.8 mL/min for 0.5 min and a 3.85 min hold, followed by a decreased rate of 0.6 mL/min for 0.15 min. Re-equilibration of the column was achieved at 100% mobile phase A for 0.75 min at a flow rate of 0.4 mL/min and held for 7.65 min. Prior to the subsequent injection, the flow was set to forward flow at 0.25 mL/min and the column temperature was set at 35°C. Transitions for quantifier and qualifier ions are shown below:

| *Metabolite* | *Precursor* | *Product* | *Fragmentor* | *Collision Energy* | *Polarity* | *Retention Time (min)* |
| --- | --- | --- | --- | --- | --- | --- |
| Dihydroorotate | 157 | 113 | 116 | 5 | Negative | 6.995 |
|  |  | 42.2 | 116 | 30 | Negative |  |
| Orotate | 155 | 40 | 68 | 44 | Negative | 8.885 |
|  |  | 42.1 | 68 | 26 | Negative |  |
| UMP | 323 | 79.1 | 126 | 48 | Negative | 10.49 |
|  |  | 97 | 126 | 24 | Negative |  |
| UDP | 403 | 79 | 108 | 48 | Negative | 14.698 |
|  |  | 158.9 | 108 | 28 | Negative |  |
| UTP | 483 | 158.9 | 146 | 36 | Negative | 16.995 |
|  |  | 385 | 146 | 20 | Negative |  |
| CTP | 482 | 79 | 142 | 40 | Negative | 16.817 |
|  |  | 158.8 | 142 | 40 | Negative |  |

Authentic metabolite standards (Sigma) were injected throughout the run to confirm retention times and mass spectra. All LC-MS solvents were purchased from Fisher (Ottawa, ON, Canada). External calibration curves (prepared in water) were run to elucidate relative concentrations. As no corrections were made for ion suppression effects, metabolite levels are presented as relative to the calibration curves and are not considered as absolute concentrations. Data were analyzed using Profinder software (Agilent). The steady state level was determined for each metabolite and normalized to cell number.

Metabolite levels from GC-MS and LC-MS data were depicted as fold change of treated (rapamycin or Torin1) cells relative to vehicle (DMSO) treated cells at each respective time point. Visualization of fold changes was carried out with the Morpheus software package (https://software.broadinstitute.org/morpheus).

#### Immunoblot and immunoprecipitation

Cells were counted on a Z2 Coulter counter and centrifuged for 7 min at 400 x g. Cell pellets were washed in 1 mL PBS, resuspended in lysis buffer (20 mM Tris-HCl pH 8.0, 137 mM NaCl, 2 mM EDTA & 1% v/v Triton X-100) with 100X protease inhibitor cocktail (MedChemExpress Cat.#HY-K0010) and phosphatase inhibitors (custom-made 200X solution of 200 mM Na-Orthovanadate and 200 mM Na-Molybdate dissolved in water and 200X solution of 1 mM Cantharidin dissolved in DMSO) where relevant, using a ratio of 1 mL per 20 million cells and lysed for 20 min on ice. After clearing extracts at 21,000 x g for 20 min at 4°C, extracts were mixed with 5X SDS-sample buffer (250 mM Tris-HCl pH 6.8, 50% w/v SDS, 50% v/v glycerol, 0.25% w/v bromophenol blue & DTT 0.5 M), heated for 5 min at 95°C, and 20 µL loaded on SDS-PAGE (BioRad). For high molecular weight immunoblots to detect POLR2A, we used Tris-Acetate NuPAGE 3 to 8% pre-cast acrylamide gels (Thermo). Gels were transferred to PVDF membrane (BioRad) at 100V for 90 min, then membrane was blocked for 15 min in 5% milk TBST (20 mM Tris, 150 mM NaCl, 0.1% Tween-20) and hybridized overnight at 4°C with primary antibodies dissolved in 1% milk TBST. Membranes were washed 3 times 10 min with TBST, incubated with a 1:5000 dilution of matched HRP-conjugated secondary antibody (Jackson laboratories) dissolved in 1% milk TBST for 1 h at RT, washed three times and fluorescence detected on a ChemiDoc (BioRad) using UltraScence Western substrate (FroggaBio).

For immunoprecipitation, we modified plasmid pCCL-c-MNDU3-ires-GFP-PGK-NEO (Guy Sauvageau, IRIC) by EcoRI-XhoI digestion, recovered the plasmid backbone by gel extraction, and ligated the backbone with phosphorylated and annealed oligos (Triple FLAG FWD for GA into plasmid 2789 and Triple FLAG REV for GA into plasmid 2789) to recreate an EcoRI cloning site downstream of an N-terminal 3XFLAG in a new plasmid, now called pCCL-c-MNDU3-3XFLAG-PGK-NEO. PABIR1 was amplified by PCR using FAM122A cDNA (NP_612206.4, a gift from Anne-Claude Gingras, Lunenfeld Tanenbaum Research Institute, Toronto) and proof-reading DNA polymerase KAPA HiFi HotStart and then cloned into EcoRI-digested pCCL-c-MNDU3-3XFLAG-PGK-NEO to generate plasmid pCCL-c-MNDU3-3XFLAG-PABIR1-PGK-Neo.

The S37A mutation was introduced into the cDNA inserted in pUC57 through site-directed mutagenesis using oligos bearing the S37A mutation by PCR-amplifying the whole plasmid and ligation by self-Gibson assembly. S37A was then cloned into pCCL-c-MNDU3-3XFLAG-PABIR1-PGK-Neo. Lentiviral particles were produced from these plasmids and used to transduce wild type NALM-6 cells, followed by selection for 8 days with 800 µg/mL G-418. The resulting cells stably expressed exogenous FLAG-tagged PABIR1. 35 mL of culture at a density of 800,000 cells per mL were pelleted for 7 min at 400 x g at 4°C, washed with 1 mL PBS1X, lysed in 1 mL lysis buffer as above in the presence of protease inhibitors for 20 min on ice and cleared by centrifugation at 21,000 x g for 20 min. Anti-FLAG magnetic beads (Sigma) were used at 20 µL of 50% slurry per condition, pre-washed with 200 µL TBS and concentrated with a DynaMag 2 magnetic tray (Invitrogen). Lysates were incubated with beads overnight at 4°C with gentle rotation. After five 1 mL washes with cold TBS1X for 5 min each, immunopurified proteins were eluted with 50 µL 100 ng/µL FLAG peptide (Sigma) dissolved in TBS with rotation for 1 h. Beads were recovered on the magnetic tray and supernatants mixed with 5X sample buffer before processing for immunoblots as described above.

#### Unbiased selection of genes implicated in the DDR

To allow clustering of compounds based on specific DNA damage response (DDR) mechanisms, including removal of general signatures due to apoptosis and other ancillary processes that would otherwise drive clustering, we generated an unbiased list of 126 DDR-associated genes based on the enrichment of hits in screens of compounds with known DNA damaging agents. We began with 18 compounds with reported DDR-related mechanisms (Table S1B), then removed the following: screens with fewer than 5 hits (0.35 µM cisplatin, 40 µM VE-822, 10 µM RS-1, 40 nM CHIR-124; all likely screened at too low concentration); 10 µM B02 which did not cluster with other DDR compound screens based on Jaccard index similarity score; 5 µM bleomycin because the apoptosis-related signature was too dominant; 0.32 µM MK-1775 because the G2/M checkpoint is not uniquely sensitive to DNA damage. We then added additional screens that had strong similarity to DDR compound screens (15 µM genistein, 5 µM menadione, 1.6 µM nutlin 3A, 16 µM pterostilbene, 16 µM resveratrol, 100 µM hydroxyurea, 5

µM dorsomorphin, 20 µM zebularine, 2 µM 5-azacytidine, 2 µM 6-thio-deoxyguanosine, 30 nM trimetrexate, 5 nM aminopterin, 10 nM methotrexate) resulting in a list of 25 compounds (Figure 6A). To select sentinel DDR genes, we considered for each gene the top 5 screens (for all screens that generated 5 or more hits) for which the gene had the highest CRANKS scores with an absolute value of at least 1.8. If 3 or more or the top 5 lowest scores (SSLs), top 5 highest scores (rescues) or top 5 scores ranked by absolute value were within the list of 25 representative DDR screens, we designated the gene as DDR-associated, resulting in a list of 126 genes (Figures 6A and S10A; Table S17). Other selected DDR genes were re-included in Figure S10A (marked with asterisks) if these were hits with many DNA damaging agents in addition to other unrelated compounds (e.g., TP53 and apoptosis genes) or strong hits with only one or two DDR compounds (e.g., SMC5 which was a hit for the crosslinker diepoxybutane only).

#### Comparison of DDR datasets

We identified 9 common compound screens in our dataset in NALM-6 cells and the Olivieri *et al.* dataset generated with TP53-/- hTERT immortalized RPE-1 cells^21^ (bleomycin, camptothecin, cisplatin, doxorubicin, etoposide, hydroxyurea, MLN4924, olaparib, pyridostatin; see Table S13)and converted deprecated gene symbols from either dataset to the newest symbols based on HGNC (retrieved February 2020). We defined hits in the Olivieri *et al.* dataset by transforming the published Z-scores into p-values and calculating FDR values separately for SSLs and rescues for each screen, resulting in a total number of CGIs very close to the published data.^21^ For compounds screened more than once in the same dataset, we used the screen (i.e., dose) with the most hits. We then plotted the gene scores in one dataset versus the other, only showing significant hits in either dataset and calculated the Pearson correlation coefficient for each compound considering the same subset of genes. We counted the fraction of hits that had a score in the same direction (i.e., SSL or rescue, less than or greater than 0, respectively) in the other dataset.

In order to compare our complete set of DDR-related hits to the Olivieri et al. dataset,^21^ we used the manually curated list of 25 screens from Figure 6A, as well as hits from our 0.2 µM MLN4924 screen and 5 µM bleomycin screen (i.e., since these two compounds are part of the Oliveri et al. dataset), compiled all unique gene hits across this dataset (900 genes, 1139 rescue and 400 SSL interactions) and across the Oliveri et al. dataset (898 genes, 64 rescue and 2317 SSL interactions). Comparison of the hit gene lists revealed an overlap of 217 hit genes. Over-represented GO biological process terms in each gene list were used to determine functional overlap between the lists and p-values of the most significant enrichments based on Fisher’s exact test (Figure S11B).

In order to benchmark our dataset as a predictor of compound MOA, we calculated all pairwise Jaccard similarities for screens from the Olivieri et al. dataset (FDR<0.05) and screens in our dataset (FDR<0.05 or absolute CRANKS score >2 threshold). We defined a set of representative DDR screens based on the list of compounds used in Figure 6A but without zebularine and 6-thiodeoxyguanosine (which did not score as similar enough to other DDR-related screens) and with MLN4924 (as part of the Olivieri et al. dataset) and all other screens with the same compounds at different doses. As a control set, we used all non-DDR screens in our dataset. For each Olivieri et al. screen and each of our non-DDR screens, we ordered all 399 screens (398 for our screens to exclude self comparisons) from highest to lowest similarity and identified the lowest ranked reference DDR screen (Figure S11A).

#### R-loop detection and cell cycle analysis

Plasmid HB-GFP (a gift from Karim Mekhail, University of Toronto; originally from Andrés Aguillera, University of Sevilla), encoding the HB domain of human RNAse H1 cloned into pEGFPC1,^146^ was used as a template to PCR-amplify the HB-GFP coding region using oligos FWD HB-GFP and REV HB-GFP. The PCR product or a control eGFP PCR product (generated with oligos FWD HB-GFP and REV eGFP) were cloned by Gibson assembly into EcoRI-digested pCCL-m-MNDU3-3XFLAG-PGK-Neo, resulting in plasmids pCCL-m-MNDU3-3XFLAG-HB-GFP-PGK-Neo and pCCL-m-MNDU3-3XFLAG-eGFP-PGK-Neo. After lentiviral particle production, we transduced either wild type NALM-6 or PABIR1 knockout clone 3B03 #11 and selected cells with 800 mg/mL G-418 over 8 days to stably express either HB-GFP or eGFP alone. Both cell line pools were greater than 95% GFP positive (Figure S11E), with eGFP alone being more intense, yet became devoid of signal once permeabilized with Triton X-100. Cells at 300,000 cells per mL were treated with inhibitors at the indicated concentrations for 18 or 24 h and fixed and permeabilized the next day at a density of 800,000 cells per mL. To assess temporal dynamics of responses, cells were seeded at 100,000 cells per mL, and, upon inhibitor treatment, 20% of the culture fixed and permeabilized each day for 4 days. To measure signal specific to HB-GFP bound to R-loops, we permeabilized cells with Triton X-100 as described.^146^ Cold PFM buffer (PBS supplied with 1% FBS and MCE Protease inhibitor at 0.4X) was used to resuspend 2.5 mL of cell culture pellet in 500 μL followed by gentle addition of 500 μL PFM with Triton X-100 (final concentration of 0.05% v/v), mixing for 5 sec, and incubation for 4 min on ice. Cells were then fixed by gentle addition 9 mL of 70% ethanol with concurrent vortexing, and then kept in the dark a 4°C until analysis by FACS. Cells were recovered by 500 x g centrifugation for 10 min, resuspended in 1.5 mL cold FACS buffer (2% FBS, 2 mM EDTA in PBS), centrifuged 2000 x g for 5 min, resuspended in 1 mL cold FACS buffer and transferred to FACS tubes through filter mesh. For cell cycle analysis, 2 million cells were recovered by centrifugation at 400 x g for 7 min in 15 mL tubes, followed by resuspension directly in 1 mL PBS prior to addition of 9 mL of 70% ethanol while swirling tubes. For cell cycle distribution, cells were washed once with 1.5 mL FACS buffer, resuspended in 1 mL FACS PI staining solution (100 μL 10% v/v Trinton-X100, 200 μL propidium iodide 1 mg/mL, 200 μL RNAse A 10 mg/mL and 9.5 mL PBS) and swirled for 30 min in a 37°C incubator before transfer in FACS tubes through filter mesh. Fluorescence was read on a BD Fortessa using 480 nM excitation (with filters 530/30) and 561 nM excitation (with filters 610/20) using Diva software, followed by data analysis with FlowJo X software.

#### Chemical synergy prediction and validation

An initial supervised approach to synergy prediction was based on compound direct targets, as obtained from the Drug Repurposing Hub^27^ or the primary literature. We selected pairs of compounds where a known drug target gene was SSL in a screen with a different compound or conversely, when a known activated target gene was a rescue hit in a screen with a different compound. These predictions, as well as additional predicted synergies based on screen similarity to a vemurafenib screen, were validated using 3-day viability assays in 384-well plate format (see above). We tested 5 x 5 dose combinations (i.e., 6 x 6 matrices including the zero dose for each), typically with 2-fold concentration increments and a median dose corresponding to the IC30 value used in screens. Each dose combination and single compound controls were run in duplicate, randomizing along with 36 or more control wells within each plate. The strategy for positioning treated and control wells across the plate varied between rounds of experiments. In earlier experiments, replicate and similar treatments were positioned next to each other, while in later experiments, we randomized the position of each combination to avoid biases caused by positioning along the plate. In most cases, we embedded a grid of control wells and normalized each treated well to the average of control wells within a 3-row and 3-column radius. For each dose combination, we calculated the Bliss score^153^ in 8 possible ways, using the 2 replicate single-compound assays for each compound and the 2 replicate combination assays, and used only the most conservative Bliss score, i.e. that with a log_2_ value closest to zero. In this way, a non-zero Bliss score indicated that the range of observed viabilities from the 2 replicates at a given dose combination did not overlap any of the 4 possible viability values expected from Bliss independence, and the smallest distance between the two ranges defined the reported Bliss interaction score. The largest absolute Bliss score obtained over the 5 x 5 dose combination matrix was reported for each compound combination (Figure 7B). Each combination was tested in this manner in two independent replicates. 9 randomly selected compound pairs were tested in the same manner to establish a baseline of Bliss scores in the absence of a predicted interaction. Monensin and vemurafenib combinations were also tested using the same approach in 3 independent replicates in NALM-6 and in one replicate in 8 melanoma cell lines (Figure S12A).

A supervised machine learning-based approach began with 15 compounds (nutlin-3A, fluvastatin, betulinic acid, MK-1775, etoposide, pterostilbene, dexamethasone, AICAR, tozasertib, apcin, bortezomib, BI2536, MLN4924, vinblastine, nifuroxazide) selected based on the number of gene hits and diversity of CGI profiles. A dose-response analysis was re-run for each drug and the estimated IC30 was used for single-dose compound combinations. Cell viability after 3 days of exposure was measured in quadruplicate wells split over two plates. The readout from each well was normalized to control wells from the same row and readouts from the first column (corresponding to the edge of the plate) were adjusted by the mean difference between controls on the edge of a plate and other controls (15% lower proliferation on the edge). Viability was defined as the relative proliferation compared to untreated controls. For each treatment, the mean viability across the four replicates was used. The Bliss model expectation corresponded to the product of the relative viability of the two single-compound treatments^153^. The log_2_ ratio of the observed relative proliferation over the expected defined the Bliss interaction scores used to train and test the model. For each target compound, a linear regression model was trained to predict the drug interaction scores of other compounds with the target compound based on each CGI profile. The predicted drug interaction score was the sum of the products of the CRANKS scores with an absolute value ≥1 and the respective gene weights fit by the model. Initial gene weights were set to zero and incrementally adjusted according to the following formula:

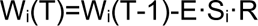

where W_i_(T) is the weight at time T for gene i, W_i_(T-1) is the previous weight, S_i_ is the CRANKS score of gene i, E (error) is the difference between the measured drug interaction score and the model prediction, and R is the learning rate, initially set to 2x10^-4^ and lowered by increments of 0.5% each time the algorithm processes all screens, or epoch. The algorithm was run to convergence, defined as a change in the error of less than 0.01% between two epochs. Leave-one-out internal cross-validation revealed significant predictive power for 14 out of 15 compounds. For these 14 compounds, we predicted Bliss interaction scores with all compounds which had been screened at the time (up to round 7) with ≥5 hits and selected both predicted synergistic and antagonistic compounds, excluding those which were deemed chemically (Tanimoto similarity ≥ 0.8) or mechanistically similar (either by reported MOA in Table S1B or previously demonstrated based on this dataset, e.g., hydroxyurea, resveratrol and pterostilbene^179^) to a compound in the training set.

To validate the machine learning-based predictions, we ran 3-day viability assays in a single 384-well plate in quadruplicates for all predicted interacting compound pairs at the IC30 dose as well as each single compound at the same dose. Luminescence values were normalized to the average of all 0.1% DMSO control wells from the plate. For each compound combination, we used the individual replicate wells to calculate 4 independent Bliss scores, combining a single-compound well for each compound with one combination-treated well in no particular order. Then, for each pair of compounds predicted to interact differently from each other when combined with the same compound from the training set (one predicted antagonist and one predicted synergist), we calculated all 16 pair-wise Bliss score differences (antagonist score – synergist score). To optimize the signal-to-noise ratio, we only considered pairs where the observed Bliss score difference had an absolute value of at least 0.3 and the predicted Bliss score difference was ≥20 in relative rank out of the 198 compounds considered. A positive average score difference was observed from most compounds, confirming that the model can effectively predict relative synergies and/or antagonism (Figure 7C).

#### Scoring alternative exons and hypothetical genes with CRANKS

Only exons that are part of RefSeq (i.e., core) genes were scored. CRANKS scores for alternative exons were calculated using only guides that were not used in gene scoring to avoid spurious correlations. The same minimal read counts per guide were used as for the regular scoring. The minimal number of guides per exon passing the read count threshold was set to two. Hypothetical genes from the extended library were scored in parallel for efficiency and guides used for these were restricted to guides with a single match across the genome and not used in RefSeq gene or exon scoring. RefSeq genes were also scored in parallel to better estimate FDR values but their scores were not output during these CRANKS runs. In total, 755 exon-level chemogenomic interactions with RefSeq exons were identified at an FDR of <0.05, involving 440 different exons (Table S18A). The interaction rate is lower than for RefSeq genes in part because on average only 3 sgRNAs target each alternative exon, meaning that a greater effect size is required to achieve statistical significance. Thus, to reduce the effects of false negatives for downstream analyses, we considered exons with an absolute CRANKS score ≥ 2.5 as chemical-exon interactions, only considering interactions where the parent gene also interacts with the compound in the same screen (FDR<0.05). In cases where the exon score was of opposite sign to that of the gene or of absolute value <0.5, the interaction was considered as not shared. Exon scores between ≥0.5 and ≤2.5 in the same direction as the parent gene were excluded from the analysis. We thus identified 629 exons with no shared interactions with the parent gene, 661 exons that share all interactions and 550 exons that share a fraction of interactions (Table S18B).

#### Tandem mass spectra for protein-coding hypothetical genes

Only 64 CGIs for 46 hypothetical genes were identified at an FDR<0.05 across all 399 experimental screens (Table S19A). The low number of significant CGIs identified can be partially attributed to having combined their scoring with that of alternative exons targeted by only ∼3 sgRNAs each, resulting in larger average top-ranking p-values and thus larger FDR values for hypothetical genes as well. We thus defined a set of 217 hypothetical genes with likely CGIs by including those with two or more CGI uncorrected p-values below 0.001. As a control set, we used hypothetical genes without any uncorrected p-values below 0.002, absolute CRANKS scores of 2.5 or greater, or within the top 500 most essential, as defined previously.^24^ All of these potential CGIs are included in Table S19A. We further filtered out from both sets genes with fewer than 4 sgRNAs, each with a single cut site, and those with no unique tryptic peptides. Using mass spectral counts obtained previously from reanalyzing public tandem mass spectrometry data,^24^ we found that hypothetical genes with likely CGIs had significantly more uniquely-matching spectra than those in the control set (p=3.8e-6, U test, Figure S12E; Table S19B).

#### CRANKS2 and evaluation of comparative predictive power

CRANKS2 models essentiality along two dimensions instead of one, the first being the level of essentiality and the second being the relative rate at which essentiality-driven sgRNA depletion occurs during the 8-day chemogenomic screens. The former is estimated from the gene RANKS sgRNA depletion score after 15 days of population growth (as it is for CRANKS) and the latter from the Pearson correlation coefficient (PCC) between the gene CRANKS scores and measured population doublings across all chemogenomic screens (Table S20). As expected, the CRANKS scores of essential genes whose phenotypes largely fail to manifest before the onset of the screen tend to be inversely correlated with population doublings (i.e. CRANKS under-corrects for their essentiality), while essential gene phenotypes that largely manifest before the screen are generally over-corrected by CRANKS (Figure S12F). Control sgRNAs from which sgRNA p-values are calculated are the same as for CRANKS for genes that are not part of the 5000 most essential genes. For genes within the top 5000 most essential, control sgRNAs are those targeting genes that correspond to the following criteria: (1) is within the 500 genes immediately more essential than the gene itself, (2) has an absolute difference in score-vs-doublings PCC of no more than 0.15 or (3) 100 in relative rank. This strict selection process sometimes results in a control set that is too small to make meaningful inference. For genes with fewer than 500 control sgRNAs corresponding to the above criteria, we relaxed the first criterion to include genes with RANKS scores with an absolute difference of ≤1, keeping the second two selection criteria unchanged. Genes with fewer than 50 control sgRNAs corresponding to the second set of criteria were discarded.

The DepMap CRISPR screen dataset^6^ was obtained in May of 2020. While all of these screen data were produced at the Broad institute (769 screens), later versions include screens from the Sanger institute, which due to the greater heterogeneity of the combined dataset, performed worse on the same metrics despite the larger number of screens. Pearson correlation was performed on all gene pairs for each dataset and gene pairs were ordered from highest to lowest by correlation score. Only screens with ≥10 hits in the CGI dataset were used to calculate correlations (210 screens). Protein-protein interaction (PPI) data were retrieved from BioGRID (2025-11-06 release, version 5.0.251)^5^ and filtered to include only interactions supported by two or more primary sources. Biological process annotations were retrieved from the Gene Ontology (GO) website (version 2023-07-29)^180^. Gene pairs were considered to share a GO term provided the term was annotated no more than 25 genes. Fold-enrichment was calculated by comparing the overlap to that of the median of a series of randomized correlation networks, in order to control for biases in the coverage of hub genes with more GO annotations or PPIs. Only correlated pairs where both genes were represented in the validation network were considered, skipping other pairs until the displayed network size was reached. Each gene in the network was added to a sampling pool of genes once for every edge. The randomized networks, which preserved node degree as much as possible, were then generated by sampling pairs of genes from this pool (with replacement) until the same network size was reached. For the top 1,000 and 2,500 pairs, 1,000 randomized networks were generated, for the top 5,000-25,000, 500 networks were generated and for 50,000-100,000, 50 networks were generated. The overlap between the PPI or GO network and each randomized network was calculated and the median value used to calculate fold-enrichment. The DepMap dataset unexpectedly produced greater fold-enrichments when considering more correlated pairs, which is likely because, being based on gene essentiality screening, its top correlated pairs are so heavily biased toward hub proteins that the same genes are highly likely to be interacting or share functions by chance, as the randomization procedure is meant to control for.

#### Comparison of the human CGI network with global yeast genetic interaction network

We downloaded the list of yeast gene pair correlation coefficients from TheCellMap.org ^181^, based on genome-wide genetic interaction profiles (Costanzo et al. 2016). Given there are two coefficients per pair in this data, we averaged the two coefficients. To control noise, we filtered down the network to the 7,260 pairs in which either of the two coefficients had an absolute value above 0.3. We calculated the mean correlation coefficient between pairs of GO terms as the sum of the correlation coefficients implicating one gene from each term divided by the total number of possible gene pairs. The same metric was used for our CGI dataset, using the top 10,000 gene pairs instead of the 0.3 cut-off. Pairs of GO terms with a mean correlation score in both species and at least 10 genes in the smallest term in both species were transformed into quantiles and plotted against each other, once including all pairs (Figure S13B) and once excluding pairs of terms with significant overlap in gene content, defined by a Jaccard index of ≥0.05 (Figure S13C).

#### Gene-gene correlation and annotation of the gene profile similarity network

Using CRANKS2 scores from screens with at least 10 significant hits (FDR<0.05), we calculated the Pearson correlation coefficient (PCC) between all pairs of RefSeq genes, with no more than 30 missing scores due to insufficient sgRNA read coverage. For the gene profile similarity network, we selected the 10,000 strongest correlations (PCC ≥ 0.3882). We then calculated the network distance between all gene pairs (as an unweighted network) and for each gene neighborhood of 2-edge or 3-edge radius (i.e., all genes within 2 or 3 connections from the central gene), we calculated GO term enrichment using Fisher’s exact test, comparing the numbers of annotated genes in each neighborhood to those of all other genes in the GO database (version 2019-03-19).^180^ For each neighborhood with one or more enrichments with FDR<0.001, we selected one of the three most significantly enriched GO terms to display, prioritizing terms with the most genes and excluding highly overlapping terms. The resulting network (Figure 7G) was visualized with Cytoscape 3.6.1 (https://cytoscape.org/) using the default layout with minor manual adjustments.^182^ Alternatively, the correlation network was visualized for the top 15,000 gene pairs (PCC ≥ 0.362) as an edge-weighted spring-embedded network layout (Cytoscape 3.10.4) only for nodes that formed the main network (Fig S13A). The layout was imported to the RISK software tool^160^ and clustered by the Markov Clustering algorithm with Jaccard distance as the linkage method (random seed = 888). Clusters with statistically significant enriched terms from PAN-GO slim v19.0.^183^ were defined by hypergeometric test (FDR < 0.05) automatically by RISK.

## Acknowledgements

We thank Sam Weiss, Artee Luchtman, Cheryl Arrowsmith, the Structural Genomics Consortium, Jerry Pelletier, and Daniela Rotin for suggesting and/or providing compounds for screens; Susan Moore and Manon Lord for technical support; the Genomics, High-Throughout Screening, Flow Cytometry, and Medicinal Chemistry platforms at IRIC for technical services; D. Avizonis of the Metabolomics Innovation Resource (MIR) at McGill University for supervision of metabolic analyses; and Daniel Durocher, Anne-Claude Gingras, Jason Moffat, Michael Costanzo, Yahya Benslimane, Charlie Boone, and Brenda Andrews for helpful discussions. This study was supported by grants from: the Canadian Institutes of Health Research (FDN-167277 to Mike Tyers, PJT190250 to Sylvie Mader, PJT190250 to Sylvain Meloche, PJT152920 to B.K., PJT-451236 to I.T., FDN–167265 to D.D., 163051 to Mike Tyers and L.H.); the National Institutes of Health (RO1 NCI80728 and RO1 NCI98571 to K.L.B.B.; R35 GM127032 to R.W.K.; R01 OD010929 to Kara Dolinski and Mike Tyers); the Leukemia and Lymphoma Society USA (TRP to K.L.B.B.); the Medical Research Council (MC_UU_00035/13 to E.E.P.); the Keck Foundation for Medical Research to J.D.A. and Mike Tyers; Stand Up To Cancer Canada Cancer Stem Cell Dream Team Research Funding (SU2C-AACR-DT-19–15 to P.B.D., Samuel Weiss and Mike Tyers; provided by the Government of Canada through Genome Canada and the Canadian Institutes of Health Research, with supplementary support from the Ontario Institute for Cancer Research through funding provided by the Government of Ontario; Stand Up to Cancer is a program of the Entertainment Industry Foundation administered by the American Association for Cancer Research); Genome Canada and Genome Quebec (LSARP4524 and LSARP13528 to G.S.; Centre for Advanced Proteomic and Chemogenomic Analyses GTP to P.T. and Mike Tyers); the Institute for Research in Immunology and Cancer Philanthropic Foundation and La Fondation Famille Godin to A.M., B.W., Mike Tyers; the Natural Sciences and Engineering Research Council (RGPIN-2022-04206 to V.A.); the McLaughlin Centre Accelerator (MC-2023-07 and MC-2024-08-3 to LH. and M.T); the Terry Fox Foundation (TFF-242122 to I.T.); the Canada Foundation for Innovation; Canada Research Chair awards to K.L.B.B., D.D., F.S., Sylvie Mader, Sylvain Meloche, Marc Therrien, G.S., I.T., and Mike Tyers; and Canadian Institutes of Health Research and IVADO postdoctoral fellowships to J.C.-H.

## Author contributions

Conceptualization J.C-H., T.B., and Mike Tyers; methodology J.C-H., T.B., and C.H.; software J.C-H., A.C.-A., H.S., C.S., B.-J.B.; validation J.C-H., T.B., C.H., and D.J.S.-C.; formal analysis J.C-H.; investigation J.C-H., T.B., C.H., D.J.S.-C., M.S.-O., D.P., K.N., J.P., Sandhya Manohar, C.S.-D., L.Z., Shannon McLaughlan; resources A.M.v.d.S., H.L., K.L.B.B., B.R., D.D., F.S., A.V., Sylvie Mader, Sylvain Meloche, Marc Therrien, P.B.D., E.E.P., R.W.K., J.D.A., P.P.R., G.S., T.H., A.M., L.H., B.K., V.A., I.T., Mike Tyers; data curation J.C-H., T.B., C.H., A.C-A., D.J.S.-C., M.S.-O., C.S., B.-J.B., Mike Tyers; writing - original draft: J.C-H., T.B., B.K., V.A., I.T., Mike Tyers; writing - review & editing J.C-H., T.B., M.S.-O., R.P., K.L.B.B., B.W., R.W.K., G.S., L.H., B.K., V.A., I.T., Mike Tyers; visualization J.C-H., T.B., M.S.-O., D.P., K.N., J.P., H.S., R.P., Mike Tyers; supervision: J.C-H., T.B., B.K., V.A., I.T., Mike Tyers; project administration J.C-H., T.B., Mike Tyers; funding acquisition K.L.B.B., B.R., D.D., Sylvie Mader, Sylvain Meloche, Marc Therrien, P.T., B.W., P.B.D., E.E.P., R.W.K., J.D.A., P.P.R., G.S., T.H., A.M., L.H., B.K., V.A., I.T., Mike Tyers.

## Declaration of generative AI and AI-assisted technologies in the writing process

Generative AI was not used in any form to assist with writing of the manuscript. The authors take full responsibility for the content of the published article.

## Supplementary figure legends

**Figure S1.** Reproducibility and quality of chemogenomic CRISPR screens, related to Figure 1

(A) CRANKS score reproducibility at two different read depths. Shared hits are in green; hits unique to the 440 read/sgRNA screen in blue; hits unique to the 54 read/sgRNA screen in red.

(B) RANKS essentiality score reproducibility at two different minimal cell counts after dilution. Color scheme as in panel A.

(C) Relationship between mRNA expression and highest absolute CRANKS score.

(D) Relationship between mRNA expression and number of CGIs.

(E) Global heatmap of chemical-genetic dataset for all 327 screens (columns) and 3178 genes (rows) with ≥1 CGIs. Clusters described in the main text and additional CGI clusters are annotated.

**Figure S2.** TRAIL screen hit validation, related to Figure 2.

(A) NALM-6 cell line with indicated gene knockouts (2 independent guides per gene) or control non-targeting sgRNAs were treated with TRAIL at the indicated concentrations and then assayed for viability using CellTiter-Glo reagent.

(B) Same genes and assays as in panel A but in Jurkat T-cell line.

**Figure S3.** Chemogenomic profiles of selected drugs to illustrate MOA, related to Figure 2. Significant rescue or SSL hits (FDR < 0.05) in single CGI profiles are colored blue or magenta, respectively. Hits shared by both screens in comparison plots are colored in red; single-screen hits are colored in black. Notable screen hits discussed in the text are circled in red.

(A) Phorbol 12-myristate 13-acetate (PMA, 570 nM) CGI profile (left). Schematic of screen hits are indicated on PMA network diagram (right).

(B) WEE1 inhibitor MK-1775 (320 nM) versus erlotinib (10 µM) CGI profile comparison. EIF2AK4 and GCN1L1 were top rescues in both screens, consistent with off-target activation of EIF2AK4. The direct target WEE1 was a strong SSL in the screen. CDK2 and its cyclin CCNA were strong rescues, consistent with the role of WEE1 in restraining CDK2 activity.

(C) DNA damaging agent temozolomide (200 µM) versus DYRK3A inhibitor GSK626616 (14 µM) CGI profile comparison. EIF2AK4 was the top rescue in both screens, consistent with off-target activation of EIF2AK4.

(D) Structures of erlotinib, MK-1775, GSK626616 and temozolomide.

(E) GSK3 inhibitor LY2090314 (3 nM) versus GSK99021 (3.81 µM) CGI profile comparison (left). Schematic of screen hits, including adenosine kinase ADK, are indicated on GSK3 network diagram (right).

**Figure S4.** Chemogenomic profiles of selected drugs to illustrate MOA, related to Figure 2. Significant rescue or SSL hits (FDR < 0.05) in single CGI profiles are colored blue or magenta, respectively. Hits shared by both screens in comparison plots are colored in red; single-screen hits are colored in black. Notable screen hits discussed in the text are circled in red.

(A) CDK4/6 inhibitor palbociclib (1 µM) CGI profile. AMBRA1 and RB1 were top rescues consistent with known roles as an E3 ubiquitin ligase for CCND3 (cyclin D3), a CDK4/6 activator, and the main downstream cell cycle target of CDK4/6, respectively. Unexpectedly, the activator CCND3 was also a rescue in the screen (see panel B).

(B) Allelic series of CCND3 indels explains rescue in the palbociclib screen. Log_2_ fold-changes for each CCND3-targeting sgRNA in the palbociclib screen versus relative targeting position within the CCND3 reading frame are shown. Distal sgRNAs were enriched in the screen.

(C) PARP inhibitor rucaparib (6.5 µM) versus olaparib (4 µM) CGI profile comparison.

(D) Heavy metal arsenate (40 µM) CGI profile. Rescues by phosphate transporter SLC20A1 and phosphorylation-based signaling factors suggested that the primary mechanism of arsenate toxicity is interference with phosphate metabolism and substrate phosphorylation.

(E) Lot of suboptimal fetal bovine serum (FBS, 10%) versus nucleoside analog cytarabine (16 nM) CGI profile comparison (left). Schematic of screen hits are indicated on cytosine metabolism network diagram (right). Cytosine transporter SLC29A1 and metabolic enzyme hits suggested that the FBS was contaminated with a toxic cytosine analog from an unknown source.

(F) Proteasome inhibitor bortezomib (5 nM) CGI profile. Proteasome subunits were both SSLs (PSMB5, PSMB6, PSMB8) and rescues (PSMB9). Translation factors were also frequent rescues (EIF4A1, EIF4H, EIF4G, DDX3X).

(G) Mitochondrial complex I inhibitors rotenone (70 nM) versus phenformin (20 µM) CGI profile comparison. The cytoplasmic glutamic-oxaloacetic transaminase GOT1 was the top SSL and the NAD+ transporter SLC25A51 was the top rescue in both screens.

(H) Antiviral guanosine analog ribavirin (10 µM increased to 15 µM on day 4) CGI profile. AMP deaminase AMPD2 was a strong sensitizer due to reduction of IMP levels and ADK was a strong rescue possibly due to restoration of AMP balance.

**Figure S5.** Chemogenomic profiles of selected drugs to illustrate MOA, related to Figure 2. Significant rescue or SSL hits (FDR < 0.05) in single CGI profiles are colored blue or magenta, respectively. Hits shared by both screens in comparison plots are colored in red; single-screen hits are colored in black. Notable screen hits discussed in the text are circled in red.

(A) Hypoxia (5% O_2_) CGI profile. ADK strongly sensitizes to low oxygen levels potentially due to disruption of AMPK activation and energy metabolism imbalance.

(B) Protein kinase A activator cAMP (200 µM) CGI profile. Adenosine kinase ADK and the equilibrative nucleoside transporter SLC29A1 were strong rescues likely due to reduction in AMP pools and equilibrative efflux of cAMP respectively. AMP deaminase AMPD2 was a strong SSL likely due to an increase in AMP pools.

(C) Purine biosynthesis intermediate and AMPK activator AICAR (240 µM) CGI profile. Many genes in purine nucleotide metabolism (ADK, AK2, ADSS, NUDT2, IMPDH2), as well as methionine biosynthesis, one carbon metabolism and other metabolic processes (MTR, MTRR, MTHFD2, MMACHC) were strong rescues. Genes implicated in purine-pyrimidine nucleoside balance (UCK2, NT5C2) were SSL.

(D) AMPK activator PF-06409577 (20 µM) CGI profile. All three AMPK subunits (PRKAB1, PRKAA1, PRKAG1) were strong rescues. The AMPK activating kinase STK11 (a.k.a. LKB1) and its activator STRADA were also strong rescues.

(E) Ribonucleotide reductase inhibitor hydroxyurea (100 µM) CGI profile. Amongst many hits in the DNA damage response, translation and metabolism, the direct target RRM2 was a strong SSL.

(F) Jaccard index similarity was more robust to batch effects than Pearson correlation (PCC). Similarity was defined as the z-score relative to screen pairs from different rounds.

(G) Jaccard index similarity was more robust to population doubling effects than Pearson correlation.

(H) p97/VCP inhibitor CB-5083 (0.4 µM) versus NMS-873 (70 nM) CGI profile comparison. Whereas CB-5083 yielded many hits consistent with general functions in protein homeostasis, the NMS-873 profile suggested it is a mitochondria complex I inhibitor, with GOT1 as a strong SSL and SLC25A51 as a strong rescue (see Figure S4G for mitochondria complex I inhibitor profiles).

(I) CGI profile comparison of the active atropisomer (R)-(+)-1,1’-binaphthyl-2,2’-diamine (R-DABN, 8 µM) and inactive atropisomer (S)-(+)-1,1’-binaphthyl-2,2’-diamine (S-DABN, 8 µM). R-DABN was rescued by spindle assembly checkpoint genes (see Figure 4), as well as by the APC/C activator CDC20.

(J) CGI profile comparison of the NAE1 inhibitor MLN4924 (200 nM) and SAE1 inhibitor ML-792 (200 nM). Unique hits for MLN4924 and ML-792 reflected respective inhibitor specificity for NEDD8 versus SUMO ubiquitin-like modifier conjugation systems. MLN4924 rescues included subunits of the COP9/signalosome deneddylation complex, while SSLs included the cullin-RING ligase CUL4A subunit and the associated E2 enzyme UBE2G1. ML-792 rescues included SUMO1/2, suggesting that toxicity may arise from accumulation of excess SUMO when SAE1 is inhibited. Both inhibitors were strongly rescued by the BEND3 transcriptional co-repressor.

**Figure S6.** Features of multidrug sensitivity and resistance genes, related to Figure 3.

(A) Gene interaction degree in the CGI dataset approximates a power-law distribution.

(B) Heatmap of core apoptosis CGIs.

(C) Previously-reported small molecule-transporter pairs tend to produce CRANKS scores ≥1.

(D) SSL compound interactions with the ABCC1 transporter. Literature citations that report each compound substrate are listed as PMIDs.

(E) GPR107 CRANKS scores are anti-correlated with compound dose to predicted solubility ratio. Values above 0 predict over-saturation.

(F) FLVCR1 CRANKS scores are anti-correlated with compound dose to predicted solubility ratio.

(G) Predicted logP (hydrophobicity) versus pKa of the most basic site. Note that pKa values below 7 do not indicate predicted acidity. Labeled red dots indicate compounds in the PLD cluster.

**Figure S7.** Additional analysis of mitotic control network, related to Figure 4

(A) Scores of genes located on chromosome 7 in combined triplicate nocodazole screens according to chromosomal position. MAD1L1 is located at the distal end of the 7p arm (7p22). Genes located between MAD1L1 and the centromere generate rescues, whereas genes on the 7q arm in general do not. A notable exception is OCM2 that may have a high score because it shares sequence similarity with OCM on the 7p arm and thereby generate incidental breaks at the OCM locus. The lower zoom-in section shows that genes positioned beyond MAD1LI, i.e., between MAD1L1 and the telomere, did not generate significant scores.

(B) Schematic to illustrate that upon Cas9-mediated double-strand breaks, mis-repair can lead to chromosome arm truncation and MAD1L1 loss.

(C) Manhattan-style plot of gene scores in the aminopterin screen as a function of genomic position. On chromosome 21, loss of the SLC19A1 transporter at 21q22.3 confers aminopterin resistance. Adventitious Cas9-induced truncations on 21q confer resistance due to incidental loss of SLC19A1.

(D) The known mitochondrial complex I inhibitor rotenone has two separable signatures: a low dose signature shared with other mitochondrial complex I inhibitors and a high dose signature shared with nocodazole and other microtubule poisons. The putative p97/VCP inhibitor NMS-873 also clustered strongly with complex I inhibitors, indicating an off-target mechanism.

(E) Schematic of PIDDosome-dependent cell death network indicating hits for the actin polymerization inhibitors cytochalasin B and latrunculin B, and the aurora kinase A/B inhibitor tozasertib. Extra centrosomes present in cells following cytokinesis failure induce this specialized form of cell death.

(F) Hit list for combined cytochalasin B (5 µM), latrunculin B (10 µM), tozasertib (0.1 µM) screens (Table S1E). 33 centrosomal genes shown in blue were strong rescues across the screens.

**Figure S8.** Additional analysis of mTOR network, related to Figure 5

(A) CGI profile comparison of 3 combined active-site mTOR inhibitor screens (Torin1 80 nM, INK128 200 nM, KU-0063794 3.8 µM) versus combined triplicate independent rapamycin (1 µM, 2 µM, 10 µM) screens. Top hits are labeled, see Tables S12B and S12C for gene scores.

(B) Supplementation of cells with nucleosides partly rescues proliferation inhibition caused by rapamycin. The same data as in Figure 5C is re-plotted for uridine, cytidine and uridine + cytidine. Additional panel shows data for supplementation with a mixture of all four nucleosides (EmbryoMax).

(C) RNA-seq analysis of PPP3CA knockout (log_2_ fold change versus non-targeting guides) versus NALM-6 cells treated for 20 h with 5 µM calcineurin inhibitor FK-506 (log_2_ fold change versus DMSO).

(D) CGI profile comparison of three combined independent rapamycin screens versus two combined independent AICAR (200 µM, 240 µM) screens.

(E) CGI profile comparison of three combined independent active-site mTOR inhibitor screens (INK128 0.2 µM, Torin1 80 nM, KU-0063794 3.8 µM) versus with two combined independent AICAR screens.

(F) Schematic network of nucleotide metabolism. Hits from two combined independent AICAR screens are indicated (Table S1E). Network was adapted from https://www.kegg.jp/pathway/hsa01232.

**Figure S9.** Network schematic of cholesterol biosynthesis, related to Figure 5. Hits from two independent screens with the HMGCR inhibitors (statins) fluvastatin (2.2 µM) and cerivastatin (150 nM) were combined and overlaid on the network.

**Figure S10.** Additional analysis of DNA damage network, related to Figure 6

(A) Summary of DDR network modules and drug mechanisms revealed by CGI hits for DNA damaging agents used in this study. Genes with asterisks were hits in some DDR screens but did not meet inclusion criteria for the list of 126 unbiased DDR genes (see Figure 6A; Table S17 and Methods).

(B) Comparison of CGI profiles for the G-quadruplex stabilizer pyridostatin (4 µM) versus combined screens for three topoisomerase II inhibitors (etoposide 100nM, doxorubicin 0.02 µM, genistein 15µM). Profile similarities suggests that pyridostatin acts as a topoisomerase II inhibitor.

(C) Comparison of CGI profiles for the DHFR inhibitor trimetrexate (30 nM) versus the average scores of the DHFR inhibitors aminopterin (5 nM) and methotrexate (10 nM). Shared hits are colored in red.

(D) Comparison of CGI profiles for low (6 nM) and high (55 nM) concentrations of the PARP inhibitor talazoparib. Shared hits are colored in red.

(E) PABIR1 knockout clone cells proliferated faster than wild type NALM-6 cells (labeled with mCherry) when co-cultured for 4 days in the presence of mTOR inhibitor rapamycin (4 µM), CHEK1 inhibitor SAR-020106 (2 µM), PP2A inhibitor LB-100 (4 µM) or WEE1 inhibitor MK-1775 (400 nM).

(F) PP2A phosphatase inhibitor LB-100 (4.1 µM) CGI profile. The PP2A inhibitor PABIR1 (formerly FAM122A) was a rescue.

(G) CTD phosphorylation status of POLR2A is not altered in response to CHEK1 inhibitor (2 µM SAR-020106, 18 h) in either WT NALM-6 or a PABIR1 knockout clone.

(H) WT NALM-6 or PABIR1 knockout cells were modified to express either eGFP (left) or the RNA/DNA binding domain of human RNAse H1 fused to eGFP (HB-GFP, right). Greater than 95% of cells were labeled and expressed equal fluorescence across the two genotypes.

(I) Specific R-loop dependent retention of HB-GFP signal. Permeabilization with Triton X-100 (see Methods) caused loss of nearly all non-specific eGFP signal with specific HB-GFP signal dependent on R-loop induction after 18 h treatment with CHEK1 inhibitor SAR-020106 (2 µM), WEE1 Inhibitor MK-1775 (500 nM) or AKT1/2 inhibitor MK-2206 (2 µM).

**Figure S11.** Additional analysis of DNA damage network, related to Figure 6.

(A) Jaccard index similarities between this dataset and the Olivieri dataset^21^ predict compound mechanism of action.

(B) Gene set enrichment for DDR modules in this study and the Olivieri dataset.^21^ The size of each circle is proportional to the number of DDR hit genes annotated with the term.

(C) Relative gene expression of hits in NALM-6 versus RPE-1 cells in part explains non-overlapping significant hits between the CGI dataset and the Olivieri dataset.^21^

(D) Comparison of scores for CGIs with 9 compounds screened both in this study and in Olivieri et al. (2020).^21^

**Figure S12.** Additional global analysis of the CGI dataset, related to Figure 7.

(A) Vemurafenib and monensin synergistically inhibited proliferation in the NALM-6 and Jurkat cell lines, as well as in 7 out of 8 tested melanoma cell lines.

(B) Exons that share some or all CGIs with the parent gene tended to be more highly expressed than non-interacting exons.

(C) Exons that share some or all CGIs with the parent gene were less likely to exhibit exon skipping.

(D) Exons that share only a subset or none of the parent gene CGIs were more likely to overlap known alternatively-spliced regions. Error bars show the bootstrapped standard error based on 100 resamplings.

(E) Hypothetical genes with probable CGIs were significantly more likely to encode expressed proteins, as indicated by a greater number of uniquely-matched peptide mass spectra compared to ORFs without CGIs.

(F) Essential gene sgRNA depletion at screen onset (pre-doxycycline to post-doxycycline essentiality) predicts correlation between CRANKS scores and the number of population doublings.

(G) Examples of two genes that were either under- or over-corrected by CRANKS. PMM2 was not depleted at the start of screens and was under-corrected by CRANKS, such that its scores negatively correlated with population doublings. PSMD3 was highly depleted at the start of screens and was over-corrected by CRANKS, such that its scores positively correlated with population doublings.

**Figure S13.** Conserved overall functional architecture inferred from the human CGI dataset and a comprehensive yeast genetic interaction dataset, related to Figure 7.

(A) Alternative analysis of gene-gene correlation network. The top 15,000 gene-gene correlations (Table S15B; PCC ≥ 0.362) were used to build a network of 2,656 genes in Cytoscape. The network was annotated using the PAN-GO v19.0 GO slim biological process term set and terms for 29 significant clusters (FDR < 0.05) assigned by RISK. Only the giant cluster comprised of 2223 nodes (83.7%) and 14,731 edges (98.5%) is shown.

(B) Comparison of mean gene pair correlation between GO term pairs, including overlapping terms. Yeast data were obtained from thecellmap.org.^181^

(C) As for panel B but without overlapping GO term pairs.

## Supplementary table legends

**Table S1.** Summary of the CGI dataset.

(A) List of sequenced samples, sequencing depth, number of sgRNAs detected and population doublings.

(B) List of independent screens, compound classifications, concentrations screened, population doublings and number of chemical-gene interactions (CGIs).

(C) Compound chemical descriptors, sources and verification by analytical methods.

(D) log_2_ RNA-seq reads per million in NALM-6 cells for all genes.

(E) Complete list of CGIs detected in this study at FDR <0.05. FDR values of zero indicate <0.001.

**Table S2.** Genome-wide CRANKS score matrix for all screens in this study. Scores were corrected for artifactual chromosome 7p hits caused by chromosome arm loss (see Figure 4). Data in this table was used to build Figure S1E.

**Table S3.** List of relevant known compound-target relationships from the Drug Repurposing Hub^24^ or source PubMed ID, as indicated.

**Table S4.** Combined CRANKS scores for two independent TRAIL screens.

**Table S5.** Jaccard similarity indices between screens in the CGI dataset. Pairs used in Figure 2D are indicated by class.

**Table S6.** Compound-transporter CGI scores. Sub-threshold scores ≥1 and ≤-1, known and unknown transporter-substrate relationships from VariDT are indicated.

**Table S7.** Dose-to-solubility ratios and correlations with CRANKS scores.

(A) Compound dose-to-solubility ratios by screen.

(B) Correlation coefficients of CRANKS scores with compound dose-to-solubility ratios.

**Table S8.** Vemurafenib SSL hits and Gene Ontology annotations.

**Table S9.** Predicted basic pKa and logP for compounds represented in the screen similarity network shown in Figure 2E.

**Table S10.** Mean genome-wide CRANKS score profile of the 28 screens in the phospholipidosis cluster. Top 19 phospholipidosis SSL genes from Kumar et al (2020)^58^ are indicated.

**Table S11.** LipidTOX Red assay determinations.

(A) Cell counts and average LipidTOX levels (phospholipid accumulation) after 24 h compound exposure.

(B) Categorical classification of compounds based on assay results.

**Table S12.** Average CRANKS scores for combined screens.

(A) Average CRANKS scores for three nocodazole screens (100 nM, 200 nM, 200 nM).

(B) Average CRANKS scores for three active site mTOR inhibitors (KU-0063794 3.8 µM, INK128 0.2 µM, Torin1 80 nM).

(C) Average CRANKS scores for three rapamycin screens (1 µM, 2 µM, 10 µM).

(D) Average CRANKS score for two AICAR (240 μM) replicate screens.

(E) CRANKS scores for pyridostatin and the average of topoisomerase II inhibitor (etoposide 100 nM, doxorubicin 20 nM, genistein 15 µM) screens.

(F) CRANKS scores for three DHFR inhibitor screens (aminopterin 0.005 µM, methotrexate 0.01 µM, trimetrexate 0.03 µM).

(G) CRANKS scores for four PARP inhibitor screens (olaparib 4 µM, rucaparib 6.5 µM, talazoparib 6 nM and 55.2 nM).

(H) CRANKS scores for two CHEK1 inhibitor screens (SAR-020106 2.2 µM, CHIR-124 0.142 µM).

**Table S13.** Comparison of CGIs for 9 shared compounds in this dataset and the Olivieri et al (2020)^21^ dataset.

**Table S14.** Data for assessment of compound synergy/antagonism.

(A-D) Luminescence readings and treatments by well for synergy/antagonism results shown in Figure 7A,B.

(E) Luminescence readings and treatments by well for monensin-vemurafenib synergy validation in 8 melanoma cell lines.

(F) Observed Bliss scores (log_2_) and compound pair classifications for Figure 7B.

(G) Luminescence readings and treatments by well for target-based synergy prediction results shown in Figure 7B.

(H) Luminescence readings and treatments by well for machine learning model training results.

(I) Luminescence readings and treatments by well for machine learning model validation results shown in Figure 7C.

**Table S15.** Genome-wide CRANKS2 scores for CGI dataset and top correlated gene pairs.

(A) Genome-wide CRANKS2 score matrix for all screens in this study. Scores were corrected for artifactual chromosome 7p hits caused by chromosome arm loss (see Figure 4).

(B) Top 20,000 gene pairs with the most correlated CRANKS2 scores across the 210 screens with ≥10 hits with GO term annotations used in Figure 7G.

**Table S16.** RNA-seq data.

(A) Upper-quartile normalized log_2_ fold-changes in FK-506 (5 µM) treated cells versus DMSO and read counts by gene.

(B) Mean upper-quartile normalized log_2_ fold-changes relative to controls for PPP3CA knockout.

(C) Mean upper-quartile normalized log_2_ fold-changes relative to controls for PPP3R1 knockout.

(D) ERCC spike-in experiment read counts and relative transcription rates in PABIR1 (FAM122A) knockout clones.

**Table S17.** List of DDR-associated genes.

**Table S18.** Alternative exon data.

(A) Significant exon-level CGIs.

(B) Exon classification and validation data for Figure S12B-D.

**Table S19.** Data for hypothetical genes.

(A) Hypothetical gene CGIs with p-values <0.002 or absolute scores ≥2.5.

(B) Hypothetical gene classification and mass spectral counts.

**Table S20.** Pearson correlation coefficients of individual gene CRANKS scores with population doublings and essentiality RANKS scores at screen onset (pre-doxycycline to post-doxycycline)

