## Supplementary Information for "A Chemical-Genetic Interaction Matrix Reveals Drug Mechanism and Genetic Architecture"

**Table S20**. Pearson correlation coefficients of individual gene CRANKS scores with population doublings and essentiality RANKS scores at screen onset (pre-doxycycline to post-doxycycline)

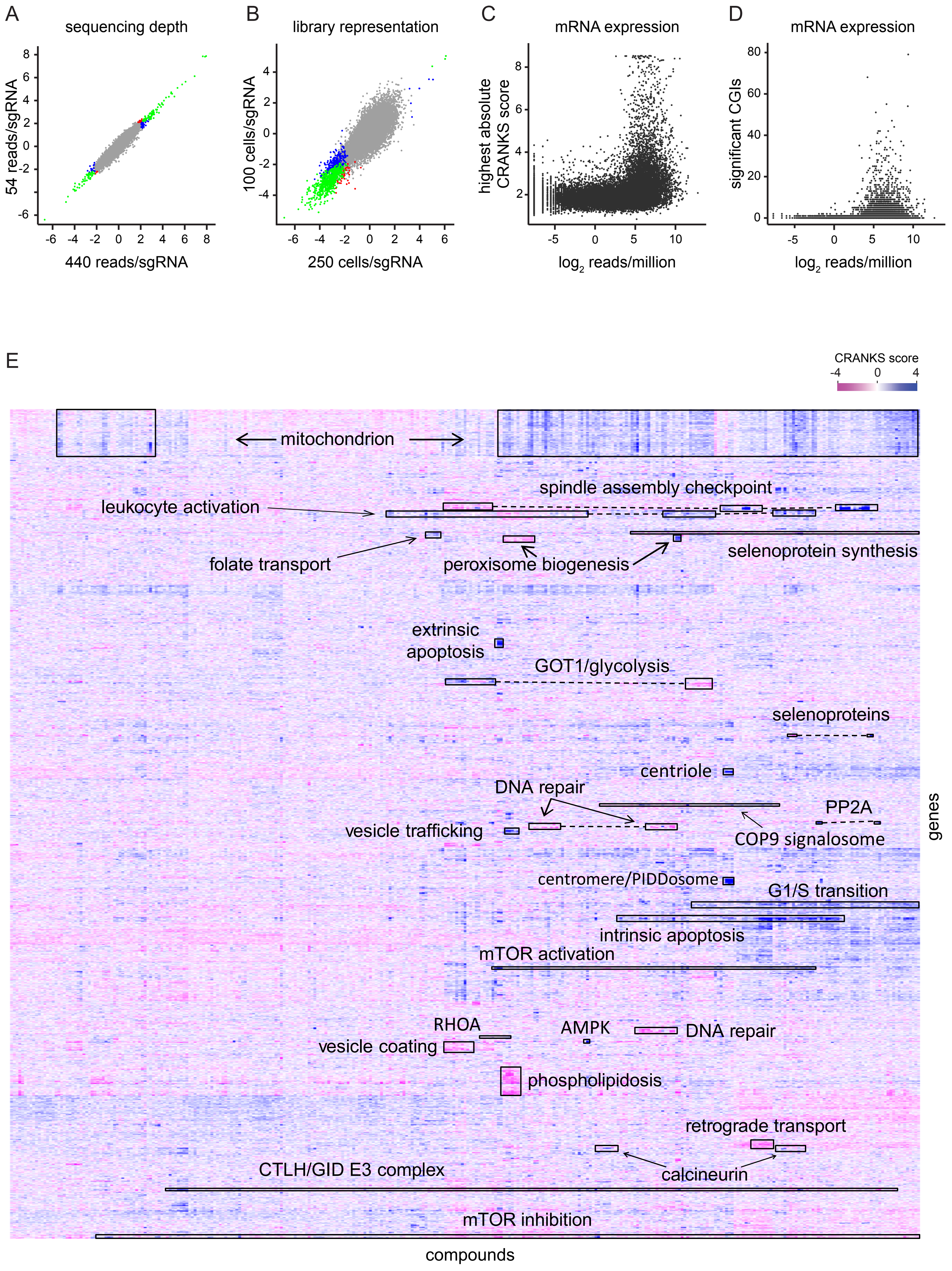
A
sequencing depth
54 reads/sgRNA
440 reads/sgRNA
B
library representation
100 cells/sgRNA
250 cells/sgRNA
C
mRNA expression
highest absolute CRANKS score
log2 reads/million
D
mRNA expression
significant CGIs
log2 reads/million
E
CRANKS score
mitochondrion
spindle assembly checkpoint
leukocyte activation
folate transport
peroxisome biogenesis
selenoprotein synthesis
extrinsic apoptosis
GOT1/glycolysis
selenoproteins
centriole
DNA repair
PP2A
vesicle trafficking
COP9 signalosome
centromere/PIDDosome
G1/S transition
intrinsic apoptosis
mTOR activation
RHOA
AMPK
DNA repair
vesicle coating
phospholipidosis
retrograde transport
CTLH/GID E3 complex
calcineurin
mTOR inhibition
compounds
genes

Figure S1

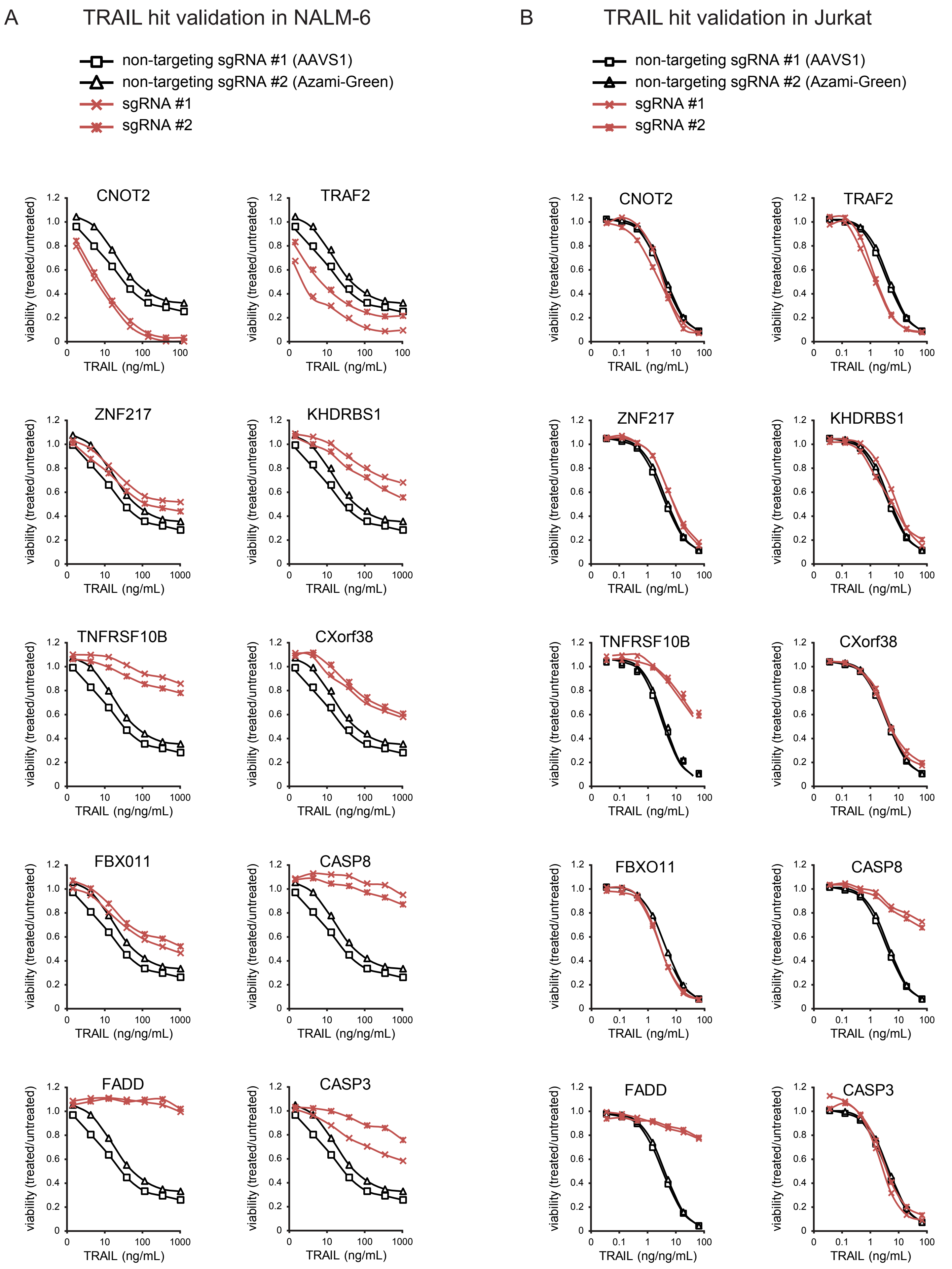
A
TRAIL hit validation in NALM-6
non-targeting sgRNA #1 (AAVS1)
non-targeting sgRNA #2 (Azami-Green)
sgRNA #1
sgRNA #2
CNOT2
TRAF2
ZNF217
KHDRBS1
TNFRSF10B
CXorf38
FBX011
CASP8
FADD
CASP3
viability (treated/untreated)
TRAIL (ng/mL)
TRAIL (ng/ng/mL)
B
TRAIL hit validation in Jurkat
non-targeting sgRNA #1 (AAVS1)
non-targeting sgRNA #2 (Azami-Green)
sgRNA #1
sgRNA #2
CNOT2
TRAF2
ZNF217
KHDRBS1
TNFRSF10B
CXorf38
FBXO11
CASP8
FADD
CASP3
viability (treated/untreated)
TRAIL (ng/mL)
TRAIL (ng/ng/mL)

Figure S2

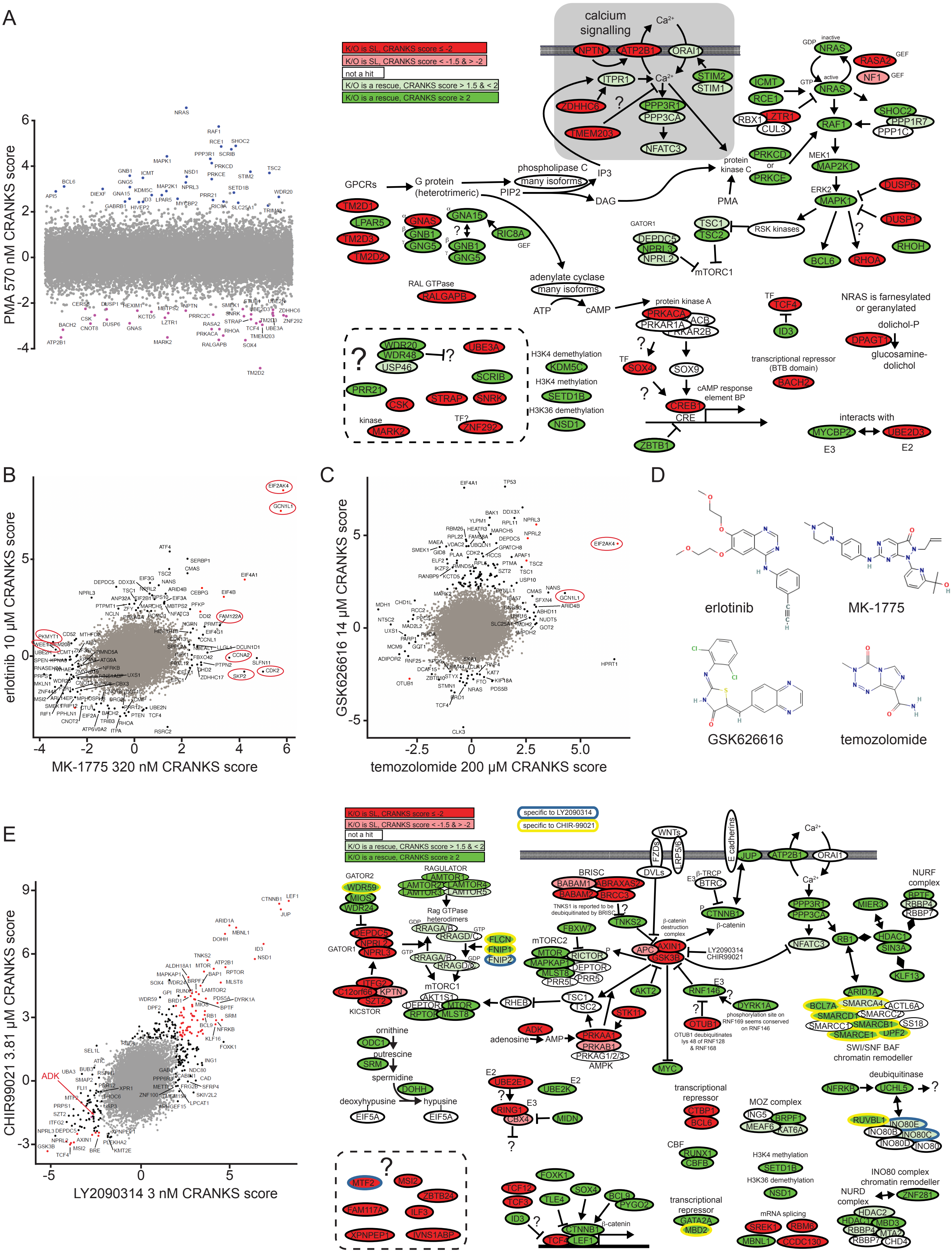
A
B
C
D
E
K/O is SL, CRANKS score ≤ -2
K/O is SL, CRANKS score < -1.5 & > -2
not a hit
K/O is a rescue, CRANKS score > 1.5 & < 2
K/O is a rescue, CRANKS score ≥ 2
PMA 570 nM CRANKS score
calcium signalling
GPCRs
G protein (heterotrimeric)
phospholipase C
many isoforms
PIP2
IP3
DAG
protein kinase C
PMA
adenylate cyclase
ATP
cAMP
protein kinase A
cAMP response element BP
CRE
RSK kinases
mTORC1
GATOR1
NRAS is farnesylated or geranylated
dolichol-P
glucosamine-dolichol
transcriptional repressor (BTB domain)
interacts with
H3K4 demethylation
H3K4 methylation
H3K36 demethylation
RAL GTPase
erlotinib 10 μM CRANKS score
MK-1775 320 nM CRANKS score
GSK626616 14 μM CRANKS score
temozolomide 200 μM CRANKS score
erlotinib
MK-1775
GSK626616
temozolomide
specific to LY2090314
specific to CHIR-99021
CHIR99021 3.81 μM CRANKS score
LY2090314 3 nM CRANKS score
GATOR2
RAGULATOR
Rag GTPase heterodimers
GATOR1
KICSTOR
mTORC1
mTORC2
BRISC
TNKS1 is reported to be deubiquitinated by BRISC
β-catenin destruction complex
β-catenin
AMPK
ornithine
putrescine
spermidine
deoxyhypusine
hypusine
adenosine
AMP
phosphorylation site on RNF169 seems conserved on RNF146
OTUB1 deubiquitinates lys 48 of RNF128 & RNF168
transcriptional repressor
MOZ complex
CBF
H3K4 methylation
H3K36 demethylation
mRNA splicing
NURF complex
SWI/SNF BAF chromatin remodeller
deubiquitinase
INO80 complex chromatin remodeller
NURD complex

Figure S3

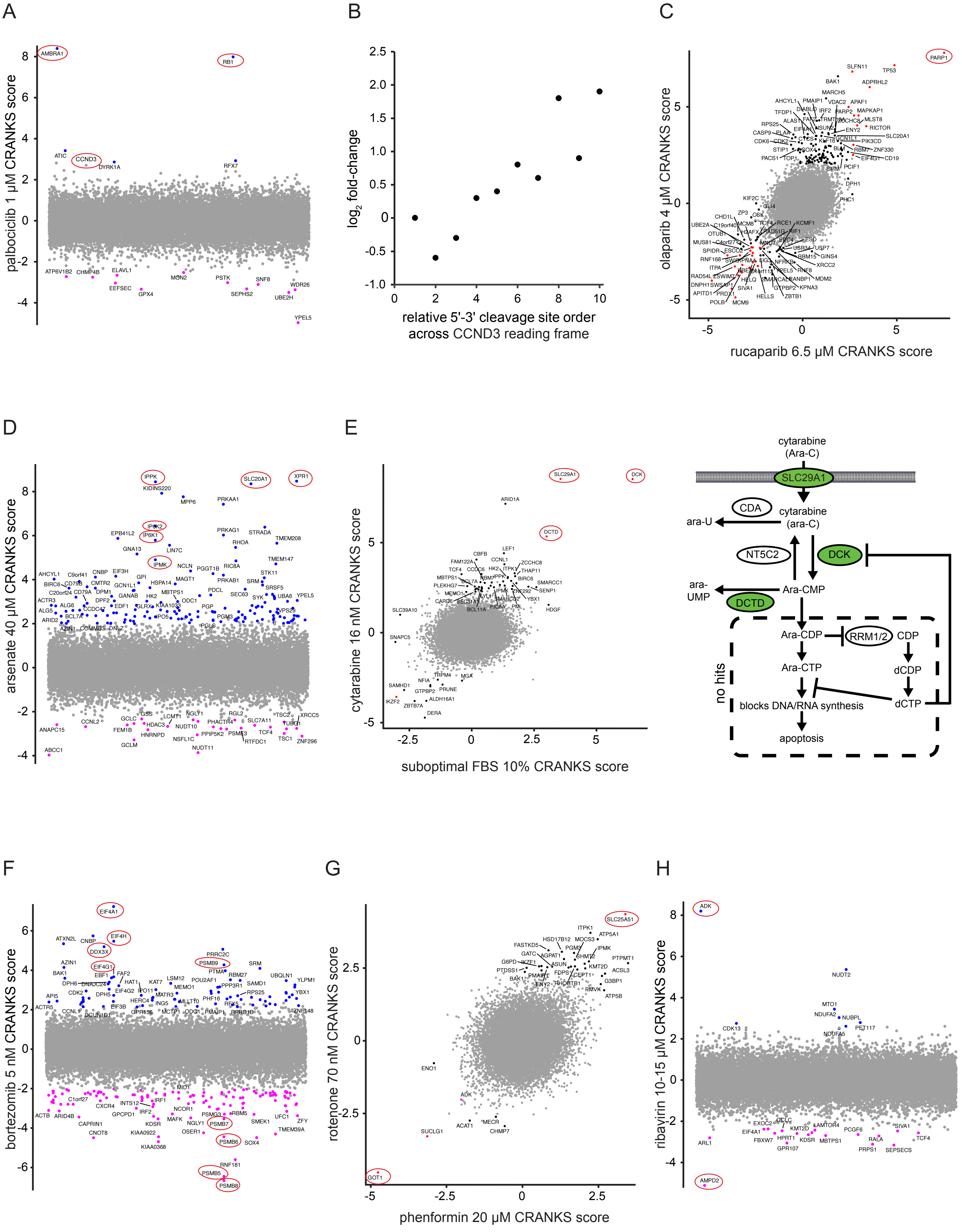
A
B
C
D
E
F
G
H
palbociclib 1 μM CRANKS score
log2 fold-change
relative 5'-3' cleavage site order across CCND3 reading frame
olaparib 4 μM CRANKS score
rucaparib 6.5 μM CRANKS score
arsenate 40 μM CRANKS score
cytarabine 16 nM CRANKS score
suboptimal FBS 10% CRANKS score
cytarabine (Ara-C)
SLC29A1
CDA
cytarabine (ara-C)
ara-U
NT5C2
DCK
ara-UMP
Ara-CMP
DCTD
Ara-CDP
RRM1/2
CDP
Ara-CTP
dCDP
no hits
blocks DNA/RNA synthesis
dCTP
apoptosis
bortezomib 5 nM CRANKS score
rotenone 70 nM CRANKS score
phenformin 20 μM CRANKS score
ribavirin 10-15 μM CRANKS score



Figure S4

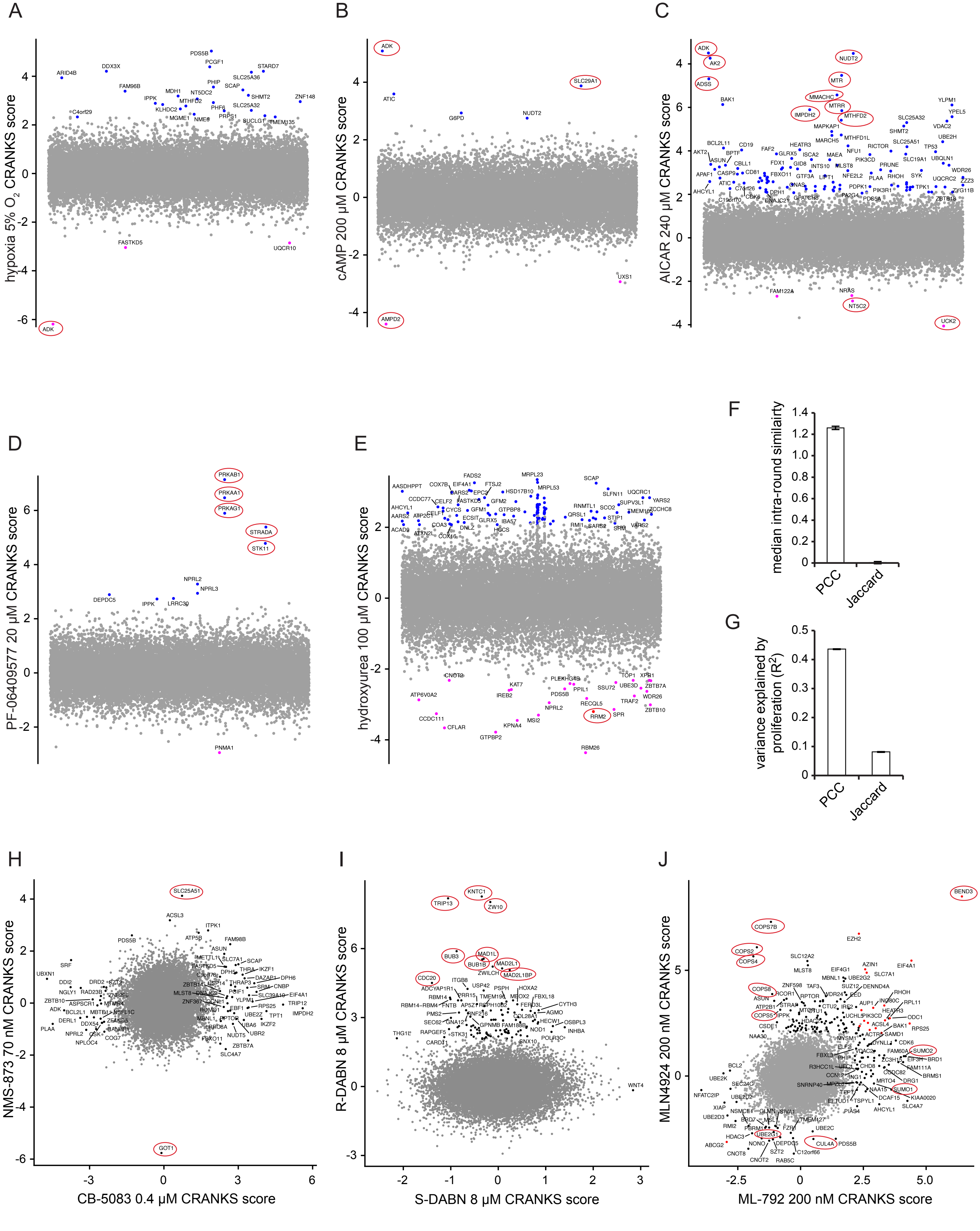
A
hypoxia 5% $O_2$ CRANKS score
ADK
FASTKD5
UQCR10
PDS5B
ARID4B
DDX3X
B
cAMP 200 µM CRANKS score
ADK
SLC29A1
AMPD2
ATIC
G6PD
NUDT2
UXS1
C
AICAR 240 µM CRANKS score
ADK
AK2
ADSS
NUDT2
MTR
MMACHC
MTRR
IMPDH2
MTHFD2
NT5C2
UCK2
BAK1
D
PF-06409577 20 µM CRANKS score
PRKAB1
PRKAA1
PRKAG1
STRADA
STK11
NPRL2
NPRL3
DEPDC5
PNMA1
E
hydroxyurea 100 µM CRANKS score
RRM2
RBM26
F
median intra-round similairty
PCC
Jaccard
G
variance explained by proliferation ($R^2$)
PCC
Jaccard
H
NMS-873 70 nM CRANKS score
CB-5083 0.4 µM CRANKS score
SLC25A51
GOT1
I
R-DABN 8 µM CRANKS score
S-DABN 8 µM CRANKS score
TRIP13
KNTC1
ZW10
BUB3
MAD1L
BUB1B
MAD2L1
MAD2L1BP
CDC20
WNT4
J
MLN4924 200 nM CRANKS score
ML-792 200 nM CRANKS score
BEND3
COPS7B
COPS2
COPS4
COPS8
COPS5
SUMO2
SUMO1
UBE2G1
CUL4A
EZH2



Figure S5

A

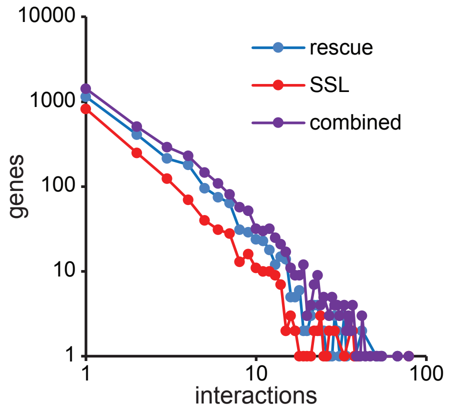


B

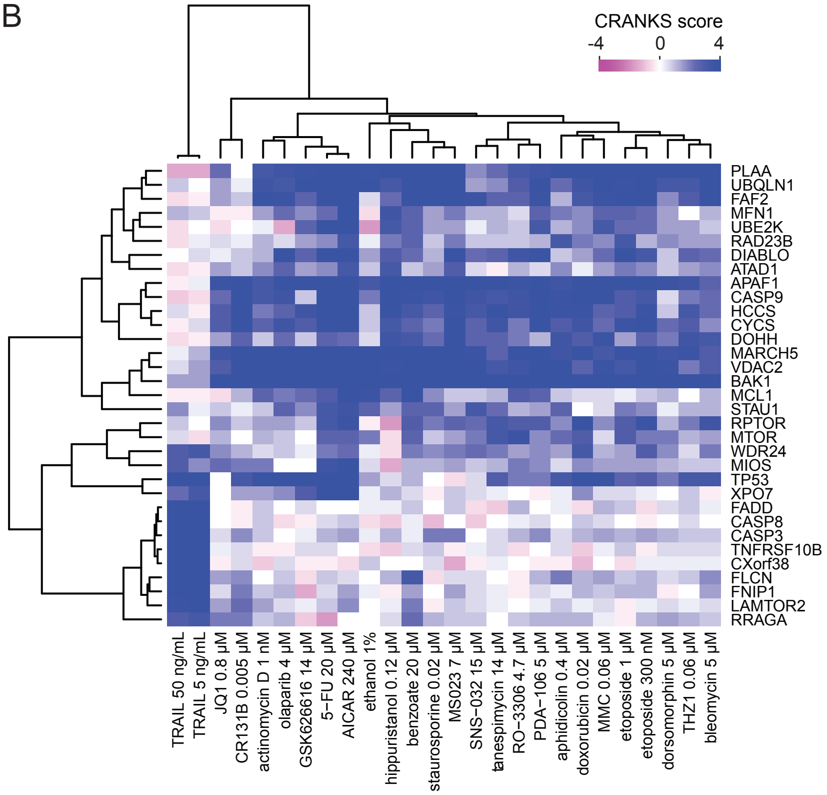


C

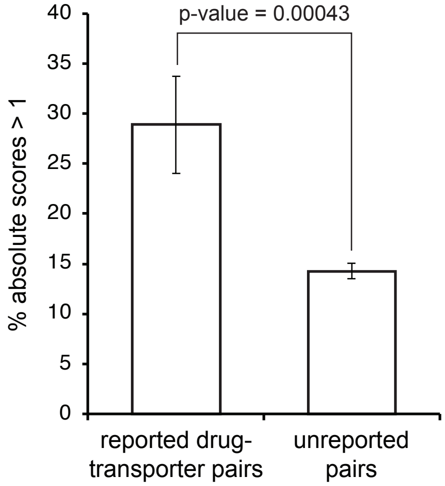


D

| compound | dose | ABCC1 CRANKS score | reported effect | PMID |
|---|---|---|---|---|
| rapamycin | 10 µM | -7.22 | substrate of P-glycoproteins | 9223585 |
| menadione | 5 µM | -5.84 | ABCG2 substrate | 18065489 |
| thimerosal | 0.85µM | -5.26 | Hg is an ABCC1 substrate | 15585369 |
| vincristine | 0.5 nM | -5.09 | ABCC1 substrate | 15717060 |
| YM155 | 1 nM | -4.55 | ABCB1 substrate | 21205082 |
| etoposide | 100 nM | -4.25 | ABCC1 substrate | 15717060 |
| arsenate | 40 µM | -3.98 | As is exported by ABCC1 | 15161912 |
| colchicine | 9 nM | -3.85 | ABCB1 substrate | 15717060 |
| UM0131023 | 50 nM | -3.84 | N/A | N/A |
| $As_2O_3$ | 60 µM | -3.29 | As is exported by ABCC1 | 15161912 |
| ibrutinib | 10 µM | -3.24 | inhibits ABCC1 | 25164592 |
| 6 others | N/A | -2 to -3 | N/A | N/A |

E

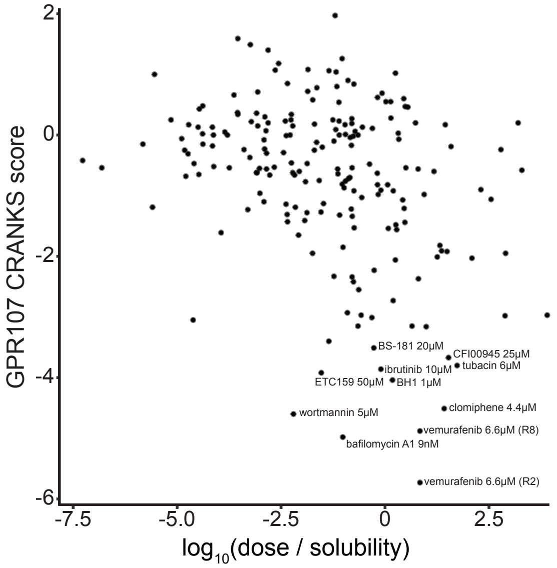


F

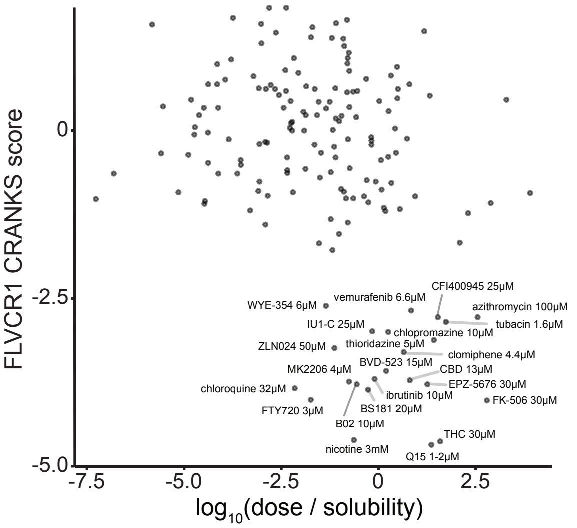


G

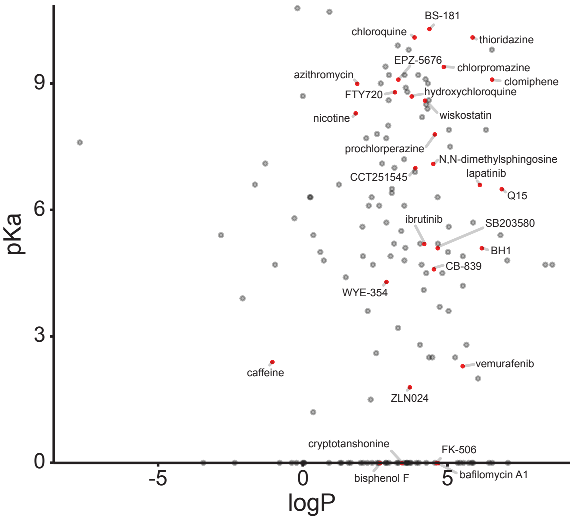




Figure S6

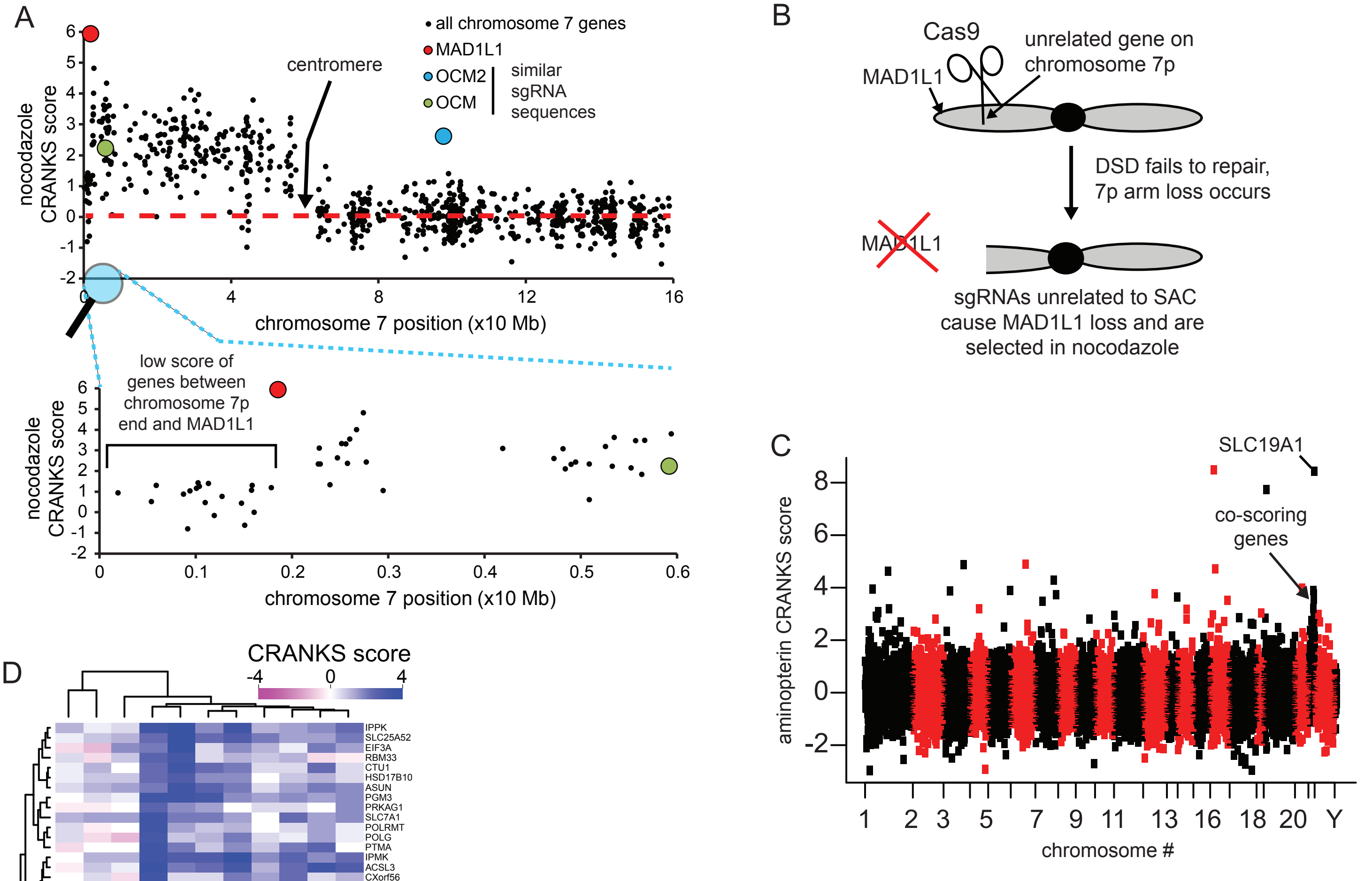


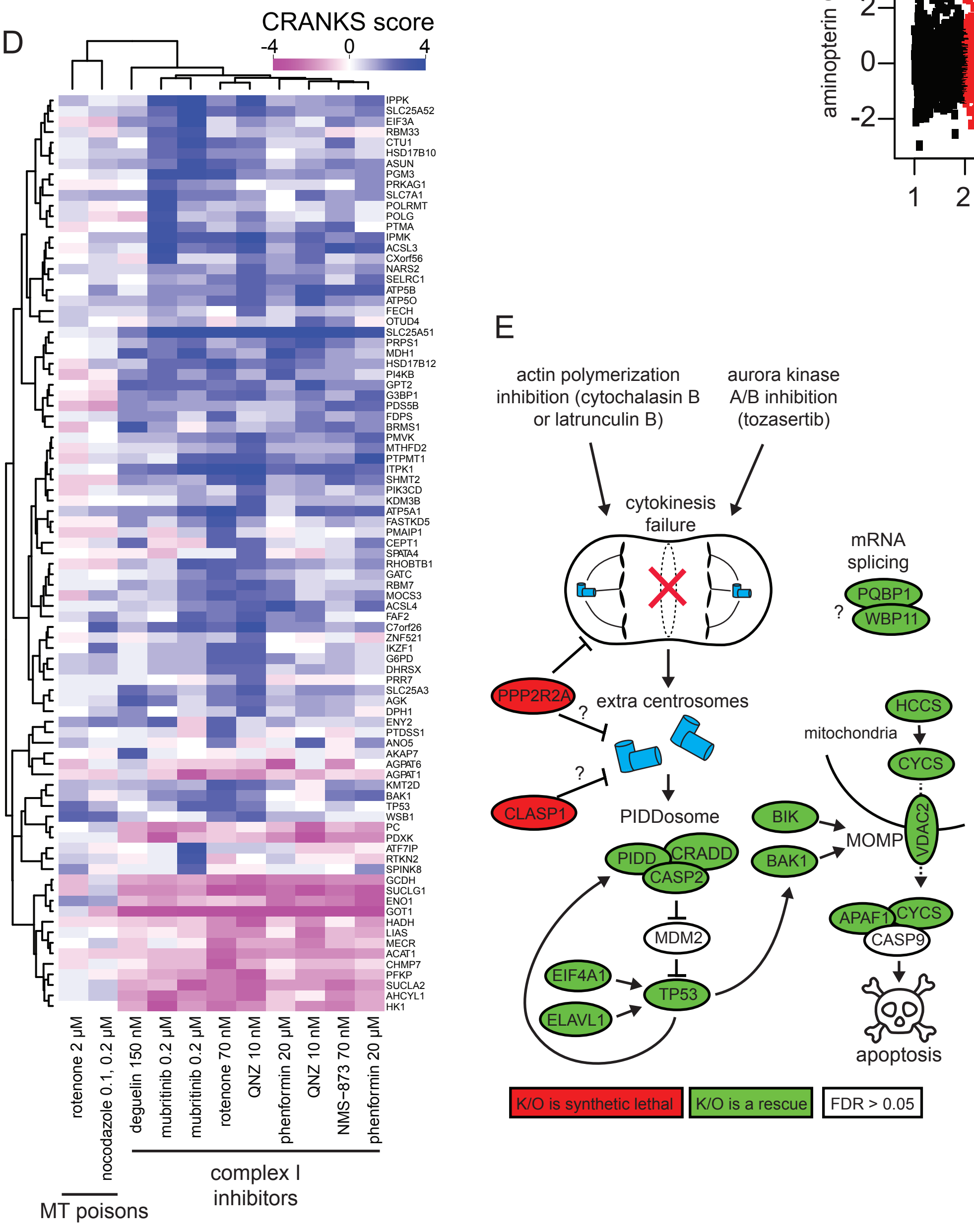


F

| gene | rank | CRANKS score | p-value | FDR | note |
|---|---|---|---|---|---|
| CLASP1 | 1 | -2.87 | 4.00E-06 | 0.024 | |
| PPP2R2A | 2 | -2.85 | 5.00E-06 | 0.024 | cytokinesis acticator |
| SASS6 | 18694 | 1.53 | 0.01237 | 0.338 | centrosome |
| POC1A | 18861 | 1.94 | 0.00104 | 0.066 | centrosome |
| CASP9 | 18870 | 1.99 | 0.00079 | 0.055 | proapoptotic |
| BIK | 18896 | 2.17 | 0.00028 | 0.023 | proapoptotic |
| KIAA1731 | 18900 | 2.2 | 0.00049 | 0.037 | centrosome |
| RBM14 | 18902 | 2.2 | 0.00102 | 0.065 | centrosome |
| CYCS | 18907 | 2.25 | 0.00018 | 0.016 | proapoptotic |
| NEURL4 | 18914 | 2.35 | 9.90E-05 | 0.009 | centrosome |
| CEP350 | 18920 | 2.4 | 0.00017 | 0.015 | centrosome |
| HCCS | 18937 | 2.54 | 3.20E-05 | 0.003 | proapoptotic |
| SDCCAG8 | 18949 | 2.78 | 7.00E-06 | 0.001 | centrosome |
| FGFR1OP | 18951 | 2.83 | 1.60E-05 | 0.002 | centrosome |
| CNTRL | 18958 | 3.02 | 5.00E-06 | 0 | centrosome |
| MARCH5 | 18959 | 3.04 | 2.00E-06 | 0 | proapoptotic |
| UBQLN1 | 18962 | 3.08 | 1.00E-06 | 0 | proapoptotic |
| CENPJ | 18966 | 3.18 | 2.00E-06 | 0 | centrosome |
| VDAC2 | 18968 | 3.26 | 1.00E-06 | 0 | proapoptotic |
| APAF1 | 18969 | 3.28 | 1.00E-06 | 0 | proapoptotic |
| CEP152 | 18980 | 3.66 | 1.00E-06 | 0 | centrosome |
| PQBP1 | 18984 | 3.81 | 1.00E-06 | 0 | mRNA splicing |
| KIAA0586 | 18986 | 4.23 | 1.00E-06 | 0 | centrosome |
| ELAVL1 | 18987 | 4.36 | 1.00E-06 | 0 | p53-related |
| BAK1 | 18988 | 4.39 | 1.00E-06 | 0 | proapoptotic |
| WBP11 | 18989 | 4.41 | 1.00E-06 | 0 | mRNA splicing |
| FOPNL | 18990 | 4.62 | 1.00E-06 | 0 | centrosome |
| HYLS1 | 18991 | 4.85 | 1.00E-06 | 0 | centrosome |
| TUBD1 | 18992 | 4.97 | 1.00E-06 | 0 | centrosome |
| RTTN | 18993 | 5.01 | 1.00E-06 | 0 | centrosome |
| BUB3 | 18994 | 5.03 | 1.00E-06 | 0 | |
| CEP78 | 18995 | 5.03 | 1.00E-06 | 0 | centrosome |
| TUBE1 | 18996 | 5.15 | 1.00E-06 | 0 | centrosome |
| TP53 | 18997 | 5.24 | 1.00E-06 | 0 | p53-related |
| SCLT1 | 18998 | 5.34 | 1.00E-06 | 0 | centrosome |
| EIF4A1 | 18999 | 5.4 | 1.00E-06 | 0 | p53-related |
| CEP44 | 19000 | 5.49 | 1.00E-06 | 0 | centrosome |
| C16orf59/TEDC2 | 19001 | 5.99 | 1.00E-06 | 0 | centrosome |
| NIN | 19002 | 6.39 | 1.00E-06 | 0 | centrosome |
| ODF2 | 19003 | 6.53 | 1.00E-06 | 0 | centrosome |
| CASP2 | 19004 | 6.56 | 1.00E-06 | 0 | PIDDosome |
| ANKRD26 | 19005 | 6.75 | 1.00E-06 | 0 | centrosome |
| CEP76 | 19006 | 6.76 | 1.00E-06 | 0 | centrosome |
| CRADD | 19007 | 6.94 | 1.00E-06 | 0 | PIDDosome |
| C14orf80/TEDC1 | 19008 | 6.96 | 1.00E-06 | 0 | centrosome |
| CEP128 | 19009 | 7.06 | 1.00E-06 | 0 | centrosome |
| CEP120 | 19010 | 7.14 | 1.00E-06 | 0 | centrosome |
| PIDD | 19011 | 7.14 | 1.00E-06 | 0 | PIDDosome |
| OFD1 | 19012 | 7.36 | 1.00E-06 | 0 | centrosome |
| PIBF1 | 19013 | 7.7 | 1.00E-06 | 0 | centrosome |
| KIAA0753 | 19014 | 8.08 | 1.00E-06 | 0 | centrosome |
| CCDC41 | 19015 | 8.17 | 1.00E-06 | 0 | centrosome |
| C2CD3 | 19016 | 8.37 | 1.00E-06 | 0 | centrosome |

Figure S7

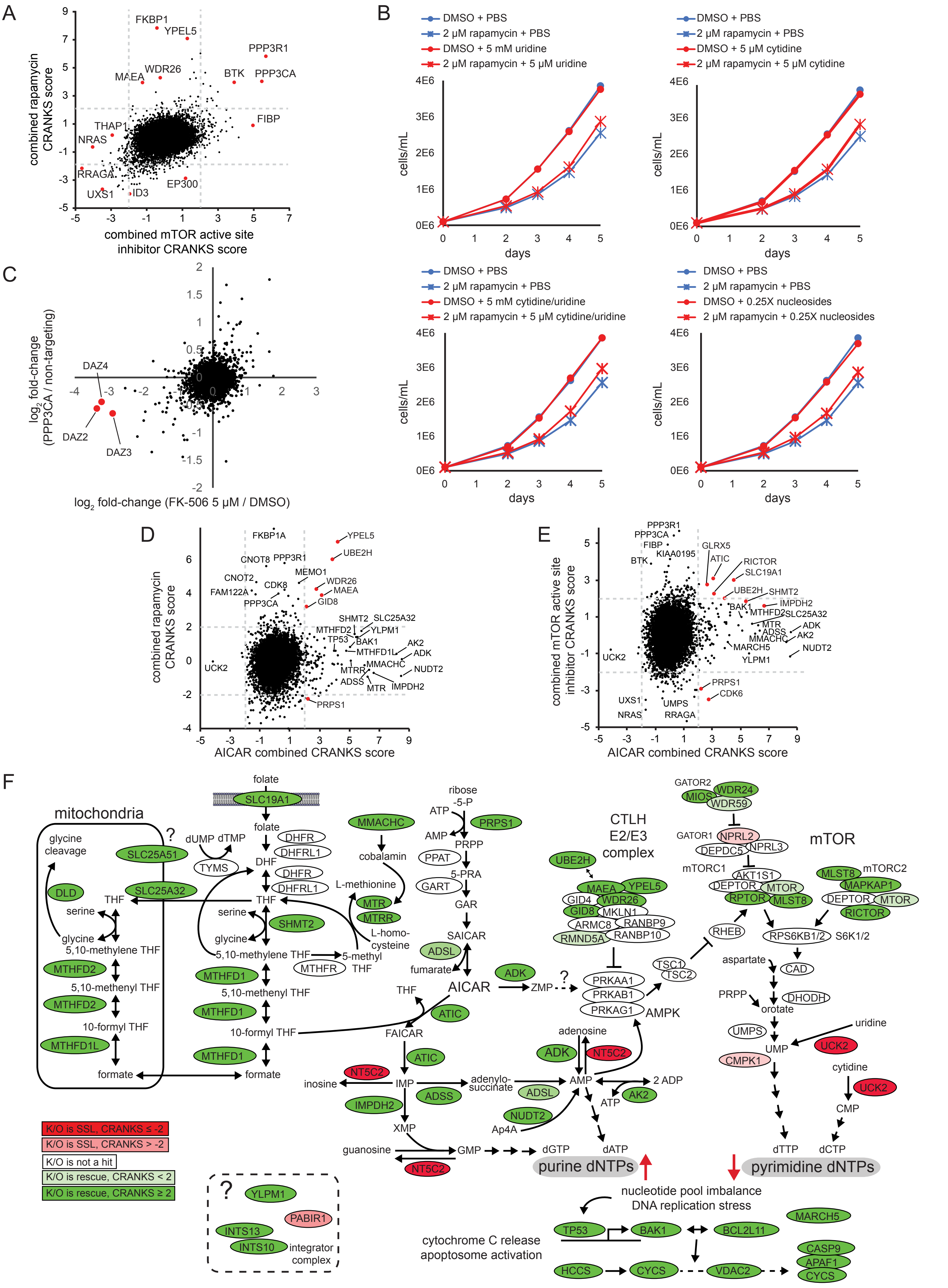
A
combined rapamycin CRANKS score
combined mTOR active site inhibitor CRANKS score
FKBP1
YPEL5
PPP3R1
WDR26
MAEA
BTK
PPP3CA
FIBP
THAP1
NRAS
RRAGA
EP300
UXS1
ID3
B
DMSO + PBS
2 μM rapamycin + PBS
DMSO + 5 mM uridine
2 μM rapamycin + 5 μM uridine
DMSO + 5 μM cytidine
2 μM rapamycin + 5 μM cytidine
DMSO + 5 mM cytidine/uridine
2 μM rapamycin + 5 μM cytidine/uridine
DMSO + 0.25X nucleosides
2 μM rapamycin + 0.25X nucleosides
cells/mL
days
C
log2 fold-change (PPP3CA / non-targeting)
log2 fold-change (FK-506 5 μM / DMSO)
DAZ4
DAZ2
DAZ3
D
combined rapamycin CRANKS score
AICAR combined CRANKS score
FKBP1A
YPEL5
UBE2H
CNOT8
PPP3R1
MEMO1
CNOT2
CDK8
FAM122A
WDR26
MAEA
GID8
PPP3CA
SHMT2
SLC25A32
MTHFD2
YLPM1
TP53
BAK1
AK2
MTHFD1L
ADK
MMACHC
NUDT2
MTRR
ADSS
MTR
IMPDH2
UCK2
PRPS1
E
combined mTOR active site inhibitor CRANKS score
AICAR combined CRANKS score
PPP3R1
PPP3CA
FIBP
KIAA0195
BTK
GLRX5
ATIC
RICTOR
SLC19A1
UBE2H
SHMT2
BAK1
IMPDH2
MTHFD2
SLC25A32
MTR
ADK
ADSS
MMACHC
AK2
MARCH5
NUDT2
YLPM1
UCK2
PRPS1
CDK6
UXS1
UMPS
NRAS
RRAGA
F
mitochondria
glycine cleavage
DLD
SLC25A51
SLC25A32
THF
serine
glycine
5,10-methylene THF
MTHFD2
5,10-methenyl THF
MTHFD2
10-formyl THF
MTHFD1L
formate
folate
SLC19A1
folate
dUMP dTMP
TYMS
DHF
DHFR
DHFRL1
DHFR
DHFRL1
THF
serine
SHMT2
glycine
5,10-methylene THF
MTHFR
5-methyl THF
MTHFD1
5,10-methenyl THF
MTHFD1
10-formyl THF
MTHFD1
formate
MMACHC
cobalamin
L-methionine
MTR
MTRR
L-homo-cysteine
ribose-5-P
ATP
AMP
PRPS1
PRPP
PPAT
5-PRA
GART
GAR
SAICAR
ADSL
fumarate
AICAR
ADK
ZMP
THF
ATIC
FAICAR
ATIC
NT5C2
inosine
IMP
adenylo-succinate
ADSS
ADSL
IMPDH2
XMP
guanosine
GMP
NT5C2
dGTP
dATP
purine dNTPs
CTLH E2/E3 complex
UBE2H
MAEA
YPEL5
GID4
WDR26
GID8
MKLN1
ARMC8
RANBP9
RMND5A
RANBP10
PRKAA1
PRKAB1
PRKAG1
AMPK
adenosine
ADK
NT5C2
AMP
2 ADP
ATP
AK2
NUDT2
Ap4A
GATOR2
MIOS
WDR24
WDR59
GATOR1
NPRL2
DEPDC5
NPRL3
mTOR
mTORC1
AKT1S1
DEPTOR
MTOR
RPTOR
MLST8
mTORC2
MLST8
MAPKAP1
DEPTOR
MTOR
RICTOR
RHEB
TSC1
TSC2
RPS6KB1/2
S6K1/2
aspartate
CAD
PRPP
DHODH
orotate
UMPS
uridine
UCK2
UMP
CMPK1
cytidine
UCK2
CMP
dTTP
dCTP
pyrimidine dNTPs
nucleotide pool imbalance
DNA replication stress
TP53
BAK1
BCL2L11
MARCH5
cytochrome C release
apoptosome activation
CASP9
APAF1
CYCS
HCCS
CYCS
VDAC2
K/O is SSL, CRANKS ≤ -2
K/O is SSL, CRANKS > -2
K/O is not a hit
K/O is rescue, CRANKS < 2
K/O is rescue, CRANKS ≥ 2
?
YLPM1
PABIR1
INTS13
INTS10
integrator complex



Figure S8

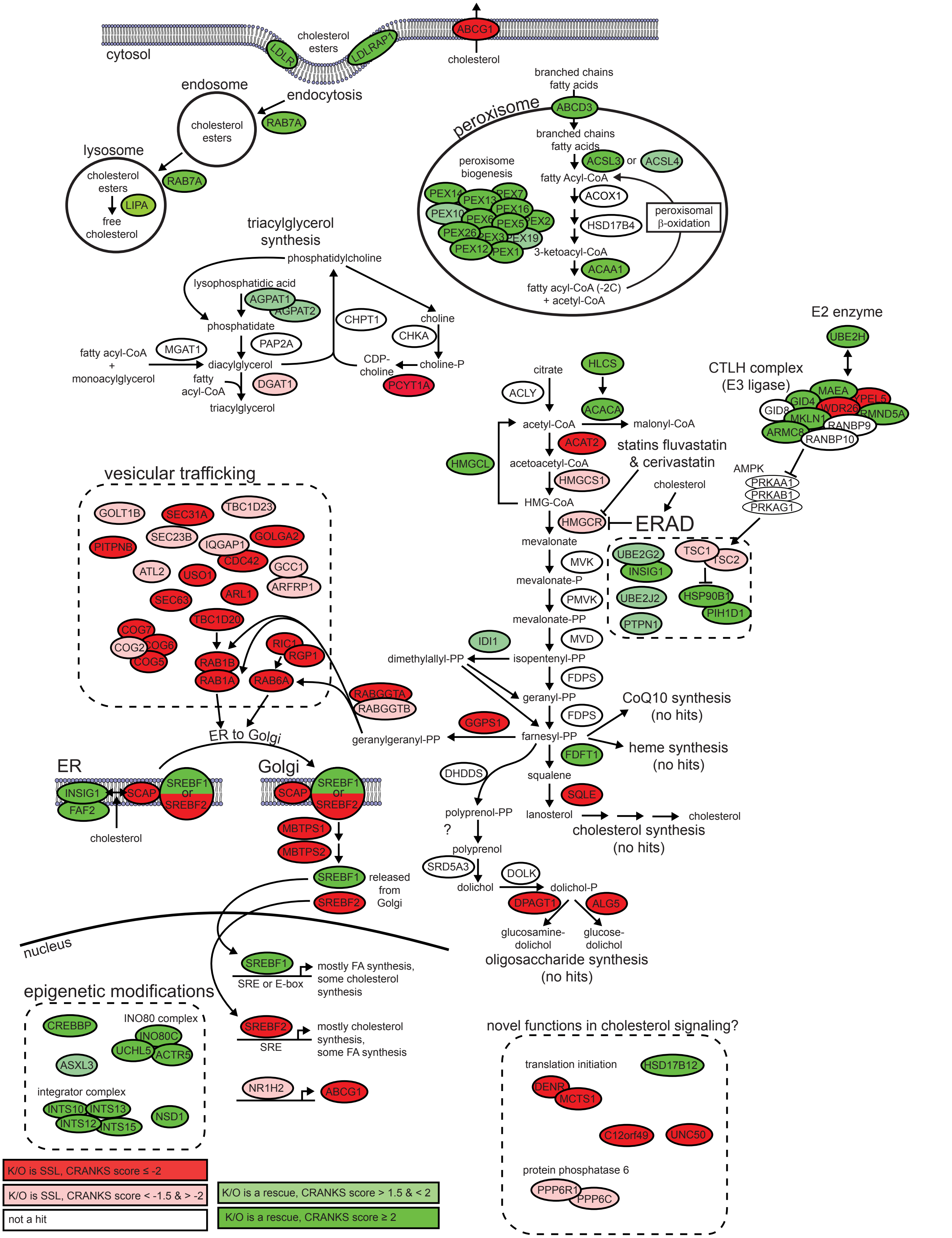

cytosol
cholesterol esters
LDLR
LDLRAP1
ABCG1
cholesterol
endosome
endocytosis
cholesterol esters
RAB7A
lysosome
cholesterol esters
LIPA
free cholesterol
RAB7A
peroxisome
branched chains fatty acids
ABCD3
branched chains fatty acids
ACSL3
or
ACSL4
fatty Acyl-CoA
ACOX1
HSD17B4
3-ketoacyl-CoA
ACAA1
fatty acyl-CoA (-2C) + acetyl-CoA
peroxisomal β-oxidation
peroxisome biogenesis
PEX14
PEX7
PEX13
PEX10
PEX16
PEX6
PEX5
PEX2
PEX26
PEX3
PEX19
PEX12
PEX1
triacylglycerol synthesis
phosphatidylcholine
lysophosphatidic acid
AGPAT1
AGPAT2
phosphatidate
CHPT1
choline
CHKA
PAP2A
MGAT1
fatty acyl-CoA + monoacylglycerol
diacylglycerol
fatty acyl-CoA
DGAT1
triacylglycerol
CDP-choline
choline-P
PCYT1A
E2 enzyme
UBE2H
CTLH complex (E3 ligase)
MAEA
GID4
YPEL5
GID8
WDR26
RMND5A
MKLN1
RANBP9
ARMC8
RANBP10
citrate
HLCS
ACLY
ACACA
acetyl-CoA
malonyl-CoA
ACAT2
HMGCL
acetoacetyl-CoA
HMGCS1
HMG-CoA
statins fluvastatin & cerivastatin
cholesterol
AMPK
PRKAA1
PRKAB1
PRKAG1
HMGCR
ERAD
mevalonate
MVK
mevalonate-P
PMVK
mevalonate-PP
MVD
IDI1
dimethylallyl-PP
isopentenyl-PP
FDPS
geranyl-PP
FDPS
GGPS1
farnesyl-PP
geranylgeranyl-PP
CoQ10 synthesis (no hits)
heme synthesis (no hits)
FDFT1
squalene
SQLE
lanosterol
cholesterol
cholesterol synthesis (no hits)
DHDDS
polyprenol-PP
?
polyprenol
SRD5A3
dolichol
DOLK
dolichol-P
DPAGT1
ALG5
glucosamine-dolichol
glucose-dolichol
oligosaccharide synthesis (no hits)
UBE2G2
INSIG1
TSC1
TSC2
UBE2J2
HSP90B1
PIH1D1
PTPN1
vesicular trafficking
GOLT1B
SEC31A
TBC1D23
SEC23B
GOLGA2
PITPNB
IQGAP1
CDC42
GCC1
ATL2
USO1
ARFRP1
ARL1
SEC63
TBC1D20
COG7
COG2
COG6
COG5
RAB1B
RAB1A
RIC1
RGP1
RAB6A
RABGGTA
RABGGTB
ER to Golgi
ER
INSIG1
FAF2
SCAP
SREBF1 or SREBF2
cholesterol
Golgi
SCAP
SREBF1 or SREBF2
MBTPS1
MBTPS2
SREBF1
SREBF2
released from Golgi
nucleus
SREBF1
SRE or E-box
mostly FA synthesis, some cholesterol synthesis
SREBF2
SRE
mostly cholesterol synthesis, some FA synthesis
NR1H2
ABCG1
epigenetic modifications
CREBBP
INO80 complex
INO80C
UCHL5
ACTR5
ASXL3
integrator complex
INTS10
INTS13
INTS12
INTS15
NSD1
novel functions in cholesterol signaling?
translation initiation
HSD17B12
DENR
MCTS1
C12orf49
UNC50
protein phosphatase 6
PPP6R1
PPP6C
K/O is SSL, CRANKS score ≤ -2
K/O is SSL, CRANKS score < -1.5 & > -2
not a hit
K/O is a rescue, CRANKS score > 1.5 & < 2
K/O is a rescue, CRANKS score ≥ 2




Figure S9

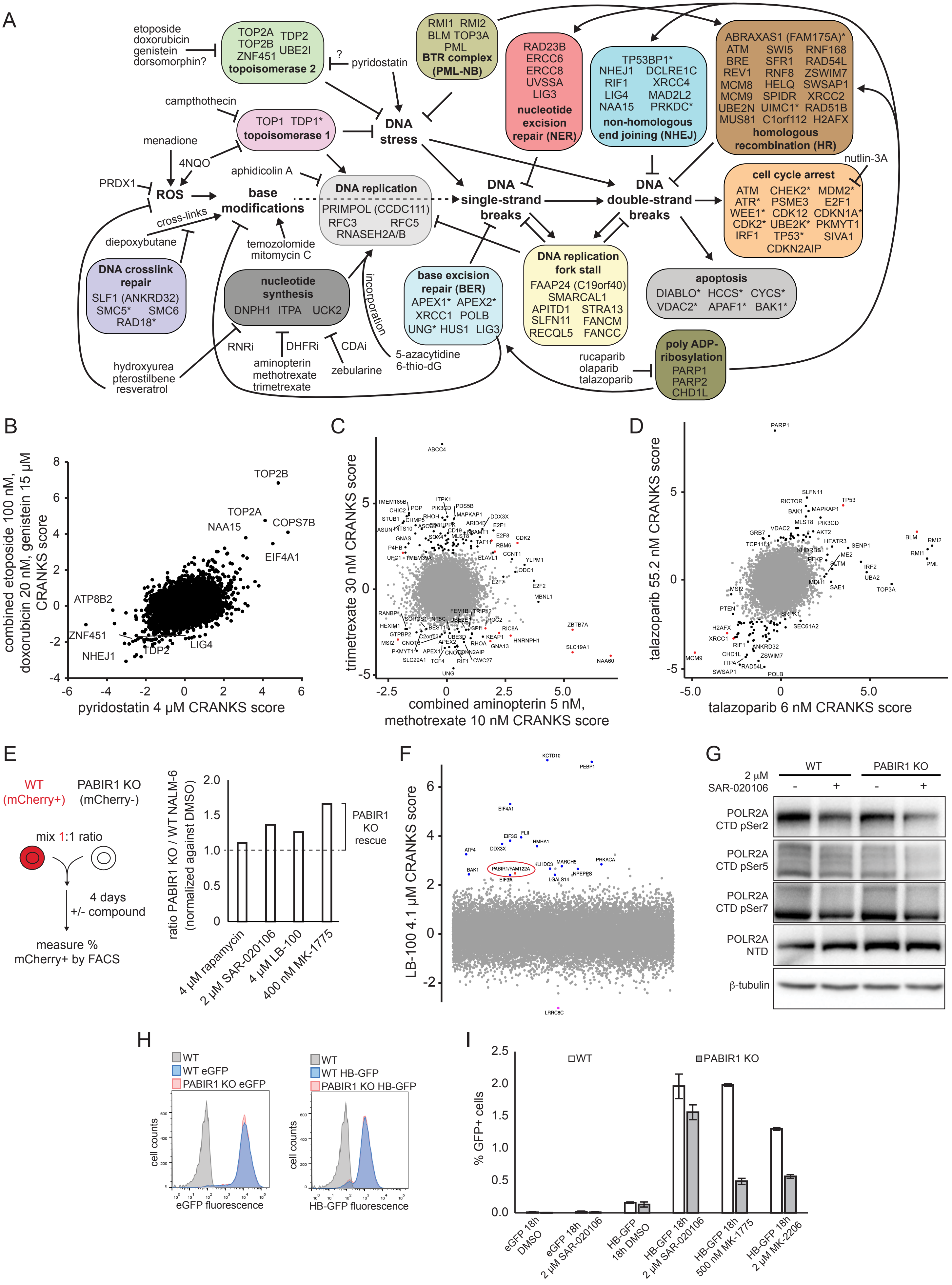
A
etoposide
doxorubicin
genistein
dorsomorphin?
TOP2A TOP2B ZNF451 TDP2 UBE2I
topoisomerase 2
?
pyridostatin
RMI1 RMI2
BLM TOP3A
PML
BTR complex
(PML-NB)
RAD23B
ERCC6
ERCC8
UVSSA
LIG3
nucleotide excision repair (NER)
TP53BP1*
NHEJ1 DCLRE1C
RIF1 XRCC4
LIG4 MAD2L2
NAA15 PRKDC*
non-homologous end joining (NHEJ)
ABRAXAS1 (FAM175A)*
ATM SWI5 RNF168
BRE SFR1 RAD54L
REV1 RNF8 ZSWIM7
MCM8 HELQ SWSAP1
MCM9 SPIDR XRCC2
UBE2N UIMC1* RAD51B
MUS81 C1orf112 H2AFX
homologous recombination (HR)
campthothecin
TOP1 TDP1*
topoisomerase 1
DNA stress
menadione
4NQO
PRDX1
ROS
base modifications
aphidicolin A
DNA replication
PRIMPOL (CCDC111)
RFC3 RFC5
RNASEH2A/B
DNA single-strand breaks
DNA double-strand breaks
nutlin-3A
cell cycle arrest
ATM CHEK2* MDM2*
ATR* PSME3 E2F1
WEE1* CDK12 CDKN1A*
CDK2* UBE2K* PKMYT1
IRF1 TP53* SIVA1
CDKN2AIP
cross-links
diepoxybutane
temozolomide
mitomycin C
DNA crosslink repair
SLF1 (ANKRD32)
SMC5* SMC6
RAD18*
nucleotide synthesis
DNPH1 ITPA UCK2
incorporation
base excision repair (BER)
APEX1* APEX2*
XRCC1 POLB
UNG* HUS1 LIG3
DNA replication fork stall
FAAP24 (C19orf40)
SMARCAL1
APITD1 STRA13
SLFN11 FANCM
RECQL5 FANCC
apoptosis
DIABLO* HCCS* CYCS*
VDAC2* APAF1* BAK1*
RNRi
DHFRi
CDAi
hydroxyurea
pterostilbene
resveratrol
aminopterin
methotrexate
trimetrexate
zebularine
5-azacytidine
6-thio-dG
rucaparib
olaparib
talazoparib
poly ADP-ribosylation
PARP1
PARP2
CHD1L
B
combined etoposide 100 nM, doxorubicin 20 nM, genistein 15 µM CRANKS score
pyridostatin 4 µM CRANKS score
TOP2B
TOP2A
COPS7B
NAA15
EIF4A1
ATP8B2
ZNF451
NHEJ1
TDP2
LIG4
C
trimetrexate 30 nM CRANKS score
combined aminopterin 5 nM, methotrexate 10 nM CRANKS score
D
talazoparib 55.2 nM CRANKS score
talazoparib 6 nM CRANKS score
PARP1
BLM
RMI2
RMI1
PML
TOP3A
MCM9
E
WT (mCherry+)
PABIR1 KO (mCherry-)
mix 1:1 ratio
4 days
+/- compound
measure % mCherry+ by FACS
ratio PABIR1 KO / WT NALM-6 (normalized against DMSO)
PABIR1 KO rescue
4 µM rapamycin
2 µM SAR-020106
4 µM LB-100
400 nM MK-1775
F
LB-100 4.1 µM CRANKS score
KCTD10
PEBP1
EIF4A1
PABIR1/FAM122A
G
WT
PABIR1 KO
2 µM SAR-020106
POLR2A CTD pSer2
POLR2A CTD pSer5
POLR2A CTD pSer7
POLR2A NTD
β-tubulin
H
WT
WT eGFP
PABIR1 KO eGFP
cell counts
eGFP fluorescence
WT
WT HB-GFP
PABIR1 KO HB-GFP
HB-GFP fluorescence
I
WT
PABIR1 KO
% GFP+ cells
eGFP 18h DMSO
eGFP 18h 2 µM SAR-020106
HB-GFP 18h DMSO
HB-GFP 18h 2 µM SAR-020106
HB-GFP 18h 500 nM MK-1775
HB-GFP 18h 2 µM MK-2206

Figure S10

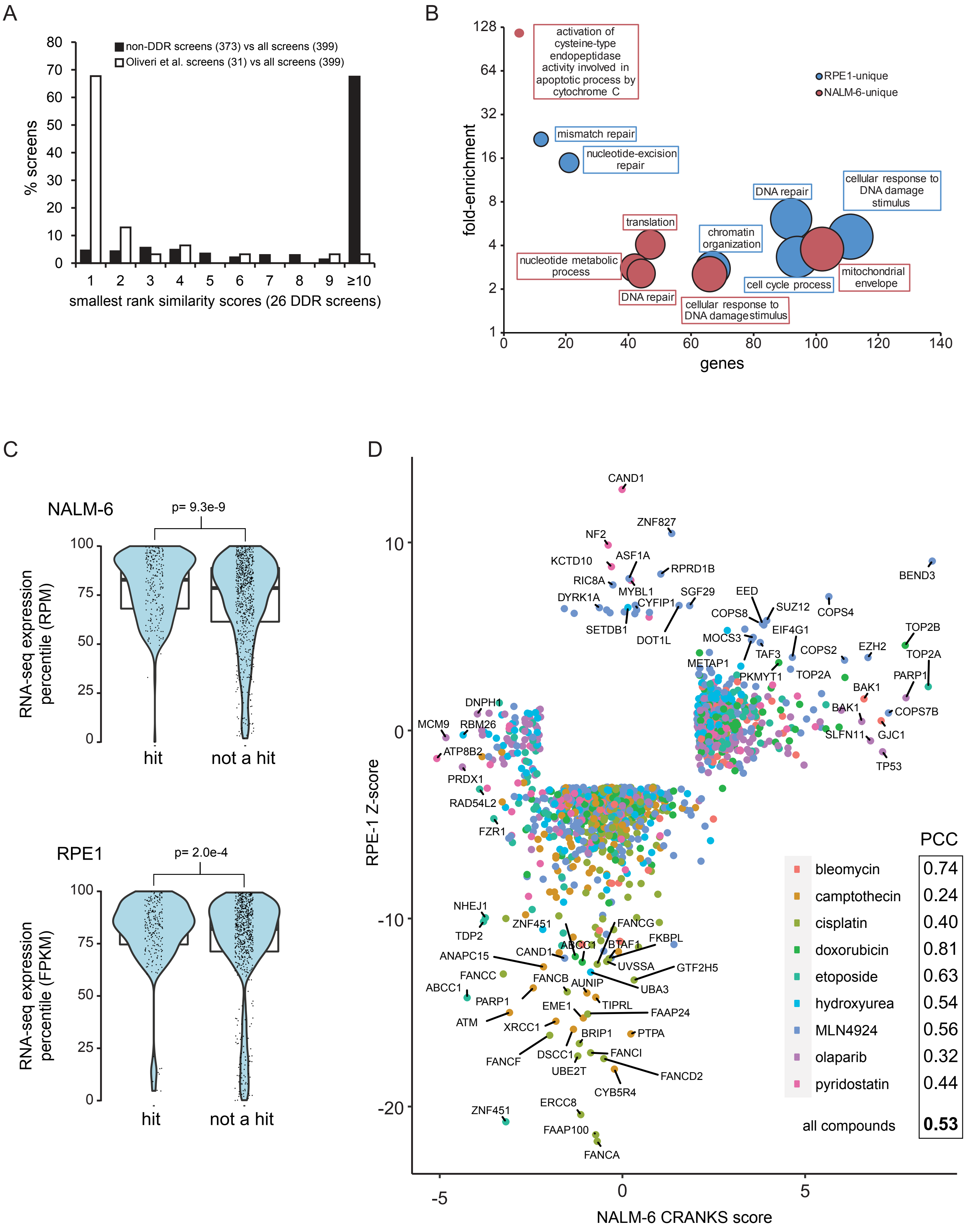
A
non-DDR screens (373) vs all screens (399)
Oliveri et al. screens (31) vs all screens (399)
% screens
smallest rank similarity scores (26 DDR screens)
B
fold-enrichment
genes
RPE1-unique
NALM-6-unique
activation of cysteine-type endopeptidase activity involved in apoptotic process by cytochrome C
mismatch repair
nucleotide-excision repair
DNA repair
cellular response to DNA damage stimulus
translation
chromatin organization
nucleotide metabolic process
DNA repair
cell cycle process
mitochondrial envelope
cellular response to DNA damagestimulus
C
NALM-6
p= 9.3e-9
RNA-seq expression percentile (RPM)
hit
not a hit
RPE1
p= 2.0e-4
RNA-seq expression percentile (FPKM)
hit
not a hit
D
RPE-1 Z-score
NALM-6 CRANKS score
PCC
bleomycin 0.74
camptothecin 0.24
cisplatin 0.40
doxorubicin 0.81
etoposide 0.63
hydroxyurea 0.54
MLN4924 0.56
olaparib 0.32
pyridostatin 0.44
all compounds 0.53



Figure S11

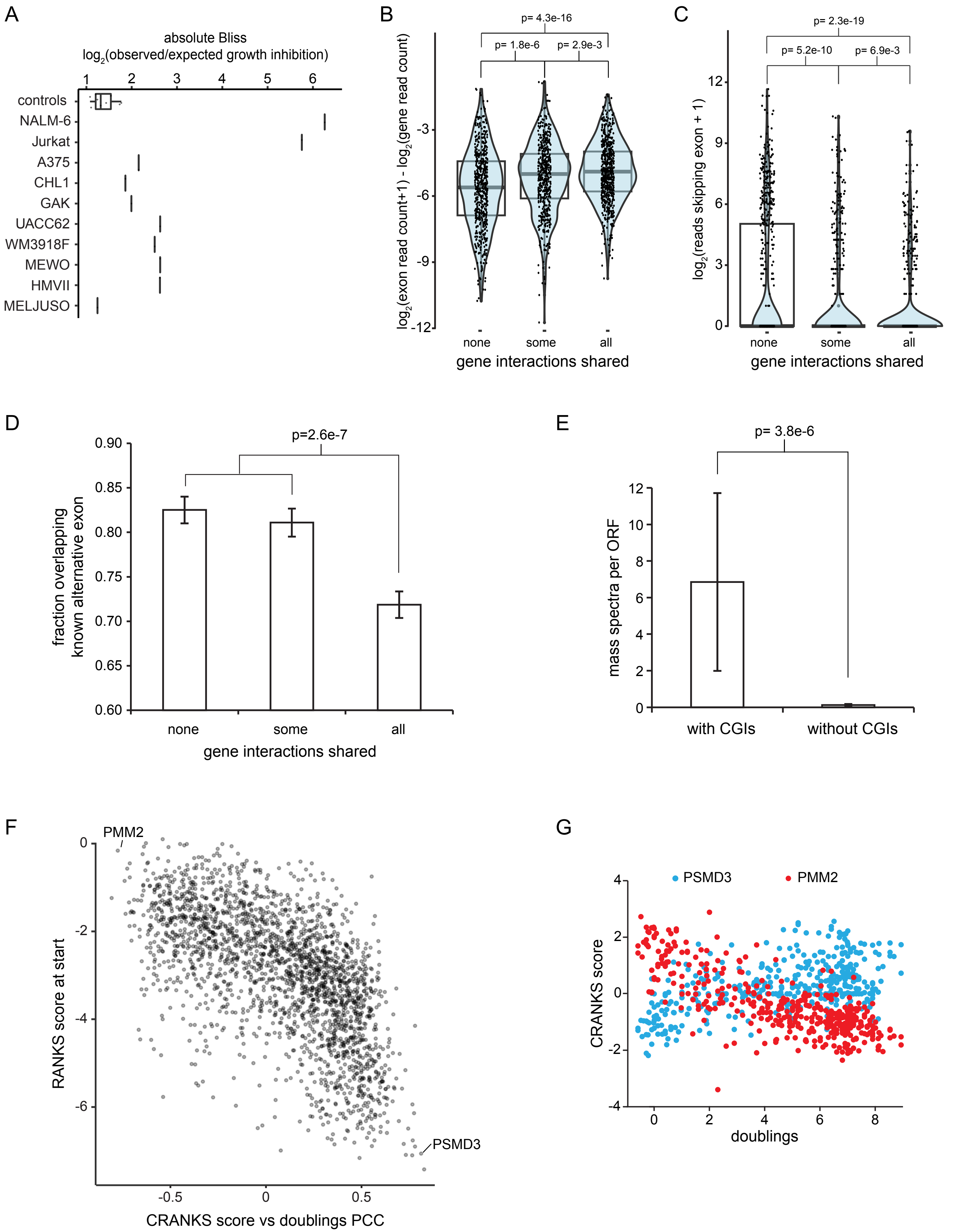
A
absolute Bliss
log2(observed/expected growth inhibition)
1 2 3 4 5 6
controls
NALM-6
Jurkat
A375
CHL1
GAK
UACC62
WM3918F
MEWO
HMVII
MELJUSO
B
p= 4.3e-16
p= 1.8e-6
p= 2.9e-3
log2(exon read count+1) - log2(gene read count)
-3
-6
-9
-12
none
some
all
gene interactions shared
C
p= 2.3e-19
p= 5.2e-10
p= 6.9e-3
log2(reads skipping exon + 1)
12
9
6
3
0
none
some
all
gene interactions shared
D
p=2.6e-7
fraction overlapping known alternative exon
0.90
0.85
0.80
0.75
0.70
0.65
0.60
none
some
all
gene interactions shared
E
p= 3.8e-6
mass spectra per ORF
12
10
8
6
4
2
0
with CGIs
without CGIs
F
PMM2
PSMD3
RANKS score at start
0
-2
-4
-6
-0.5
0
0.5
CRANKS score vs doublings PCC
G
PSMD3
PMM2
CRANKS score
4
2
0
-2
-4
0
2
4
6
8
doublings



Figure S12

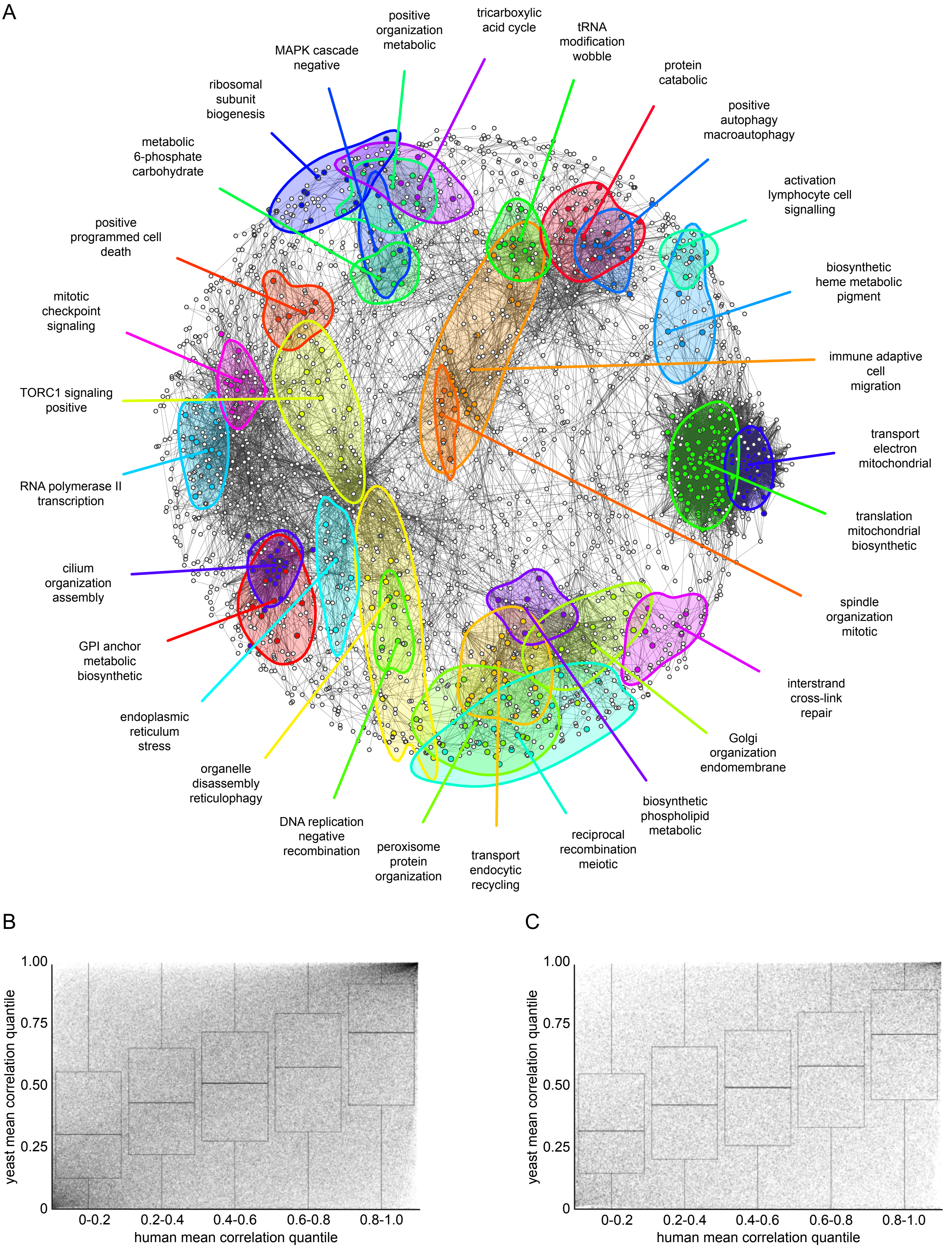
A
positive organization metabolic
tricarboxylic acid cycle
tRNA modification wobble
MAPK cascade negative
protein catabolic
ribosomal subunit biogenesis
positive autophagy macroautophagy
metabolic 6-phosphate carbohydrate
activation lymphocyte cell signalling
positive programmed cell death
biosynthetic heme metabolic pigment
mitotic checkpoint signaling
immune adaptive cell migration
TORC1 signaling positive
transport electron mitochondrial
RNA polymerase II transcription
translation mitochondrial biosynthetic
cilium organization assembly
spindle organization mitotic
GPI anchor metabolic biosynthetic
interstrand cross-link repair
endoplasmic reticulum stress
Golgi organization endomembrane
organelle disassembly reticulophagy
biosynthetic phospholipid metabolic
DNA replication negative recombination
reciprocal recombination meiotic
peroxisome protein organization
transport endocytic recycling
B
1.00
0.75
0.50
0.25
0
yeast mean correlation quantile
0-0.2
0.2-0.4
0.4-0.6
0.6-0.8
0.8-1.0
human mean correlation quantile
C
1.00
0.75
0.50
0.25
0
yeast mean correlation quantile
0-0.2
0.2-0.4
0.4-0.6
0.6-0.8
0.8-1.0
human mean correlation quantile

Figure S13
